# Modelling mouse barrel cortex: from anatomical structure to cortical dynamics

**DOI:** 10.64898/2026.11.24.768894

**Authors:** Aleksandra Teska, Tâm J Nguyên, Alexis Arnaudon, Lida Kanari, Hugo Dictus, Elvis Boci, James Isbister, Felix Schürmann, Sylvain Crochet, Carl Petersen

## Abstract

Detailed biophysical simulations of neuronal networks might help understand brain function. Here, we built a model of mouse barrel cortex focused on its somatotopic map, comprising 33 barrel columns. We investigated dendritic polarity in layer 4 and implemented it in the model, building two circuits differing in a single property of basal dendritic synthesis. In the polarised circuit, layer 4 dendrites were confined to their home barrel; in the symmetric circuit, they were not. We simulated three conditions: i) quiet periods without whisker movement; ii) active free-whisking periods without touch; and iii) active whisking with touch of the C2 whisker. The model exhibited complex spatiotemporal dynamics of neuronal activity across cortical layers, cortical columns, molecularly-defined classes of neurons, and brain states. Using cyclicity analysis, spiking responses were decomposed into activity shared between the two circuits and activity unique to either. Layer 4 dendritic polarisation sharpened the touch response in space and advanced it in time.

## Introduction

Mice use their whiskers to sense their immediate surroundings, often through exploratory whisking and active touch. The neuronal circuitry underlying early steps of whisker sensorimotor processing of tactile information has begun to be studied. Through the somatotopic lemniscal pathway, tactile signals travel from the whisker follicle through the trigeminal ganglion, the principal trigeminal nucleus in the brainstem, the ventral posterior medial nucleus of the thalamus, and subsequently reach the barrel field of the primary somatosensory cortex (Figure 1A) (recently reviewed by: Bosman et al., 2011; Diamond et al., 2008; Petersen, 2019; Staiger and Petersen, 2021). The somatotopic map between whiskers and barrels is discrete and one-to-one: each whisker corresponds to a single ‘barrel’, visible in layer 4 (Van Der Loos & Woolsey, 1973; Woolsey & Van Der Loos, 1970). Each cortical column therefore has a known principal whisker input, so a functional response can be studied against a fixed anatomical structure. For this reason, mouse barrel cortex has been a natural target for many modern recording modalities providing new insight into cortical function, from intracellular and voltage-imaging measurements of membrane potential (Ferezou et al., 2006; Poulet & Petersen, 2008) to multisite extracellular electrophysiological recordings and calcium imaging of population spiking activity (Chang et al., 2024; Chen et al., 2013; Isett et al., 2018; Peron et al., 2015) along with increasingly large-scale reconstruction of cortical wiring with synaptic-resolution through volume electron microscopy (Motta et al., 2019). In principle, a biophysically detailed in silico model of mouse barrel cortex could help integrate across these levels to further mechanistic understanding, thus motivating the current work.

**Figure 1.**
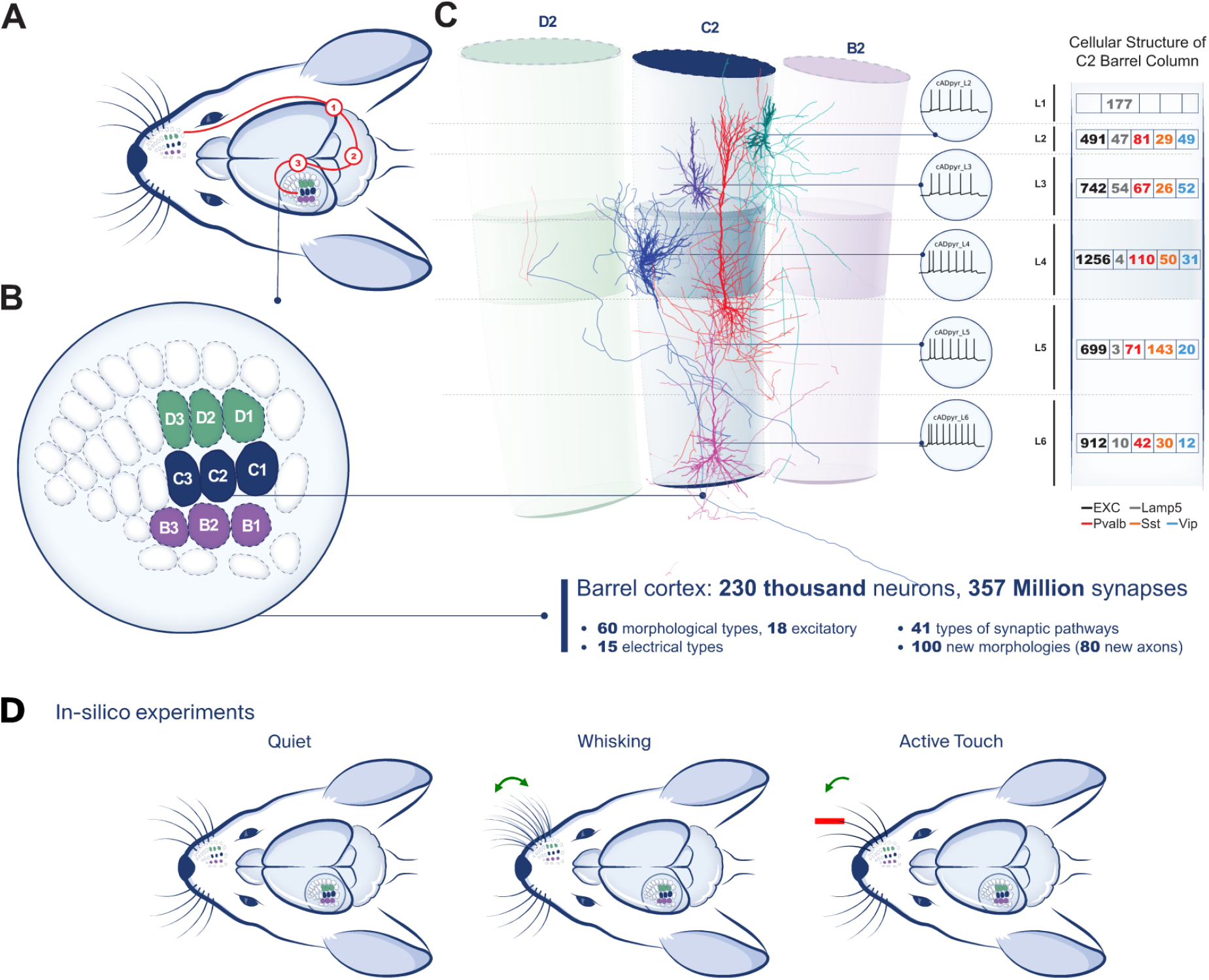
Atlas-based biophysical model of mouse primary somatosensory barrel cortex. **A.** Whisker pathway activity is relayed through trigeminal ganglion primary sensory neurons (1), brainstem neurons (2) and thalamic neurons (3) before reaching the barrel cortex. We present a cortical model with thalamic input modelled explicitly as VPM and POm fibre projections. **B.** Flatmap view of the modelled barrel field, comprising 33 barrel columns registered to the CCFv3 atlas. The highlighted 3×3 grid of barrel columns (rows B, C, D × arcs 1, 2, 3) and the associated septum, centred on C2, is the volume used for experiments and analysis. **C.** *Left:* the modelled D2, C2 and B2 barrel columns, shown as their reconstructed atlas volumes, with example excitatory morphologies from C2 and their corresponding electrophysiological behaviour (middle). *Right:* cellular structure of the C2 barrel column split by molecular class — EXC (black), Lamp5 (grey), Pvalb (red), Sst (orange), Vip (blue); the column comprises 5,200 neurons. **D.** The three in silico experiments performed in this study: (1) quiet, (2) whisking states, and (3) active touch of the C2 whisker.

The most obvious specialisation of the barrel cortex is the one-to-one mapping of the whiskers onto barrels in layer 4 of the primary somatosensory cortex, formed by whisker-specific patches of thalamocortical axons. At a cellular level in layer 4, it is thought that some dendritic trees of layer 4 neurons specifically grow towards and within the barrel that carries their principal whisker input. The bias is instructed during development: it is established by thalamic drive in early postnatal life, redirected under sensory deprivation (Harris & Woolsey, 1981; Jeanmonod et al., 1977) and lost when thalamocortical transmission is genetically disrupted (Abdel-Majid et al., 1998; Li et al., 2013; Narboux-Nême et al., 2012). The clearest case is the layer-4 spiny stellate cell, which arises from a pyramidal precursor by retracting its apical dendrite and polarising its basal tree towards the barrel centre, a transformation that depends on thalamocortical input (Nakazawa et al., 2018). Polarity has been documented in layer-4 spiny stellate and untufted pyramidal cells (Staiger, 2004), non-Martinotti somatostatin cells (Scala et al., 2019) and, in rat, parvalbumin cells (Koelbl et al., 2015). These reports establish that polarity exists, but each describes it qualitatively and in one cell type at a time; it has not been measured with a common descriptor across morphological types, so how far it extends beyond these layer-4 cases is unknown. Its contribution to function remains untested, as dendritic polarity cannot be removed without also disrupting the thalamocortical input that instructs it.

Bottom-up modelling allows perturbations to be introduced independently: dendritic polarity can be imposed or removed while the thalamocortical wiring is maintained. Detailed circuits are reconstructed from cell densities, morphologies and predicted connectivity, most comprehensively for rat non-barrel somatosensory cortex (Isbister et al., 2024; Reimann et al., 2024). The rat barrel field has been reconstructed (Egger et al., 2020; Narayanan et al., 2017; Udvary et al., 2022) and used to study layer 5 pyramidal tract responses (Bast et al., 2024). In mouse, barrel cortex has been simulated with simplified neurons (Huang et al., 2022), and a detailed large-scale model exists for visual cortex (Billeh et al., 2020). None yet exists for mouse barrel cortex at this level of detail.

Mouse datasets for such a model continue to accumulate, increasingly for molecularly defined cell classes (Pala & Petersen, 2018; Tremblay et al., 2016; Yu et al., 2019). They can be registered to the Allen Common Coordinate Framework (CCFv3), in which a bottom-up model can be built directly (Kim et al., 2017; Wang et al., 2020). We adapt the rat workflow (Isbister et al., 2024; Reimann et al., 2024), narrowing it from a multi-region volume to a single region grounded in the geometry of individual barrel columns. Morphologies are synthesised to match reconstructed cells while conforming to local geometry, so dendritic polarity becomes a parameter that can be applied to selected morphological classes or removed.

We quantified dendritic polarity across curated mouse barrel cortex reconstructions with a single descriptor set, and used the result to choose which classes to constrain at synthesis. We built two circuits differing only in this constraint, polarised and symmetric; in an animal this comparison is unavailable, as preventing barrel formation also perturbs the input pathway. In both circuits we simulated three conditions (Figure 1D): i) quiet periods, ii) active whisking without touch, and iii) active whisking with touch of the C2 whisker. We ask how well the model reproduces reported mouse dynamics, which cells carry the touch response, and what changes when the dendritic constraint is removed.

## Results

### Biophysical model

We built a model of mouse primary somatosensory barrel cortex (SSp-bfd) in the Allen Mouse Brain Common Coordinate Framework (CCFv3) atlas space (Wang et al., 2020), enhanced with data-driven barrel column annotations and a flatmap of the region (Bolaños-Puchet, Teska, et al., 2024). This volume comprises 33 barrel columns and approximately 230,000 neurons, connected by around 357 million synapses. Neurons are drawn from 60 morphological types and 15 electrical types, paired to define the morpho-electrical composition of each layer. Experimental work identifies inhibitory cells by molecular class, so we additionally assign every inhibitory neuron one of four molecular classes: parvalbumin-positive (Pvalb), somatostatin-positive (Sst), vasoactive intestinal peptide-expressing (Vip) or Lamp5-expressing (Lamp5), sampled from a probabilistic mapping of its morpho-electrical type (Section *Molecular identity*). Each neuron carries a synthesised dendritic tree, generated to match the statistical properties of reconstructed cells while conforming to the local geometry of the region. Connectivity is predicted from the anatomical overlap of axons and dendrites.

### Atlas-based neuron placement

CCFv3 is a voxelised reference atlas at 25 µm resolution in which each voxel carries a Cartesian coordinate, a region label, and a layer label. In previous work we enhanced this atlas for the barrel field, annotating 33 barrel columns, constructing depth measurements, and a flatmap of the region (Bolaños-Puchet, Teska, et al., 2024). The columns are non-overlapping, separated by septa. Each column is subdivided with layers, so that barrel, septum, and layer identity are defined jointly for every voxel of the field. The flatmap is a two-dimensional representation of the horizontal extent of the layers, obtained by collapsing the depth dimension through a structured, area-preserving projection. A point in the flatmap corresponds to a curved, layer-spanning column of tissue in 3D, and conversely any such column maps back to a region of the flatmap. Neurons are placed voxel by voxel from the density recipe (Section *Densities*). Each soma inherits the orientation vector of its voxel, which points toward the cortical surface and sets the axis along which its morphology is aligned, so that cells follow the curvature of the cortex everywhere in the volume. The flatmap gives the horizontal coordinates used for spatial analysis throughout.

#### Densities

Cell densities were derived following the same approach as the rat model, using layer-specific density profiles from (Keller et al., 2018) extended to mouse. On this baseline, we applied refinements where mouse-specific data indicated necessary differences, concentrated in the interneuron populations of the granular and infragranular layers. In layer 4, we refined the excitatory composition so that spiny stellate cells (SSC) make up 76% of excitatory neurons, and set layer 4 Sst neurons to 96% non-Martinotti (Scala et al., 2019). We further introduced a barrel-septum distinction absent from the uniform baseline: the densities of layer 4 Vip and Sst neurons differ between barrel and septum compartments (Almási et al., 2019), a refinement made possible by the explicit barrel-column geometry of our atlas. Finally, we adjusted the depth profiles of GABAergic neurons, refining the density of layer 5 Sst neurons (Tremblay et al., 2016). One feature of the reference data was not reproduced: our profiles do not capture the pronounced peak of Pvalb neurons in layer 4 (Tremblay et al., 2016).

### Molecular identity

The model defines neurons by their morpho-electrical (me-types) properties, whereas experimental work typically identifies inhibitory cells by their molecular identity: Pvalb, Sst, Vip or Lamp5. Structural, physiological and in silico analyses throughout this study group cells by molecular class, so we assigned an identity to every inhibitory neuron. Excitatory neurons form a single class, subdivided into layers, that is already carried by the morphological type. We used the probabilistic mapping of Roussel et al. (2023), who co-embedded the rat me-types and Cre-driver line expression in a shared latent space of morphological and electrophysiological features. Cluster composition gives, for each me-type, a distribution over the four markers. Every inhibitory neuron drew its marker stochastically from the distribution matching its me-type, so a single me-type contributes cells to several molecular classes. A molecular class pools morphological types that differ in laminar axon targeting, permitted postsynaptic compartment and spine reach. Pvalb pools the basket and chandelier types, the only inhibitory types to contact pyramidal somata; Sst the Martinotti types; Vip the bipolar and double bouquet types; Lamp5 the neurogliaform types and the layer 1 families. No constraint in the model acts on molecular class: every placement, touch and pruning rule is defined on morphological and electrical types, and the mapping is applied afterwards. Structure in the molecular-class matrices reported throughout is therefore an output of the mapping, not a target of the recipe.

### Morphological reconstructions

The model’s dendritic morphologies draw on two sources: the reconstruction library of the rat somatosensory cortex model (Reimann et al., 2024), adapted here to mouse, and a set of mouse barrel cortex reconstructions curated for this work.

#### Mouse barrel cortex morphologies

We assembled reconstructions from public repositories (Ascoli et al., 2007; MouseLight Project Team, 2019) and published datasets (Scala et al., 2019), differing in laboratory, protocol and animal age. An initial set of 324 dendritic reconstructions passed quality control with NeuroR (Anwar et al., 2009) and was mapped onto a consolidated morphological-class scheme. A stricter final curation retained 100 morphologies, 80 of them with reconstructed axons, which we rotated, repaired and annotated. These curated reconstructions provided the basis for the dendritic polarity analysis. Notable populations include layer 4 spiny stellate, pyramidal cells, layer 2/3 bipolar and Martinotti cells (Figure 2 A).

**Figure 2.**
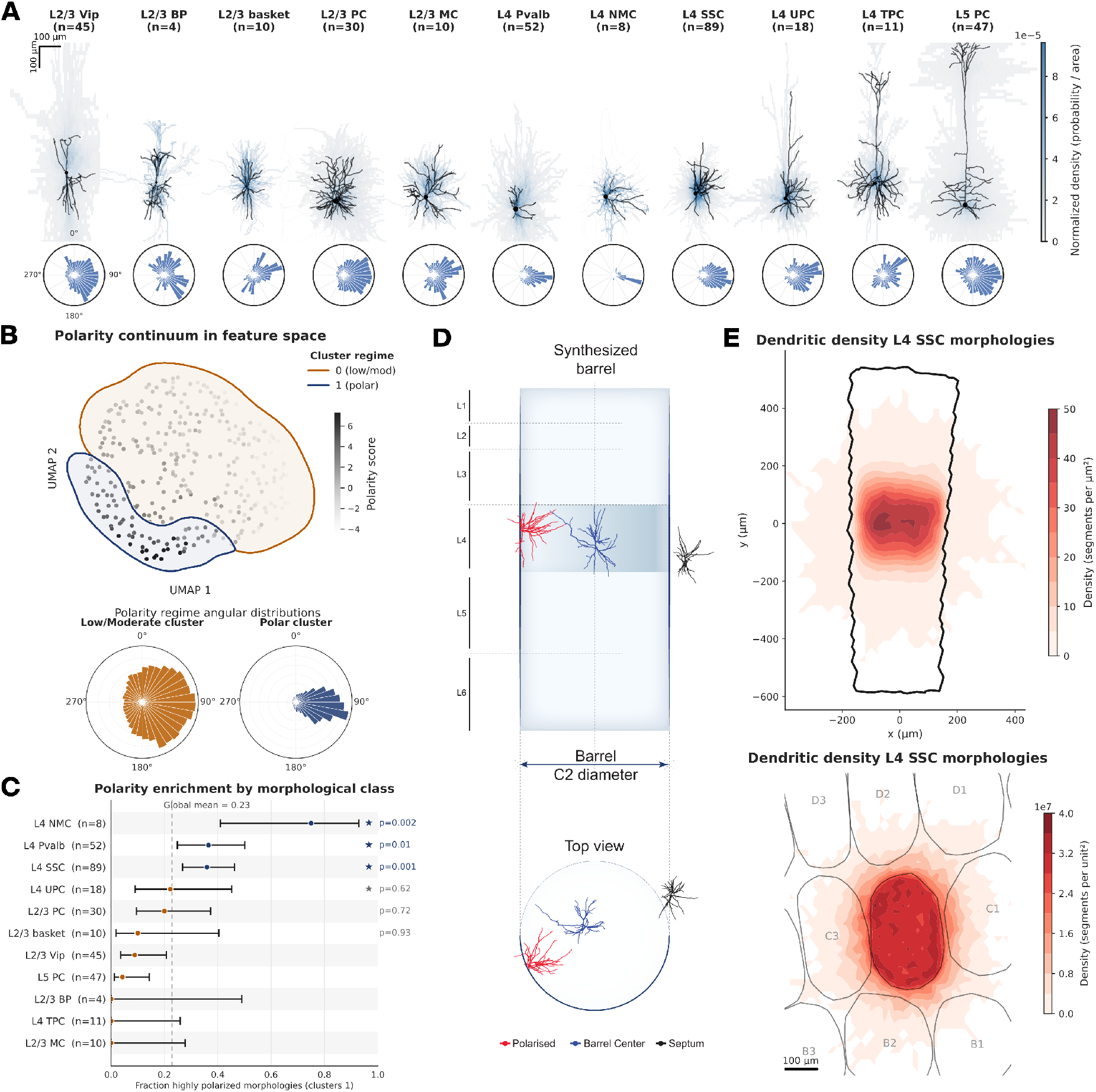
Mouse barrel cortex morphologies, dendritic polarity, and boundary-guided synthesis. **A.** Curated mouse barrel cortex reconstructions grouped by morphological class (number of cells in brackets). Top: individual example morphology (black) over the normalised dendritic density of the full class sample. Bottom: length-weighted angular distribution of dendritic mass in the flatmap plane - the polar histogram - pooled per class after rotating each cell to a common mean direction. **B.** Polarity in feature space. Five geometric descriptors were extracted per cell from single and two-component von Mises fits to its polar histogram, and cells were embedded and clustered on these five features, separating a polar from a low/moderate regime. *Top:* UMAP embedding (McInnes et al., 2018), shaded by the composite polarity score, outlined by cluster. *Bottom:* mean polar histogram per cluster - a single dominant direction in the polar cluster, close to uniform in the low/moderate one. Clustering provides an objective assignment of cells to the two regimes. **C.** Fraction of cells in the polar cluster per morphological class, with Wilson 95% confidence intervals and the number of reconstructions per class, against the global polar fraction (0.23, dashed). Enrichment was tested per class against all other classes pooled, one-sided Fisher exact test on polar and non-polar counts; stars mark *p* < 0.05, and all three classes remain significant after Benjamini-Hochberg correction (Methods). L4 SSC, L4 NMC and L4 Pvalb are significantly enriched. L4 UPC was constrained in the model on the basis of prior literature reports. **D.** Boundary-guided dendritic growth. Atlas-derived meshes define the boundary of each cell’s home barrel column, and synthesis is biased against crossing it, keeping the constrained L4 classes within their volumetric limits. Side and top view of the synthesised C2 barrel showing a constrained cell at the barrel edge (red), a cell at the barrel centre where the constraint has little effect (blue), and an unconstrained septal cell (black). **E.** Dendritic density of synthesised L4 SSC morphologies with soma in the C2 barrel. *Top*: side view within the C2 outline. *Bottom*: view across the barrel grid.

#### Adapting the rat somatosensory reconstructions

Detailed dendritic reconstructions of mouse somatosensory cortex remain sparse. The large-scale mouse datasets (Gouwens et al., 2020; Scala et al., 2021) and whole-brain efforts (Peng et al., 2021; Winnubst et al., 2019) sample a variety of cortical areas, and do not cover the barrel field evenly across morphological types. The rat somatosensory reconstructions (Kanari et al., 2019; Markram et al., 2015) span all 60 morphological types, so we adapted them to mouse anatomy (“mousification”). Rat reconstructions were first curated with established morphology workflows (Anwar et al., 2009) to correct reconstruction gaps and diameter artefacts, then compared against public mouse reconstructions (Ascoli et al., 2007; Allen Institute) to derive species-specific transformations: because cortical thickness is reduced in mouse (DeFelipe et al., 2013), morphologies were uniformly scaled by 0.66 of their original size, and axonal and dendritic diameters - including tapering profiles - were rescaled to the mouse distributions. Transformations were fitted separately for pyramidal cells and interneurons, which differed substantially. The resulting mousified morphologies preserve the cell-type diversity of the rat reconstructions while matching key mouse morphometric features (morphological types examples in Supplementary Figure S2 A).

### Dendritic polarity across morphological classes

Which morphological classes are polarised, and whether the property extends beyond the known layer 4 cases, has not been measured. We quantified dendritic polarity across morphological classes, and used the result to decide which classes to constrain in the model. The analysis used mouse barrel cortex reconstructions only (Section *Morphological reconstructions*). For every reconstruction, we measured the orientation of each basal dendritic segment in the flatmap plane, relative to the soma, and weighted it by segment length, giving a length-weighted angular distribution of dendritic mass around the cell (Fig 2A). Because neurons were reconstructed at different positions and orientations within the barrel, each distribution was rotated to a common mean direction, so that polarity could be compared across the population (Supplementary Figure S2 B).

From each aligned histogram, we derived five geometric descriptors. A single von Mises fit gave the overall concentration *κ* describing how tightly the histogram peaks around a single direction. A two-component von Mises mixture then gave the concentration of the dominant component *κ*_primary_, its weight *w*_1_, and the angular separation Δ*μ* between the two component means. A fifth descriptor, the effective resultant length *R*_eff_, measures how directional the histogram is overall: it runs from 0 for a flat, uniform histogram to 1 for one concentrated on a single direction (Methods, Supplementary Figure S2 C). Together these descriptors distinguish single-axis polarisation (one sharp, dominant mode) from symmetric bipolarity (two balanced modes, roughly opposed), which we do not count as polar. Dendritic polarity is best understood as a continuum, not a discrete label.

We captured it with a continuous composite score that rises with directional coherence and falls with symmetric bimodality (Figure 2B):

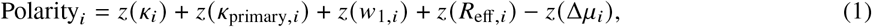

where *z* (*x_i_*) = (*x_i_* - *x̄*) /*σ_x_* denotes the z-score of each descriptor across all cells, with *x̄* and *σ_x_* the population mean and standard deviation of that descriptor. Standardizing puts the five descriptors on a common scale, and Δ*μ* enters with a negative sign because a large angular separation between the two mixture means indicates symmetric bimodality.

To decide which classes are enriched and which to constrain in the model, we grouped cells into *polar* and *low/moderate* regimes by clustering the five features (Methods). This split is a practical cut through a continuous axis rather than evidence of two separate populations, and we treat it as such throughout (Figure 2B-C, Supplementary Figure S2F). Polarity was strongly cell-type dependent (global polar fraction 0.23). Three layer-4 classes were significantly enriched above the global mean: L4 SSC (*p* = 0.001), L4 NMC (*p* = 0.002) and L4 Pvalb (*p* = 0.01) (Figure 2C). We additionally constrained L4 UPC in the model: it was not enriched in our sample (*p* = 0.62), but is reported as polarised in prior work (Staiger, 2004). The remaining classes, including the L2/3 and L5 pyramidal types, showed no enrichment and were left unconstrained. This same L4 enrichment held when cells were ranked directly by the continuous score rather than by cluster membership, confirming it does not hinge on the cluster boundary. These class-level results should be treated as indicative and preliminary, given the limited morphology availability and the lack of a dedicated dataset.

### Morphological synthesis

The curated morphology dataset provided the input for topology-guided synthesis. Within each morphological class, dendritic structure was captured by a topological morphology descriptor (a persistence barcode) that describes how the tree’s branches are organised across successive distances from the soma. We used these barcodes to grow new trees statistically indistinguishable from the reconstructions in that class (Kanari et al., 2022). Beyond diversity, synthesis opens an opportunity that reconstructions alone cannot. Because each tree grows step by step rather than being copied whole, we can control the shape it takes as it grows. In particular, growth can be constrained by the anatomical boundaries of the barrel column, so that dendrites develop within their volumetric limits.

#### Boundary-guided dendritic growth

For each cell, atlas-derived meshes define the boundary of its home barrel column. During synthesis, we bias the random walk against crossing this boundary: each candidate growth step is accepted with a probability that rises with its distance from the mesh wall (Methods, Section *Morphologies*). Deep inside the column, steps are accepted freely, so growth follows the unconstrained algorithm. Near the wall, outward steps are increasingly likely to be rejected, and the branch tends to grow back into the column. The bias acts on basal dendrites only, so apical dendrites and axons grow unconstrained. Septal neurons of these classes lie outside the barrel meshes and were left unconstrained. Applied to the constrained layer-4 classes, this keeps the dendritic trees largely within the home barrel, reproducing the polarised architecture. In C2 barrel column, L4 SSC cells keep 89.2 12.1% of their basal dendritic length inside the barrel outline, against 64.7 22.5% without the constraint (mean SD over 951 cells). The other constrained classes follow the same pattern: 88.9 13.5% against 67.9 25.7% for L4 UPC, 89.8 10.5% against 68.5 26.1% for L4 NMC, and 75.7 11.3% against 59.6 33.1% for L4 Pvalb. Cells vary in how much dendrite they keep inside the barrel, and the biggest difference between the circuits is in the tail: the 10th percentile of L4 SSC cells retains 74.0% of their dendrite under the constraint against 32.1% without. Total basal dendritic length per cell is the same in both circuits (3350 ± 1500 against 3320 ± 1460 µm for L4 SSC), so the constraint redirects growth without changing the size of the tree.

#### Barrel adaptation

The boundary constraint defines two model variants. In the polarised circuit, the constrained layer 4 classes grow under the repulsion rule, each neuron polarised towards the centre of the barrel its soma occupies; since the constraint is defined in atlas space, one rule orients every column correctly across the barrel field. In the symmetric circuit, the same classes are synthesised with the constraint removed. Everything downstream of synthesis was built for each circuit against the same targets. Connectivity is predicted from the morphologies each circuit carries, postsynaptic potential amplitudes are drawn from the same recipe, and e-models are paired to each morphology by the procedure of Section *Electrophysiological models*. The layer 4 dendritic constraint is the only design choice that differs; everything else was regenerated the same way in both.

#### Axons and appositions

Axons used in the model are reconstructions, not synthesised trees. Each neuron received a reconstructed axon of its morphological type, scaled and rotated on the vertical axis before attachment to the dendritic tree. These axons form the input to the connectome. Placement is guided by laminar targeting rules defined per morphological type, applied to axons and to apical dendrites (Section *Laminar placement rules*, Table 1). These rules are inherited unchanged from the rat somatosensory cortex model (Reimann et al., 2024), so laminar structure is imposed on the morphologies, and imposed from rat, before any synapse is placed. The rules constrain depth only. No rule in the recipe references a barrel, a column or a flatmap coordinate. Horizontal structure in the connectome follows from atlas geometry and from the extent of the morphologies alone.

**Table 1.** Laminar targeting rules per morphological type. Rules are inherited unchanged from the rat somatosensory cortex model (Reimann et al., 2024) and constrain depth only. Each interval runs from the lower boundary of the first layer to the upper boundary of the second, unless a fraction is given; a fraction is the position within that layer, from 0 at its lower boundary to 1 at its upper boundary. Types listed on one row share the rule, and types with two targets carry two independent rules. Two global rules apply to all types: dendrites remain below the pia, and axons exceed it by at most 40 *μ*m. Types not listed have no laminar rule.

| Segment | M-types | Target interval |
|---|---|---|
| Axon | L1_HAC, L23_MC, L4_MC, L5_TPC (:A, :B, :C), L5_UPC | L1 |
| Axon | L5_MC | L1; L4 |
| Axon | L6_MC | L2 → L1; L4 |
| Axon | L4_SSC | L2 → L1 |
| Axon | L2_TPC (:A, :B), L3_TPC (:A, :C) | L5; L4 |
| Dendrite | L2_TPC (:A, :B), L3_TPC (:A, :C), L5_TPC (:A, :B, :C) | L1 |
| Dendrite | L4_TPC, L4_UPC | L2 → L1 |
| Dendrite | L6_TPC (:A, :C), L6_UPC | 0.8 L5 → 0.5 L3 |
| Dendrite | L6_BPC | 0.5 L5 → 0.1 L3 |

### Electrophysiological models

The model uses 15 electrical types (e-types): five continuous-accommodating pyramidal types, one per excitatory layer (cADpyr L2 to L6), and ten inhibitory firing types - burst-accommodating (bAC), burst non-accommodating (bNAC), continuous accommodating (cACint), continuous non-accommodating (cNAC), delayed non-accommodating (dNAC), burst irregular firing (bIR), continuous irregular firing (cIR), burst stuttering (bSTUT), continuous stuttering (cSTUT) and delayed stuttering (dSTUT). The response of each e-type to depolarising and hyperpolarising somatic step current injection can be seen in Supplementary Figure S1G. The e-models were fitted to whole-cell patch-clamp recordings of rat somatosensory cortex with the BluePyEModel workflow (Reva et al., 2023) and taken here unchanged.

Each e-model is optimised on a specific morphological reconstruction, as its firing behaviour depends on the balance of input resistances between axon initial segment, soma and dendrites. To assign e-models to unique synthesised morphologies we used the sampling and generalisation workflow of Arnaudon et al. (2023). The workflow has two parts: sampling many variants of each e-model, and adapting each morphology to the variant it is paired with. Markov chain Monte Carlo sampling returns many parameter sets per e-type, all matching the target electrophysiological features; cells sharing an e-type therefore differ in their channel densities. The soma and axon initial segment of each morphology are then rescaled until its input-resistance ratios match the target for that e-type. A pairing was used in the circuit only if every electrophysiological feature of the resulting cell fell within five standard deviations of the experimental mean. The resulting e-type composition per morphological class is given in Supplementary Figure S1C.

### Connectome

We built the connectome with the apposition-based algorithm of Reimann et al. (2015), where appositions between axons and dendrites give potential synapses, which are then pruned to a biologically viable set of connections. The algorithm predicts the mean number of synapses per connection from the mean number of appositions per connection, and the multi-synaptic pruning step follows from that prediction. The rules under which appositions are detected and pruned come from a recipe inherited from the rat somatosensory cortex model (Reimann et al., 2024). It fixes the bouton density along each axon, the minimum distance from the soma at which a synapse may form, and the postsynaptic compartments each morphological type may contact. Both distance and compartment rules differ between excitatory and inhibitory cells (Methods, Section *Connectivity recipe*).

Reimann et al. (2015) used a square-root relation for most pathways, which compresses the range of apposition counts and drives the distribution towards a log-normal centred on several synapses per connection. We used a linear relation for all pathways, which preserves the shape of the underlying apposition distribution and preserves single synapse connections (Methods, Section *Connectivity recipe*). Recent dense electron-microscopic reconstruction of mouse cortex reports such connections at a higher rate than the rat paired recordings the original relation was tuned against (Schneider-Mizell et al., 2025). The recipe specifies no connection probability and no pathway-specific connection count, and no rule in it references a barrel, a column or a horizontal coordinate.

#### Cortical synaptic physiology

No complete synaptic dataset exists for mouse barrel cortex. We took synapse kinetics and short-term dynamics from the rat paired-recording models of Markram et al. (2015), which cover five main synapse families: two excitatory, facilitating (E1) and depressing (E2), and three inhibitory, facilitating (I1), depressing (I2) and pseudo-linear (I3). The model carries 41 synapse types, each falling within one of these families (Figure 3A, B). To give the model mouse synaptic strengths, we curated postsynaptic potential amplitudes per pathway from the mouse barrel cortex literature (Supplementary Figure S3B) and added the Allen Institute mouse V1 dataset (Supplementary Figure S3A, Campagnola et al., 2022). We extrapolated across layers and cell classes to cover the pathways neither source reports, giving a complete amplitude recipe (Supplementary Figure S3C).

**Figure 3.**
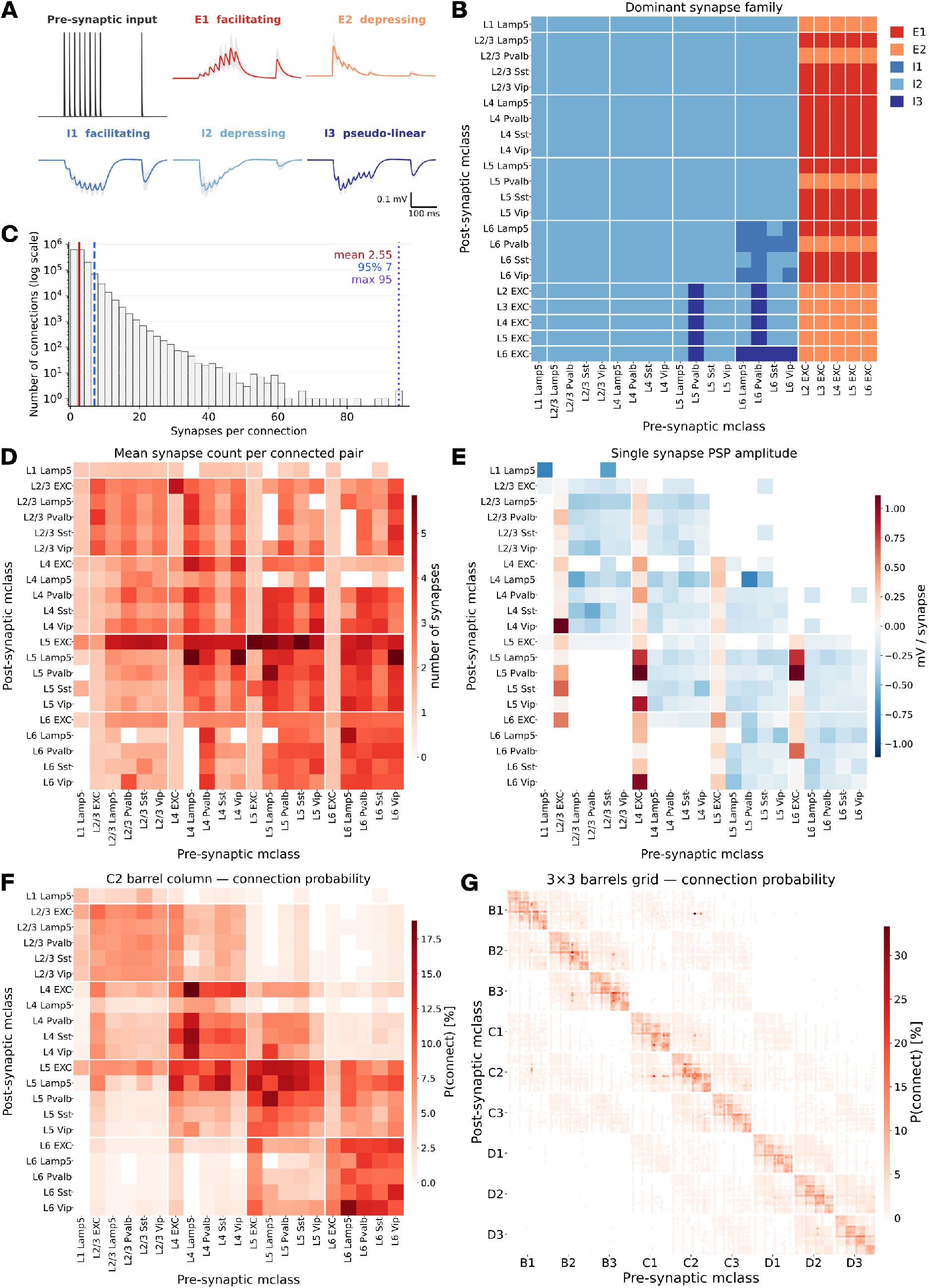
Connectome structure. **A.** The main synapse families in the model. Postsynaptic potential in response to a presynaptic spike train (black), for the two excitatory families - E1 facilitating, E2 depressing - and the three inhibitory families - I1 facilitating, I2 depressing, I3 pseudo-linear. Mean across sampled synaptic connections and trials; shading shows SD. **B.** Dominant synapse family for each pre- to post-synaptic molecular class pathway. **C.** Distribution of synapses per connected pair across the model. Mean 2.6 2.3, median 2, 95th percentile 7, maximum 95. **D.** Mean number of synapses per connected pair, per molecular pathway. **E.** Single-synapse postsynaptic potential amplitude, per molecular pathway. Positive (red) excitatory, negative (blue) inhibitory. **F.** Connection probability between molecular classes within the C2 barrel column. **G.** Connection probability across the 3×3 barrel grid, molecular classes within each barrel block.

#### Connectome structure

Most connected pairs share one or two synapses, which together account for 65.9% of connections in the C2 barrel column. Across the model, the count per pair decays monotonically into a long tail reaching 95, with a mean of 2.6 2.3 (median 2, interquartile range 1-3; Figure 3C). Pathway means run from 1.4 synapses per connection (Lamp5 to Lamp5) to 3.5 (Sst to excitatory), with the highest counts in inhibitory pathways onto excitatory cells (Figure 3D). Inhibitory connections average 3.1 2.6 synapses against 2.4 2.2 for excitatory ones, and 43.9% of excitatory connections are formed by a single synapse against 20.9% of inhibitory ones. This distribution is closer to recent dense mouse reconstructions than to the rat paired recordings, which reported fewer single synapse connections. Connection probability within the C2 barrel column is organised into within-layer blocks, with within-layer pathways three times more likely to be connected than across-layer ones (9.6% against 3.2%). Within-layer connection probability is higher in layers 5, 6 and 4 (11.3%, 10.5% and 9.9%, respectively) than in layer 2/3 (6.9%) (Figure 3F).

The excitatory neurons form the translaminar pathways, following the laminar axon targeting rules of the recipe. L4 EXC projects upward onto L2/3 and downward onto L5. L5 EXC is the broadest postsynaptic target in the column, receiving from every layer and most strongly from layer 4. L6 EXC connects with layer 5 and more weakly with layer 4. Inhibitory classes connect predominantly within their own layer. Across the 3 3 barrel grid, connectivity is concentrated in the nine diagonal blocks (Figure 3G). Connection probability is not algorithmically constrained, so the values reported here are an output of the model.

Every neuron in the model carries its own synthesised dendrite rather than a copy of a reconstruction, so neurons of the same class differ in their individual branching geometry while sharing their statistics. This bears differently on the two orders of connectome structure. First-order structure, namely distance-dependent connection probability and reciprocal bias, follows from the average dendritic and axonal extent of each class and is unaffected. Higher-order structure is sensitive to individuality: with a small pool of exemplar morphologies, neurons of a class are copies of one another, and shared partners are inflated by that duplication rather than by circuit structure (Reimann et al., 2017).

We took pairs of layer 4 neurons within 150 µm of each other and grouped them into eight bins by the percentile of their number of common neighbours, neurons connected to both members of the pair. Relative to the mean directed connection probability, the probability that two neurons are connected in either direction rises from the lowest to the highest bin, averaging 1.8 0.9 across bins (range 0.7-3.3) for L4 EXC and 1.8 1.1 (0.4-4.0) for L4 Pvalb (Supplementary Figure S3F). A distance-matched null also rises, more slowly (1.8 0.2, range 1.5-2.1, for L4 EXC; 1.8 0.2, range 1.4-2.0, for L4 Pvalb), and the model curves cross it near the median. A unique tree per neuron avoids the inflation from duplicated morphologies, so the clustered connectivity and common-neighbour bias reported here are properties of the connectome rather than of the morphology pool.

### Thalamocortical nuclei

We modelled two thalamocortical pathways: the lemniscal pathway from the ventral posteromedial nucleus (VPM) and the paralemniscal pathway from the posterior medial nucleus (POm) (Staiger & Petersen, 2021). VPM projections are somatotopic and monovibrissal, each barreloid innervating one barrel column, so we confined VPM fibres to individual barrel outlines; their synapses concentrate in layer 4. POm projections are multivibrissal and carry no barrel-aligned structure, so we distributed POm fibres evenly across the horizontal extent of SSp-bfd, septum included; their synapses concentrate in upper layer 5 and layer 1 (Figure 4A, C).

**Figure 4.**
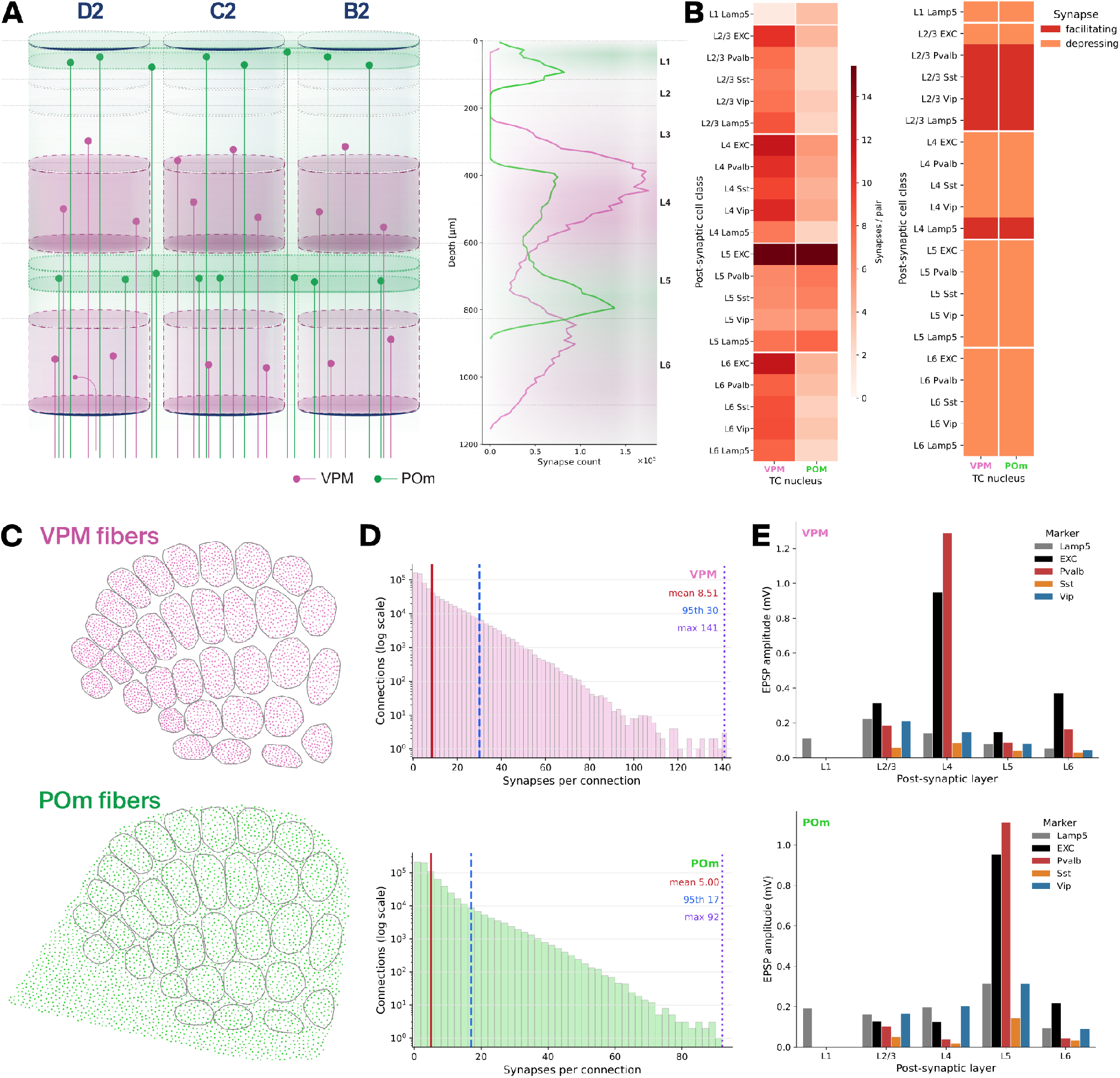
Thalamocortical projections in the model: lemniscal (VPM) and paralemniscal (POm) pathways. **A.** *Left:* schematic of thalamocortical projections onto three neighbouring columns. VPM fibres (magenta) target the barrel columns specifically; POm fibres (green) are distributed evenly across the cortex, including the septum. *Right:* depth profile of thalamocortical synapse density across the C2 barrel column for both nuclei. **B.** *Left:* mean number of synapses per connected pair between each nucleus and each postsynaptic molecular class. *Right:* dominant synapse family per pathway - both nuclei are excitatory, facilitating (red) or depressing (orange). **C.** Flatmap view of thalamocortical fibre positions, showing how fibres were placed. VPM fibres are confined to the barrel outlines; POm fibres are distributed evenly across the field. **D.** Distribution of synapses per connected pair for a single thalamocortical cell. *Top:* VPM (mean 8.5 10.3, maximum 141). *Bottom:* POm (mean 5.0 6.2, maximum 92). **E.** Postsynaptic potential amplitude from a single thalamocortical fibre onto cells of the C2 barrel column, by postsynaptic layer and molecular class. *Top:* VPM. *Bottom:* POm.

#### Digital fibres

Thalamocortical axons are not reconstructed in the model. Each projecting thalamic neuron enters as a digital fibre, following the implementation of the rat somatosensory cortex model (Isbister et al., 2024; Reimann et al., 2024). A digital fibre models a thalamocortical projection as a straight line running through the cortical depth, carrying a profile of synapse density along its length without branches. Complete reconstructions of thalamocortical axons are too few to populate thousands of afferents, so using them directly would require repetition. The abstraction avoids duplication and places the fibre count and the depth distribution of their synapses under direct experimental control.

Three ingredients define a fibre: a cortical depth profile of synapse count, a lateral spread radius, and a position in the cortex, placed in the flatmap. The fibre extends from its flatmap position along the local cortical orientation vector, so that it crosses the full depth of the cortex. Synapse positions along the fibre follow the depth profile. Each synapse is formed on a dendritic segment located within the lateral radius, with a probability that decays with distance from the fibre axis. In simulation, each fibre acts as an individual thalamic neuron, transmitting its action potentials to its thalamocortical synapses after a conduction delay.

#### Number of fibres

Nucleus-wide neuron counts, 68,142 for VPM and 10,286 for POm, follow from reported densities (Keller et al., 2018) and atlas volumes (Methods, Section *Thalamic nucleus size*). Only a fraction of these neurons project to the modelled subfield, so the counts set the scale of each nucleus rather than the number of fibres, and we constrained the fibre count separately for each pathway.

For VPM, we used the whisker-specific relationship between cortex and thalamus reported in rats (Meyer et al., 2013), where the ratio between the neuron counts of a barrel column and its matched barreloid is preserved across the barrel field. We regressed barreloid count on column count across their 24 whiskers (slope 0.011, intercept 61.2, *r*^2^ = 0.68) and applied the fit to each barrel column in our model, giving 4,650 VPM fibres across the 33 columns. The extrapolation assumes the ratio is conserved between species.

POm has no barrel-specific counterpart, so we set its fibre count from the inter-nuclear ratio of 0.76 in the rat somatosensory cortex model (100,000 VPM against 76,000 POm fibres), giving 3,534 POm fibres. Fibre counts per barrel column, and the POm total, are given in Table S1. We placed the fibres on the flatmap using Poisson disc sampling, ensuring a minimum spacing that avoids clustering. We sampled each pathway independently: VPM within each barrel outline, POm across the full barrel field flatmap (Figure 4C).

#### Depth profile and synapse density

The shape of each depth profile is derived from the fluorescence profiles of thalamocortical axonal innervation in mouse somatosensory cortex reported by El-Boustani et al. (2020), each pathway normalised to its own maximum. We took the VPM, which peaks in L4, and the POm higher-order profile for POm, which peaks in the upper half of L5 with a superficial L1 component. Absolute synapse counts are constrained by electron-microscopy measurements of thalamocortical synapse density in mouse L4 (Bopp et al., 2017), to which the L4 peak of each profile was scaled.

#### Lateral spread and conduction delay

Lateral spread was measured from thalamocortical axons reconstructed in mouse and registered to CCFv3 (MouseLight, Braintell). We took the part of each axon lying inside SSp-bfd and kept the reconstructions that were reasonably complete and clearly projected to barrels. We projected the retained segments onto the flatmap and fitted an ellipsoid to each. Its radius defines the fibre spread, 75 *μ*m for VPM and 105 *μ*m for POm. Digital fibres carry the conduction velocity used for cortical axons, 0.3 m/s, which is slower than that of real thalamocortical afferents. Thalamocortical delays in the model are therefore systematically long: the earliest 10% of layer 4 cells in C2 fire their first action potential 8.6 ms after touch onset, instead of the expected 5 ms.

#### Thalamocortical synaptic physiology

Thalamocortical synapses follow the recipe of the rat somatosensory cortex model (Isbister et al., 2024; Reimann et al., 2024), which assigns each pathway to one of two excitatory families, facilitating or depressing (Figure 4B, right). Postsynaptic potential amplitudes differ between the two species, so we recalibrated synaptic conductances against mouse data. Sermet et al. (2019) report amplitudes per postsynaptic layer and molecular class, matching the pathway definitions in the model. The reported amplitudes would not work for model calibration, as the size of the evoked response follows the level of channelrhodopsin expression, which varies between slices. Their amplitudes are normalised within each slice, to layer 4 excitatory cells for VPM and to layer 5A excitatory cells for POm. Using the normalised values allows for representation of the relative strength of thalamic inputs onto molecular classes.

We took the profile as the target, with the reference pathway set to a unitary amplitude of 1 mV, in agreement with Bruno (2006), who reported a mean of 950 *μ*V for monosynaptic VPM connections onto layer 4 barrel neurons in rat. We reproduced the protocol in silico, stimulating a single thalamocortical fibre and recording the postsynaptic potentials in cells of the C2 barrel column. We scaled the peak conductances per pathway until the amplitudes matched the target profile (Figure 4E).

#### Resulting thalamocortical connectome

The two pathways separate along depth, convergence and strength. VPM synapse density peaks in L4 with a second peak at the L5/L6 border, POm in upper L5 with a superficial L1 component (Figure 4A, right). VPM forms more synapses per connected pair than POm, and its distribution has a longer tail (mean 8.5 10.3 against 5.0 6.2, maximum 141 against 92; Figure 4D), and convergence per postsynaptic class follows the same ordering, strongest onto L5 excitatory cells (Figure 4B, left). Single-fibre potential amplitudes follow the same layout, VPM driving L4 hardest and POm L5, with Pvalb above EXC in the principal target layer of each (Figure 4E). These amplitudes were calibrated to the profile of Sermet et al. (2019) and are a target rather than a prediction. VPM and POm are both excitatory and predominantly depressing, facilitating only onto L2/3 interneurons and L4 Lamp5 cells (Figure 4B, right).

### In silico experiments

The model was then used to simulate cortical activity. Simulations were run on a subvolume: the 3 3 grid of barrel columns highlighted in Figure 1B (rows B, C and D arcs 1, 2 and 3), together with the septa and the surrounding tissue, over 130,000 neurons in total. The C2 barrel column, at the centre of the grid, is the focus of the simulations and the analysis, and the surrounding tissue is included so that the response in C2 column is free of edge effects. Analyses of the population response use the nine barrel columns and the septum immediately around C2 column, over 55,000 neurons. Membrane potential and spikes were recorded from every neuron simultaneously, and the afferent connectivity, molecular identity and thalamocortical input of each are known, so structure can be used as a predictor of function. An anatomical feature can be varied in isolation, holding everything else fixed.

Below we compare the model against a second circuit built to maximise similarity: cell placement and the morphologies of all unconstrained classes are identical, and e-models were fitted by the same procedure. The two differ only in the layer 4 dendritic constraint, which isolates what dendritic polarisation contributes to the touch response.

Three experimental contexts were simulated (Figure 1D): (1) the quiet and (2) whisking brain states, and (3) active touch of the C2 whisker, matching the experimental conditions of (Yu et al., 2019) and (Crochet et al., 2011; Kiritani et al., 2023; Yamashita et al., 2013).

### Simulating quiet and whisking states

#### Quiet and whisking inputs

The simulated sub-volume lacks the cortico-cortical and thalamic drive that sets brain state in vivo, so both states were imposed through the input (Figure 5A, B; Methods, Sections *Background conductance* and *Thalamocortical input generation*). Cortical drive was replaced by somatic Ornstein-Uhlenbeck conductance, calibrated to the reported spontaneous firing rates. In the quiet state its level alternates at 5 Hz, producing up and down states. The thalamic input is modulated in phase with it, so cortex and thalamus drive the up state together. In the whisking state it is held at an intermediate level, producing a sustained depolarisation. Thalamic rates follow those reported for the two states (Urbain et al., 2015). VPM and POm both fire faster during whisking, and VPM additionally encodes the 10 Hz whisking cycle, each fibre at its own preferred phase. Whisking was imposed in fixed 600 ms periods alternating with the quiet state, so that the two states are directly comparable within a run.

**Figure 5.**
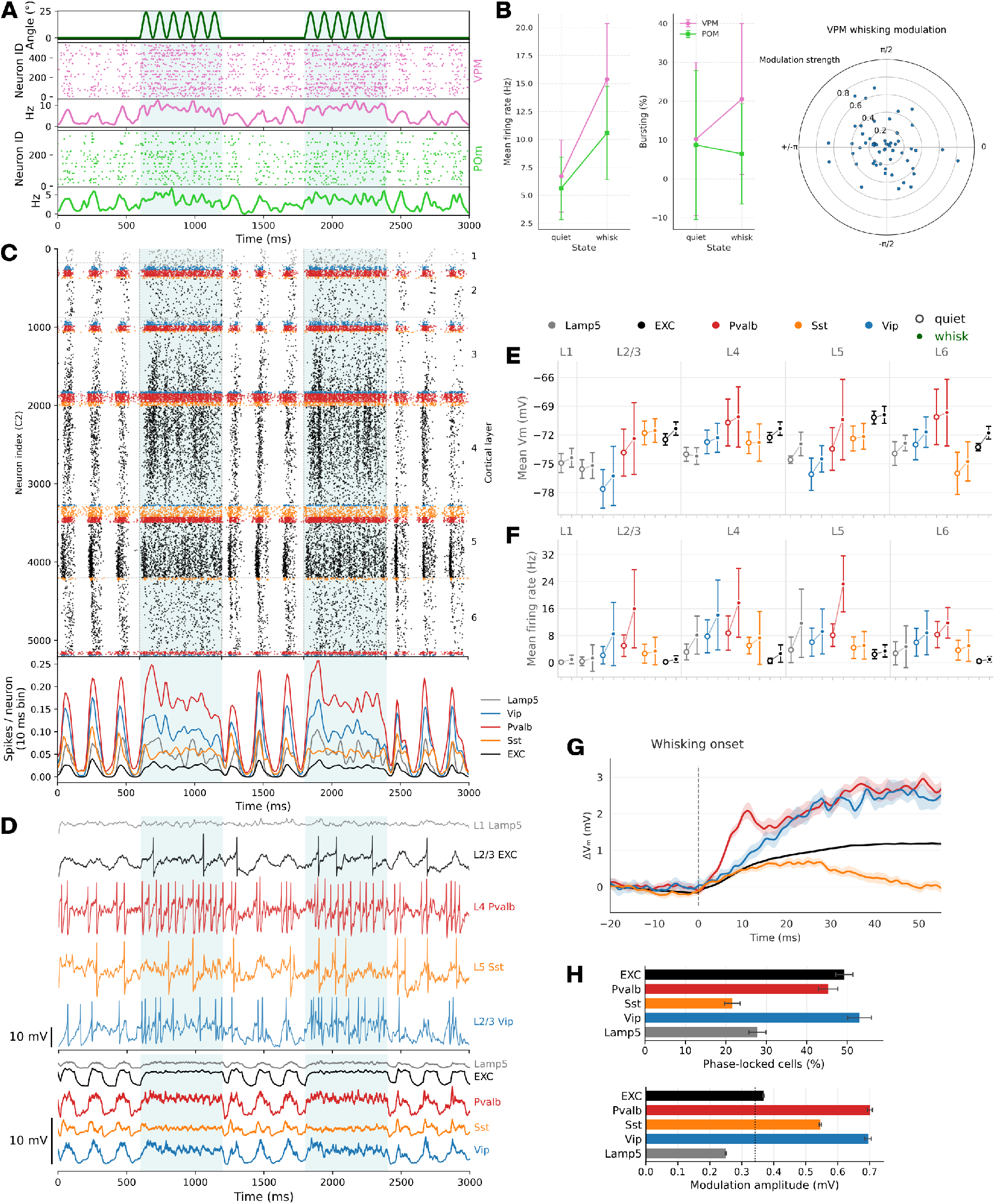
Cortical dynamics during quiet and whisking states. **A.** Simulation inputs. *Top:* whisker angle; shading marks the whisking periods. *Below:* VPM spike raster and population rate (magenta), POm spike raster and population rate (green). One representative simulation, as in C and D. **B.** Thalamic input characterisation. *Left:* thalamic firing statistics per state, from the stochastically generated input: mean firing rate and fraction of spikes in bursts, VPM (magenta) and POm (green); mean SD across fibres. *Right:* whisking modulation imposed on the VPM input: preferred phase and modulation strength of each fibre relative to the whisker angle cycle. **C.** Spike raster of the C2 barrel column, grouped by molecular class and ordered by depth within class - EXC (black), Lamp5 (grey), Pvalb (red), Sst (orange), Vip (blue); colour code used throughout. *Bottom:* population rate per molecular class, 10 ms bins. **D.** Membrane potential in the C2 barrel column, one representative simulation. *Top:* example traces, one cell per molecular class, selected as representative and labelled by class and layer; action potentials removed. Traces are shown as deviation from each cell’s mean. *Bottom:* mean across all cells of each molecular class, same simulation and baseline subtraction. **E.** Mean membrane potential per molecular class, quiet (open) and whisking (filled) states, in the C2 barrel column. Per cell, membrane potential was averaged over seven 600 ms epochs per state and 5 simulation seeds; mean SD across cells. **F.** Mean firing rate per molecular class, same epochs, seeds and averaging as E. **G.** Membrane potential change at whisking onset, per molecular class, from the whisking-onset simulation set. Whisking onset falls at a different point in the up/down cycle in each seed, so no up-state transition is locked to onset, and the preceding up state averages out. Mean SEM across 5 seeds. **H.** Subthreshold phase locking to the whisking cycle, across 25 seeds of a whisking-only simulation set. Whisking phase was taken from the analytic signal of the VPM population rate bandpassed at 7-13 Hz. Null distributions come from circularly shifting the phase clock relative to the membrane potential. *Top:* percentage of cells phase-locked, defined as across-seed phase coherence above the 95th percentile of the null. *Bottom:* modulation amplitude per molecular class, mean SEM across seeds; dotted line marks the median null amplitude.

### Cortical dynamics during quiet and whisking states

#### Activity in C2 barrel column across the two states

Combining the background conductance with the thalamo-cortical input produced sustained cortical activity in both states, and the two are clearly separable in the C2 barrel column (Figure 5C, D). During quiet, activity alternates between up and down states, and the alternation appears at every level of description - in the population raster, in individual membrane potential traces, and in the class means. During whisking, the network settles into continuous activity, with all classes depolarised.

#### Membrane potential, firing and recruitment across the two states

Every molecular class fires faster during whisking, and every class depolarises more when pooled across layers (Figure 5E, F). Quiet epochs were averaged over the full window and include both up and down phases. During quiet, pooled across layers, firing rates are 7.1 ± 4.3 Hz in Pvalb, 4.1 ± 3.1 Hz in Sst, 4.0 ± 4.2 Hz in Vip, 0.8 ± 1.1 Hz in EXC and 0.4 ± 1.3 Hz in Lamp5. During whisking, they reach 17.4 ± 10.5 Hz in Pvalb, 9.7 ± 9.4 Hz in Vip, 5.2 ± 5.2 Hz in Sst, 1.9 ± 2.0 Hz in EXC and 1.4 ± 3.4 Hz in Lamp5. The inhibitory classes fire the most: during whisking, Pvalb, Vip and Sst fire at 9.0, 5.0 and 2.7 times the excitatory rate, while Lamp5 fires below it. Mean membrane potential rises by 0.9 ± 0.7 mV across classes and layers (range −0.2 to 3.1 mV), with the biggest rise in layer 5 Pvalb.

Recruitment, the fraction of cells in a class firing at least one spike over the whole of a state within a run, separates the excitatory and inhibitory classes (Supplementary Figure S4C). During quiet, excitatory recruitment is set by layer, from around 40% in L2/3 to above 80% in L5 and L6, while L4 Sst, L6 Sst, L6 Pvalb and L6 Vip are silent or near-silent. During whisking, excitatory recruitment barely moves, and inhibitory recruitment rises instead, most of it in the deeper layers: L4 Sst, L6 Sst, L6 Pvalb and L6 Vip are recruited to between half and most of the population. Vip cells are the most broadly recruited class during whisking, above 65% in every layer.

#### Whisking onset

The transition into whisking was examined in a dedicated set of simulations in which the phase of the up and down cycle at whisking onset differs between seeds. No up-state transition is therefore locked to onset, and the preceding up state averages out across seeds. Membrane potential and firing rate both change rapidly upon the whisking onset (Figure 5G, Supplementary Figure S4E). Pvalb cells depolarise fastest, reaching 90% of their 4.1 0.2 mV peak by 25 ms (mean SEM across seeds), Vip and EXC more slowly and over a longer rise, at 30 ms and 37 ms. Sst cells depolarise briefly, peaking at 0.97 0.06 mV at 15 ms, before turning to a sustained hyperpolarisation of 0.18 0.10 mV over the second half of the window. Resolved by layer, the fastest response is in L4 Pvalb, which peaks at 10 ms after onset (3.6 0.3 mV) before settling to 1.2 0.2 mV, while L5 Pvalb is the largest, at 6.7 0.3 mV. The firing rate change follows the same pattern, with L4 Pvalb again rising most sharply (Supplementary Figure S4E). The step in background conductance at onset is the same relative step for every cell, and the thalamic drive rises across the whole population, so we looked to the afferent connectivity for the source of the Sst divergence. Sst cells are the only major molecular class receiving more inhibitory than excitatory drive (Supplementary Figure S3E), with a median excitatory-to-inhibitory ratio of 0.10 in layer 2/3 rising to 0.44 in layer 6. Most of that inhibition comes from Pvalb cells, 46-67%, against 4-14% from Vip, and Vip cells do not inhibit Sst cells more than the other interneuron classes do.

#### Subthreshold phase locking to the whisking rhythm

Since the whisking cycle is encoded in the VPM input, we asked whether it is also represented in the membrane potential of cortical cells. Phase was read from the VPM population rate of the C2 barreloid, bandpassed at 7-13 Hz, rather than from the commanded sine, giving a clock that follows the input the cortex actually received, cycle by cycle (Supplementary Figure S6A). Phases are therefore referenced to the peak of the C2 VPM rate rather than to whisker protraction, and are not directly comparable to experimental values. Each cell’s membrane potential was aligned to that clock and a sinusoid fitted to the phase-binned trajectory, giving one modulation amplitude and one preferred phase per cell per seed (Methods, Section *Whisking phase locking*). A cell counts as phase-locked if its preferred phase is consistent across seeds, above a floor set by a time-shifted null.

Close to half of the cells in C2 are phase-locked to the whisking rhythm (46.4%; Figure 5H). Every major molecular class is represented, and modulation amplitude is largest in Pvalb and Vip, at 0.7 0.0 mV each, against 0.4 mV in excitatory cells (mean SEM across 25 seeds; SEM at most 0.01 mV). Sst cells are the least locked, at only 21.6%, against 27.8% in Lamp5 and 45 to 53% in the other classes (EXC 49.3%, Pvalb 45.3%, Vip 53.1%). Preferred phase is class-specific: excitatory cells of layers 4 and 5 concentrate on a preferred phase, the resultant length of their phase distribution reaching 0.9 and 0.8, while those of layers 2/3 and 6 spread more widely, at 0.7 in both (Supplementary Figure S6G).

As a control, we repeated the analysis with the thalamic input present or absent and cortical connectivity intact or removed (Supplementary Figure S6E). With VPM removed, the phase modulation disappears, whether or not cortical connectivity is intact: the population modulation amplitude falls from 0.1 mV to 0.0 mV. With VPM present, it survives disconnection of the cortex, so the connected circuit modulates its amplitude rather than creating it.

### Cortical response to active touch of the C2 whisker

Following the brain state investigation, we next studied the response of the in-silico cortex to active touch during whisking. To compare with equivalent experimental data, we modelled active contacts between an object and a single whisker, C2. Active touch is modelled as a VPM input only, with a log-normal spike density fitted to the VPM population response of Yu et al. (2019). POm carries no touch signal in the model, and receives no additional drive at touch. Whilst the C2 barrel column received the touch input, the other barrels of the grid kept their whisking input (Figure 6A). Each touch recruited half of the C2 whisking fibres, a quarter of all fibres projecting to the C2 column. Touches were delivered in isolation, one every 500 ms, a multiple of the 100 ms whisking cycle, so that every touch arrived at the same phase of the whisking cycle.

**Figure 6.**
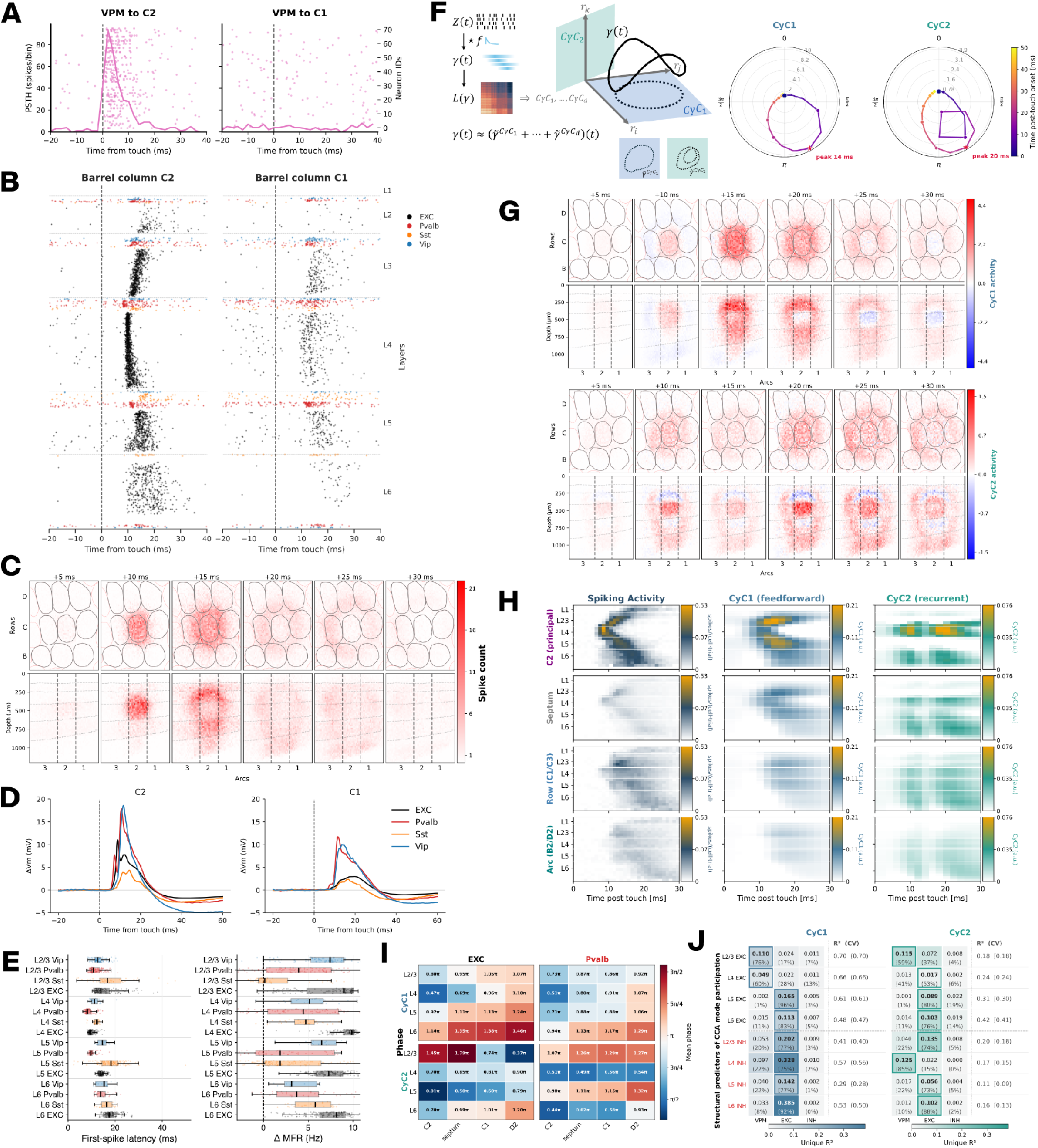
Cortical response to active touch of the C2 whisker. **A.** VPM input during active touch. Fibres projecting to C2 barrel column (left) and C1 barrel column (right), aligned to touch onset. Spike raster on the right axis, PSTH on the left, 2 ms bins. **B.** Cortical spike raster for C2 column (left) and C1 column (right), grouped by molecular class and ordered by depth within class - EXC (black), Pvalb (red), Sst (orange), Vip (blue). Lamp5 cells are too sparsely active during touch to plot and are omitted from B, D and E. **C.** Spatiotemporal spread of the touch response, in 5 ms frames from touch onset. Colour intensity gives the spike count summed over the cells of each flattened 3D voxel. *Top:* flatmap plane across the 3 3 barrel grid, barrel outlines drawn. *Bottom:* coronal view along arcs 1-3, dashed lines marking the C2 barrel column boundary. **D.** Mean membrane potential change per molecular class, for C2 column (left) and C1 column (right). **E.** Touch response characterisation per molecular class, C2 barrel column. *Left:* first-spike latency, first spike within 50 ms of touch onset. *Right:* change in mean firing rate, 0-100 ms after touch relative to the 100 ms before. One value per cell, averaged over trials. Boxes give the quartiles and the median, whiskers extend to 1.5 the interquartile range, outliers not drawn; overlaid points are a random sample of up to 150 cells per class. **F.** Cyclicity analysis decomposition. *Left:* the spike raster *Z* is filtered into continuous signals *γ* (2 ms PSTH, exponential kernel *τ* = 5 ms), from which a lead matrix *L* is built, each entry giving the lead between a pair of neurons. Eigendecomposition of *L* returns pairs of eigenvectors spanning orthogonal planes, the cyclic components CyC. CyC1 is the plane maximising the enclosed area of the projection of *γ* onto it; CyC2 is obtained the same way on the residual *γ γ̃*^CyC1^, capturing deviation from the CyC1 plane. Each response is approximated by the sum of its projections, additional loops appearing as higher frequencies. The number of components retained is set from the reconstruction error curve (Supplementary Figure S8D). *Right:* population trajectory in the complex plane of CyC1 and CyC2, radius giving mode amplitude and angle giving phase, coloured by time from touch onset. Peak times marked. **G.** Spatiotemporal structure of CyC1 (top) and CyC2 (bottom) participation, plotted as in C. **H.** The same response as a continuous time course rather than in binned frames. Cortical depth on the vertical axis, time from touch on the horizontal, summed over the area named at left - C2 column, septum, row neighbours (C1/C3) and arc neighbours (B2/D2). *Left:* spiking activity. *Middle:* CyC1. *Right:* CyC2. Colour scales are two-slope and non-linear for spiking activity, to keep the intensity of the response visible across areas. **I.** Circular mean phase across cells, per layer and barrel group, for CyC1 (top) and CyC2 (bottom), EXC (left) and Pvalb (right). CyC1 phase advances in a columnar spread, CyC2 in a horizontal, laminar one. **J.** Structural predictors of cyclic component participation. A complex-valued generalised linear model, in which the complex response encodes mode amplitude and phase jointly, was fitted per cell class using afferent synaptic drive alone, asking how far participation follows from static connectivity. Unique *R*^2^ of each predictor group - VPM, cortical EXC, cortical INH - for CyC1 (blue) and CyC2 (green), with each group’s share of the summed unique *R*^2^ in parentheses. *Right of each block:* full-model *R*^2^, cross-validated *R*^2^ in parentheses. Boxed cells mark the dominant predictor. Dashed line separates excitatory from inhibitory classes. Classes are excitatory cells per layer and all inhibitory cells of a layer pooled. Panels C, D, G and H are averaged over 5 seeds, 14 trials each. The cyclicity decomposition in F-J was obtained from one seed.

#### Response across layers and classes

We present the results for the principal column C2 and for its neighbours side by side throughout the analysis, taking C1 column as the illustrated neighbour and pooling C1 column and C3 column as the row neighbours where numbers are given. Touch drives a fast, dense response in C2 column and a delayed, sparse one in C1 column (Figure 6B, D). Mean first-spike latency is 13.9 4.4 ms in C2 column against 21.1 5.9 ms in the row neighbours (mean SD across cells), and the mean firing rate change 7.8 5.9 Hz against 4.1 4.6 Hz. In the principal column, the population response orders by layer, earliest in layer 4, then layers 2/3 and 5, and latest in layer 6. We found Pvalb cells to lead excitatory cells of the same layer, by 1.2 ms in layer 4, 3.7 ms in layer 2/3, 3.8 ms in layer 5 and 3.4 ms in layer 6, and Sst cells to activate last in layers 2/3, 4 and 5. Overall, Pvalb and excitatory cells carry the largest and fastest depolarisation. Deep layers also receive direct thalamic input, since the VPM profile carries a second peak at the layer 5 and layer 6 border. This behaviour matches the experimental findings reported in mouse barrel cortex during active touch, where fast-spiking interneurons of the layers driven by the thalamus respond earliest and most strongly, excitatory neurons of the same layers follow, and Sst neurons respond latest and most weakly, both in spiking (Yu et al., 2019) and subthreshold (Kiritani et al., 2023).

In C1 barrel column, excitatory cells spike less reliably, but the delayed onset of membrane potential depolarisation is still clear, so the confinement of the response to the principal column is much sharper above the action potential threshold. Layer 4 excitatory cells respond to 98% of touches in C2 barrel column and to 55% in the row neighbours; for layer 6 the values are 76% and 36%. The C2 column depolarisation is also more transient, large at its peak and quick to return, while the C1 column response is flatter and slower (Figure 6D). Pooled across layers, excitatory cells depolarise by 19.1 0.3 mV at the peak in C2 column against 4.1 0.1 mV in C1 barrel column (mean SD across seeds). Pooled across layers, excitatory cells depolarise by 19.1 0.3 mV at the peak in C2 column against 4.1 0.1 mV in C1 barrel column (mean SD across seeds), and in layer 4 the depolarisation stays above half its peak for 1.6 0.05 ms in C2 column against 12.8 0.1 ms in C1 column. In C1 column, the inhibitory classes dominate both the membrane potential and the spiking response. Pvalb depolarises by 16.8 0.5 mV in layer 2/3 and 19.2 1.0 mV in layer 5, against 2.7 0.1 mV and 4.6 0.2 mV for the excitatory cells of the same layers, and reaches its peak first, at 11.8 0.3 ms and 11.5 0.2 ms. The depolarisation is followed by a hyperpolarisation across classes, and Sst cells in layers 2/3 and 5 never depolarise on average (Supplementary Figure S7B), their mean deviation over the response window dropping to 0.46 0.05 mV and 0.40 0.05 mV (mean SD across seeds).

Vip cells respond as strongly as Pvalb cells in both columns in mean firing rate, though less strongly in membrane potential in C2 column (peaks of 23.3 0.5, 19.4 1.1 and 19.1 0.6 mV in layers 2/3, 4 and 5, against 26.6 0.5, 23.4 1.2 and 32.0 0.8 mV for Pvalb), and more strongly than reported in vivo, where touch only weakly modulates Vip (Yu et al., 2019). In C2 column, the mean firing rate change of Vip cells reaches 8.3 8.5 Hz in layer 2/3 against 8.2 18.5 Hz for Pvalb, and 6.7 12.2 Hz in layer 4 against 6.7 10.6 Hz. They arrive later: their first spikes follow those of the layer 4 excitatory population rather than accompanying the Pvalb response, and cortical excitatory input outweighs VPM as a predictor of their participation in the response (Figure 6J). The Vip response is therefore delivered through the cortical circuit rather than directly by the thalamus.

Resolved per class and layer, the difference between the columns is one of timing, reliability and strength (Supplementary Figure S7A-E). First-spike latencies in C2 barrel column are tight and clustered, whereas in C1 barrel column they are later and scattered, with no common onset across cells. Layer 4 excitatory cells respond at 9.96 1.31 ms in C2 column against 20.43 4.90 ms in the row neighbours (mean SD), and layer 6 excitatory cells at 17.87 2.78 ms against 24.70 4.27 ms. The laminar profile changes with it: the granular-to-deep progression of C2 column gives way to activity concentrated in layer 4 and the supra-granular layers. Responses in C2 column are reliable across trials in the excitatory and Pvalb populations, and stronger than in C1 column both above and below threshold (Supplementary Figure S7C).

Three population responses stand out. Pvalb cells of layer 5, rather than of layer 4, carry the largest mean touch-evoked rate increase in C2 barrel column, 12.3 24.3 Hz against 6.7 10.6 Hz, driven by a subset of cells (median 1.9 against 4.4 Hz), in contrast to the whisking state, where layer 4 Pvalb cells respond first and the other layers follow more gradually. Sst cells in C2 column are recruited by touch across layers 2/3 to 5, with a delay, which contrasts with their hyperpolarisation at whisking onset. Layer 6 excitatory cells give the latest membrane potential change in the column, peaking at 17.7 ± 0.3 ms against 8.74 ± 0.05 ms in layer 4, and the largest sustained one among the excitatory classes, their mean deviation over the full response window reaching 3.8 ± 0.05 mV against 0.87 ± 0.06 mV in layer 4. The deep-layer response outlasts the granular one.

#### Spatiotemporal spread

Within C2 column, the response follows a laminar sequence (Figure 6C, H, left). It appears first in layer 4, at a mean first-spike latency of 10.0 1.5 ms, and spreads from there into layers 2/3 and 5 at 15.0 4.8 ms and 14.3 3.1 ms and into layer 6 at 17.6 2.9 ms. Within layers 5 and 6, latency increases smoothly with depth, from 13.6 ms in the excitatory cells closest to layer 4 to 19.4 ms in the deepest, so the response reaches the deep layers from above rather than arriving there directly. Almost every cell of the column responds. The deep response persists after the granular one has decayed: the population rate falls back below a fifth of its peak at 12.5 ms in layer 4, 17.5 ms in layer 2/3, 18.5 ms in layer 5 and 23.5 ms in layer 6, so the column returns to baseline from layer 4 outward.

Laterally, the response leaves the C2 barrel column into the septum and the neighbouring barrels, delayed and weaker than the columnar response. In the C2 column, mean first-spike latency is 13.9 ± 4.4 ms and the mean firing rate change is 7.8 ± 5.9 Hz. In the septum, latency rises to 17.2 ± 5.2 ms and the rate change falls to 5.9 ± 5.4 Hz. In the row neighbours, latency is 21.1 ± 5.9 ms and the rate change 4.1 ± 4.6 Hz, and in the arc neighbours, 22.5 ± 6.8 ms and 2.4 ± 4.0 Hz. The septum keeps the laminar order of the principal column, with layer 4 earliest at 13.8 4.0 ms. In the neighbouring barrels, the population response also begins in layer 4 (Figure 6H), but fewer cells respond and their first-spike latencies are scattered, so mean latencies are less clearly ordered by layer.

### Cyclic decomposition of the touch response

#### Decomposition and component selection

To further investigate the structure of the touch response, we decomposed the spiking activity using cyclicity analysis (Figure 6F, left). Cyclicity analysis is built on lead-lag relations: it compares every pair of neurons to determine which becomes active first, and combines these comparisons into an order of activation across the population. The analysis was applied to all neurons of the simulated volume together, giving a single set of components that describes the response across the whole grid. The decomposition returns a set of orthogonal planes, the cyclic components, ordered by the area enclosed by the population trajectory projected onto them. CyC1 is the plane maximising that area, and CyC2 is obtained the same way on the residual *γ γ̃*^CyC1^, capturing deviation from the CyC1 plane. The planes come from the lead matrix *L*, built from the filtered raster *γ* (2 ms PSTH, exponential kernel *τ* = 5 ms), each entry giving the lead between a pair of neurons. *L* is skew-symmetric, so its eigenvalues are purely imaginary and occur in conjugate pairs, and each pair spans one of the planes. The response is approximated as the sum of its projections, with additional loops appearing as higher frequencies.

Cyclicity analysis differs from principal component analysis in what it maximises. Principal components are built from the covariance matrix and rank the cells by shared variance, which discards the order in which they were active. Here the components are built from the lead matrix and rank planes by the area enclosed by the common trajectory, so the temporal relations between cells are retained, and each cell carries a phase within the response.

We retained two components. The reconstruction error curve changes steeply over the first three components and flattens beyond them (Supplementary Figure S8D), so the gain from a third is real but small. CyC3 was examined and set aside, as it carries a complex triple-looped trajectory with no interpretable spatiotemporal organisation. Retaining two is therefore a decision made on interpretability, with the reconstruction error setting the upper bound at three.

CyC1 traces a single loop in the complex plane, with radius giving the strength of the population response in that mode and angle giving phase, while CyC2 traces a repeating path, passing through similar states twice within the response (Figure 6F, right). The two peak at different times from touch onset, CyC1 at 14 ms and CyC2 at 20 ms. The decomposition was obtained from a single simulation seed. Spatial averages throughout this section pool five seeds of 14 touches.

#### Spatiotemporal structure of the two modes

Projecting the activity back onto each plane gives a partial reconstruction of the response, the signal carried by that component alone. Each plane approximates the signal by a single harmonic, so a reconstruction can take negative values over parts of the time course, which get corrected by the higher components. Reconstructed in this way, CyC1 and CyC2 can be placed back on the flatmap and followed through the grid as the spiking response was, in 5 ms frames (Figure 6G) and as a continuous time course by cortical depth, summed over area (Figure 6H, middle and right). The strongest participation in the whole simulation grid is the early response of C2 barrel column in layers 2/3 and 4, between 8 and 12 ms after touch, and it is the strongest feature of the spiking response as well. It contributes to both modes, so the two are not separated by where the response begins but by how it propagates.

CyC1 is focused on the principal column. Participation is concentrated in C2 column and falls off with distance from the barrel, and its time course follows the spiking response, beginning in layer 4 and extending through the depth of the column. Little of the mode is carried by the septum or the neighbouring barrels. CyC2 is distributed across the grid, organised in laminar bands that span the barrels and the septum. We can, however, observe a strong component in layer 4 of C2 barrel, where the response begins. Over the time course, the signal repeats, with a second wave of strong amplitude following the first.

Counting the cells in the top 10% by radius gives the same picture (Supplementary Figure S8A). C2 barrel column is the most enriched region of the grid, followed by the septum and the row neighbours, and enrichment falls off with distance from the principal barrel. A large share of these cells participate in both CyC1 and CyC2. The two components are orthogonal by construction, but their strongest participants overlap substantially; what is divided between them is the activity of those cells rather than the cells themselves.

#### Phase gradients and radius magnitude

Each cell has a preferred phase within each component, so within the molecular classes we can reduce to a single phase value per subset area (Figure 6I). Here we confirm and quantify the spatiotemporal observations about the contributions of the two components. CyC1 phase has a clear columnar spread in both EXC and Pvalb cells, originating from layer 4. In C2 barrel column, the phase is earliest in layer 4 and advances upward into layer 2/3 and downward into layers 5 and 6, and at each depth it advances again with distance from the principal barrel, through the septum and the row and arc neighbours in turn. CyC2 phase has a more laminar behaviour, in particular for Pvalb cells, where the phase is close to constant across the barrel groups within a layer and differs between layers instead. The excitatory cells show the same lack of a radial gradient, though their phase is less consistent between layers.

Sst and Vip cells are less ordered in both components: neither carries a consistent phase gradient, laminar or radial. Their radii also fail to stand out. Sst has the smallest radius of any class, and Vip is moderate but unusually uniform across layers and barrel groups (Supplementary Figure S8C). Layer 5 Pvalb cells carry a large radius in both components and, unlike the excitatory cells, retain it across the septum and the neighbouring barrels, so their participation is not confined to the principal column. Across the major molecular classes, no single one dominates either component (Supplementary Figure S8B). Every class divides into cells reaching the threshold for CyC1 alone, for CyC2 alone, and for both, and all three groups are present in each class. Pvalb and Vip carry the largest participating fractions, peaking in layer 5 at about half the class, against under a fifth for the excitatory classes.

#### Structural predictors of participation

Radius and phase describe how a cell behaves during the response, while its connectivity is fixed. We asked how far participation follows from static connectivity alone. A complex-valued generalised linear model, in which the complex response encodes amplitude and phase jointly, was fitted per cell class using afferent synaptic drive as the only predictor (Figure 6J). For each component, we report the unique *R*^2^ of three predictor groups: thalamic input from VPM, cortical excitatory input and cortical inhibitory input.

CyC1 participation is far better predicted than CyC2 participation, with a full-model *R*^2^ of 0.53 0.14 (0.29-0.70) for CyC1 against 0.22 0.10 (0.11-0.42) for CyC2 (mean SD across the eight cell classes, range in brackets). Cross-validated *R*^2^ was close to the full-model value (0.52 0.14 and 0.21 0.10), so the fits are not overfitted. The columnar geometry follows from static connectivity to a greater extent than the laminar one. For CyC1, VPM dominates in L2/3 and L4 EXC, and cortical excitatory input dominates in the deeper excitatory classes and in all four inhibitory groups. Predictability declines with depth in the excitatory classes. For CyC2, VPM dominates in L2/3 EXC and L4 INH, and cortical excitatory input dominates elsewhere; the two are close to equal in L4 EXC. Between the two components, the VPM share falls in the superficial excitatory classes and holds elsewhere. Cortical inhibitory input contributes little throughout, the largest share being 19% for L5 EXC in CyC2.

Two qualifications apply. The unique *R*^2^ values are lower bounds: they sum to far less than the full-model *R*^2^, so most of the explained variance is shared between the predictor groups and is not attributable to any one of them. The CyC1 against CyC2 contrast is less affected, as the same sharing applies to both. The full-model *R*^2^ is also bounded, since a static measure of connectivity cannot capture the part of participation that arises from the dynamics of the response.

### Touch responders, choristers and leaders

#### Defining the populations

We complement the decomposition with a closer look at single cells, defining populations in the touch and whisking conditions separately and asking whether the resulting groups coincide.

First, we focus on the cells active during touch. Most neurons do not respond consistently to every touch, so we search for those whose spiking reports the stimulus on nearly every trial, at a fixed latency, and separably from their own baseline. Four criteria enforce this: modulation index at least 0.5, reliability at least 0.95, jitter at most 2.5 ms and *d*^′^ at least 1.35 (Figure 7A, left). We find 1,298 such cells, 695 of them in the C2 barrel column. The remainder of the section focuses on C2 column alone.

**Figure 7.**
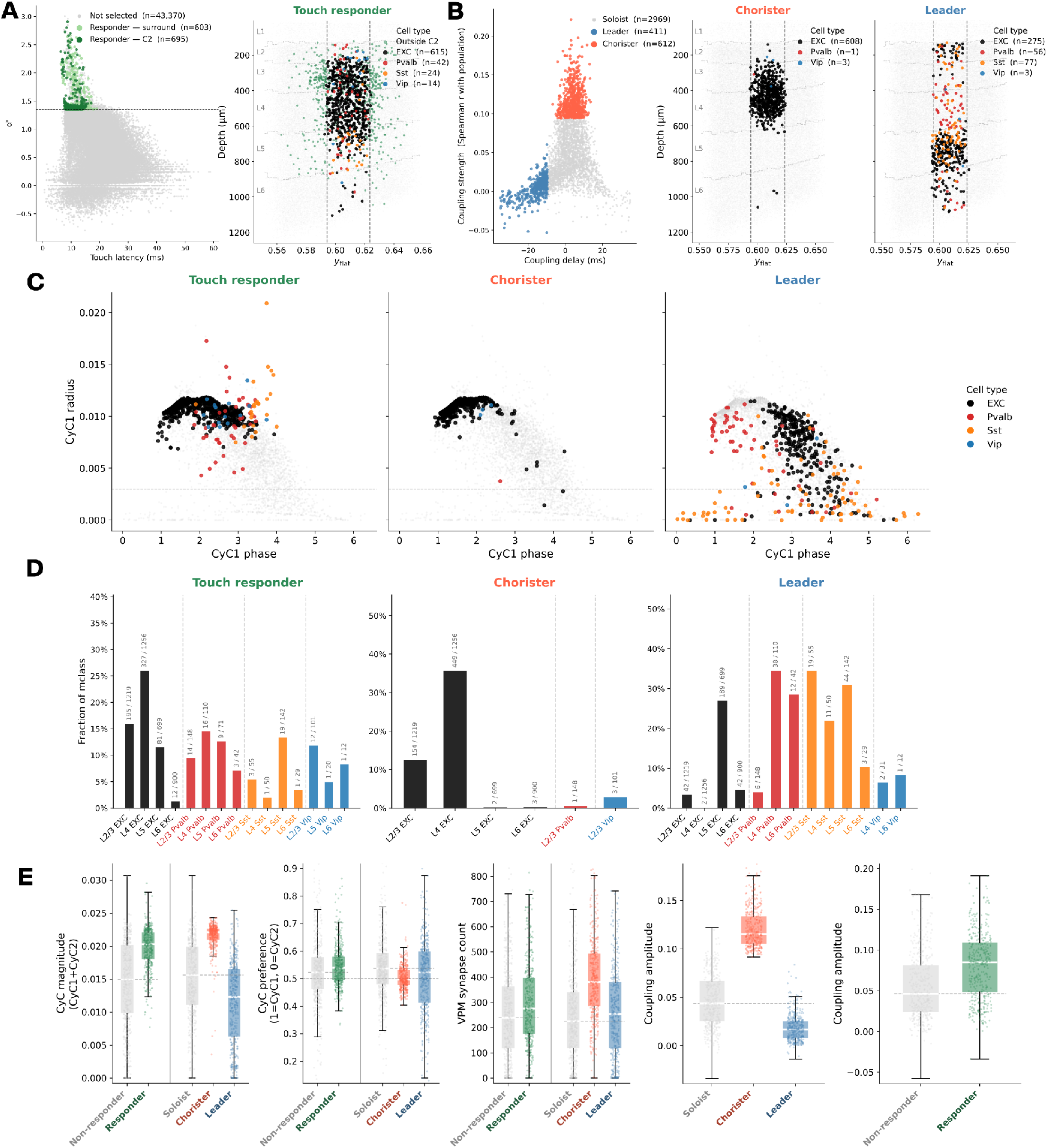
Functional cell populations during touch and whisking. **A.** Touch responder selection. *Left: d*^′^ against touch latency for all cells; responders pass modulation index 0.5, reliability 0.95, jitter 2.5 ms and *d*^′^ 1.35 (dashed line). Light green marks responders outside the C2 barrel column, reported here only; all subsequent panels are C2 column. *Right:* selected responders in the C2 column, plotted by depth against flatmap position, coloured by molecular class. Grey points are all remaining cells. **B.** Population coupling classification during whisking. *Left:* coupling strength against coupling delay for all cells, giving choristers (top 15% by strength) and leaders (earliest 10% by delay); all remaining cells are soloists. *Middle* and *right:* choristers and leaders in the C2 column, plotted as in A. **C.** CyC1 participation of each population. Radius against phase for touch responders (left), choristers (middle) and leaders (right), coloured by molecular class; grey points are all cells. Dashed line marks the radius threshold of 0.003. CyC2 in Supplementary Figure S10. **D.** Fraction of each cell class assigned to the three populations. Counts above each bar give selected cells over class size. Note the differing *y*-axis ranges. **E.** Response and connectivity properties of the three populations. Left to right: combined CyC1 and CyC2 magnitude; CyC preference, from 0 (pure CyC2) to 1 (pure CyC1); VPM synapse count; coupling delay; coupling strength. Touch responders are compared against non-responders, choristers and leaders against soloists, as marked. Boxes give quartiles and median, whiskers 1.5 times the interquartile range, points individual cells. Dashed lines give the median of the comparison group. Lamp5 cells are too sparsely active to pass the spike-count filter for any of the three populations and are omitted throughout, as in Figure 6. Touch panels pool 5 simulation seeds, whisking panels 25.

Second, we study the cells during whisking, asking whether a cell fires together with the surrounding population or independently of it, and how far it leads or follows the activity. We address this with the approach of Bimbard et al. (2026). For each cell, we take the summed firing rate of all other cells and average it around the times of that cell’s own spikes, giving the spike-triggered population rate (Supplementary Figure S9A). A cell locked to collective activity produces a peak in this profile, while a cell driven independently of its neighbours produces a flat one. The height of the profile at zero lag, the moment the cell itself spikes, gives the coupling strength, separating strongly coupled choristers from weakly coupled soloists (Okun et al., 2015). Its centre of mass gives the coupling delay, negative in leaders, which fire ahead of the majority of their neighbours (Luczak et al., 2015). We take the top 15% by strength as choristers and the earliest 10% by delay as leaders, leaving 2,969 soloists (Figure 7B, left).

#### Anatomical distribution

The three populations differ in their laminar and class composition (Figure 7A, right, and B, middle and right). Touch responders sit mainly in layers 4 and 2/3, which contain 81.7% of them (568 of the 695 responders). They are dominated by excitatory cells at 88.5% (615 cells), with a sparse scattering in the deeper layers. Choristers are almost exclusively excitatory (99.3%, 608 of the 612 choristers) and confined to layer 4 and the deep part of layer 2/3, which together hold 99.2% of the population (607 cells). Leaders spread across the full depth of the column and carry a large inhibitory fraction (33.1%, 136 of the 411 leaders), mostly Sst (18.7%, 77 cells) and Pvalb (13.6%, 56 cells), alongside a substantial group of excitatory cells in layer 5 (189 cells). The fraction of each class assigned to each population separates them further (Figure 7D); here the denominator is the number of cells of that class in C2 barrel column. Touch responders draw from every class except L4 Vip, reaching 26.0% of L4 EXC (327 of 1,256 L4 EXC cells). Choristers come almost entirely from two classes, 35.7% of L4 EXC (449 of 1,256) and 12.6% of L2/3 EXC (154 of 1,219 L2/3 EXC cells), with no other class above 3.0% (at most 3 of 101 L2/3 Vip cells). Leaders invert this: L4 EXC contributes 0.16% (2 of 1,256), while 34.5% of L4 Pvalb (38 of 110), 28.6% of L6 Pvalb (12 of 42), 34.5% of L2/3 Sst (19 of 55) and 31.0% of L5 Sst (44 of 142) are selected, together with 27.0% of L5 EXC (189 of 699).

We then looked into the overlap between the populations (Supplementary Figure S10D). Choristers and leaders never co-occur, which the measures partly enforce: strength is read at zero lag, which an early-centred profile misses. Choristers overlap substantially with touch responders, 208 of 612 cells, most of all in L4 EXC, where 159 belong to both, while leaders share only 21 cells with the responders. The same L4 excitatory cells that fire with the whisking population also report touch reliably; the cells that lead the population do not.

#### Cyclic mode participation

The decomposition assigns every cell a radius and a phase in each component. The radius represents how strongly a cell contributes to that component, and the phase its timing within the cycle. The three populations can therefore be placed directly into the dynamics described at the population level. Touch responders carry a high CyC1 radius and occupy the early phases through to the middle of the cycle (Figure 7C, left). In CyC2, they spread across the full range of both phase and radius (Supplementary Figure S10A, left). Plotted against each other, the two phases cover the whole first half of the CyC2 cycle together with the transition band between the two components (Supplementary Figure S10B, left).

Choristers are more sharply confined. In CyC1, they form a single group of excitatory cells at high radius and early phase (Figure 7C, middle), and in CyC2 they again sit at high radius within the first half of the cycle (Supplementary Figure S10A, middle). Their two phases advance together, placing them on the excitatory diagonal (Supplementary Figure S10B, middle).

Leaders show neither pattern. Their CyC1 participation splits in two, an early Pvalb group and an excitatory group at mid-phase (Figure 7C, right). A large fraction of leaders also falls below the radius threshold, so much of the population contributes little to the component at all. In CyC2, they sit largely in the transition band, without concentrating in either phase or radius (Supplementary Figure S10A and B, right).

Two of the three populations map onto the decomposition. Touch responders align with CyC1 and with the early half of CyC2, expected given that both are extracted from the touch simulations. Choristers are the strongest and earliest contributors to both components while also being the cells most tightly coupled to the population activity during whisking, placing the same cells at the front of the touch response and at the centre of whisking activity. Leaders are the exception. Their spread in phase and their low radii mean that coupling delay and component participation are measuring different things.

#### Response and connectivity properties

We then compared the populations against their reference groups on response magnitude, component preference and thalamic input (Figure 7E). Touch responders carry a higher combined CyC1 and CyC2 magnitude than non-responders and lean towards CyC1. Choristers are raised above the soloists in magnitude, while leaders fall below them, so the split seen in their radii holds at the population level. All three populations sit close to the preference midpoint, and only the responders are shifted towards CyC1 relative to their comparison group; choristers sit marginally on the CyC2 side of the soloists.

The three also differ in their median thalamic input. Touch responders (*n* = 695) receive slightly more VPM synapses than non-responders (*n* = 4,190), 276 against 240 (interquartile ranges 176-398 and 118-363), and choristers (*n* = 612) stand out against soloists (*n* = 2,969), 382 against 226 (286-494 and 119-341), while leaders (*n* = 411) sit only marginally above the soloist median (254, 119-380). The final two panels report coupling delay and coupling strength, the measures on which the whisking populations were defined, and serve to characterise them rather than to add a further comparison. Across all three comparisons, the responders and the choristers behave alike, and the leaders behave differently. Two of the measures we applied, touch reporting and coupling to the whisking population, converge on the same L4 excitatory cells: they participate most strongly in both components, receive the most thalamic input, and overlap heavily in membership despite being selected in different conditions from unrelated quantities. Coupling delay selects a different population, one that shares none of these properties.

### Polarised versus symmetric circuits

Dendritic polarity in layer 4 is an anatomical feature of the barrel field, present in the reconstructions and reproduced in the model through boundary-guided synthesis. We compare two circuits directly: the polarised circuit (POL) confines layer 4 dendrites to the home barrel, while the symmetric circuit (SYM) grows the same cells without that boundary (Figure 8A). The comparison asks what dendritic polarity contributes to the spatial and temporal structure of sensory-evoked activity during active touch.

**Figure 8.**
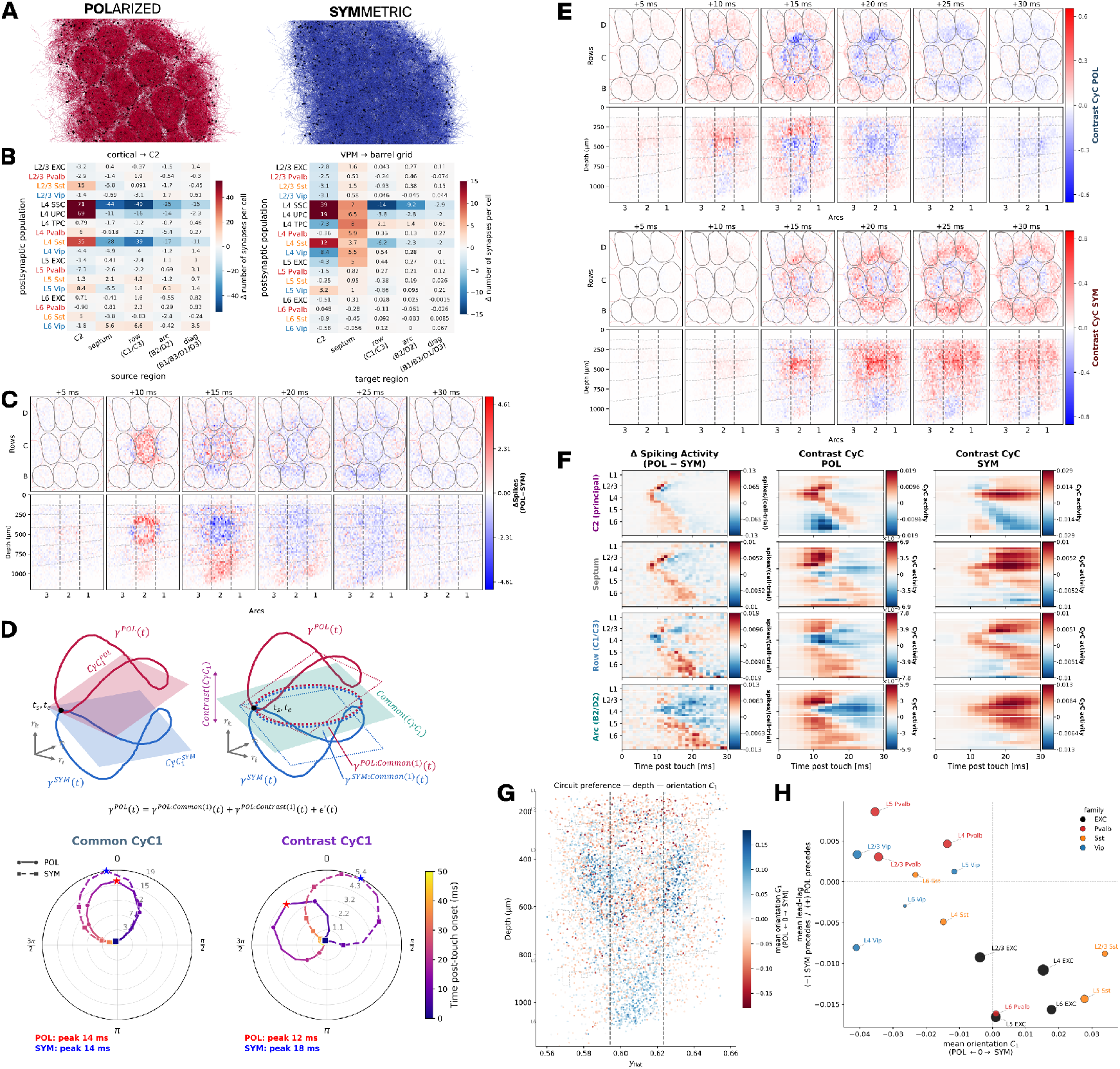
Functional comparison of the polarised and symmetric circuits. The two circuits share cell positions, axons, thalamocortical input optimisation and postsynaptic potential recipe. They differ only in the layer 4 dendritic constraint described in Figure 2: polarised (POL) dendrites are confined to the home barrel, symmetric (SYM) ones are not. All comparisons are POL SYM, so positive values indicate POL. **A.** Sample of L4 SSC dendritic trees across the barrel grid, POL (left) and SYM (right), somata in black. **B.** Difference in synapse count per cell between circuits, by postsynaptic population. *Left:* cortico-cortical inputs to C2 barrel column, by source region. *Right:* VPM inputs, by the region containing the postsynaptic cell. **C.** Difference in touch-evoked spiking between circuits, in 5 ms steps from touch onset, plotted as in Figure 6C. **D.** Common and contrast decomposition. *Top:* the projections of *γ*^POL^ and *γ*^SYM^ onto their cyclic components overlap substantially. For each CyC, the plane maximising that overlap is taken as the common plane and the remaining dimension assigned to contrast space, so each signal splits into activity shared with the other circuit and activity unique to it. *Bottom:* population trajectory in the common (left) and contrast (right) plane of CyC1, shown as the leading example of the three components used throughout. POL solid, SYM dashed, coloured by time from touch onset. Peak times marked. **E.** Spatiotemporal structure of the contrast activity in POL (top) and SYM (bottom), integrated over CyC1-CyC3 and plotted as in C. Negative values are part of the decomposed signal. **F.** Continuous time course by cortical depth, summed over the area named at left, as in Figure 6H. *Left:* difference in spiking activity. *Middle* and *right:* contrast activity in POL and in SYM. **G.** Spatial distribution of circuit preference. Contrast orientation per cell in the C2 column, plotted by depth against flatmap position. Orientation is an independent measure with its own sign, negative towards POL and positive towards SYM. Dashed vertical lines mark the C2 barrel boundary; dashed horizontal steps the layer boundaries. Panels C, E, F and G pool 5 simulation seeds of 14 touches for each circuit, matched between POL and SYM. The cyclicity decomposition was obtained from a single seed. **H.** Circuit preference per layer and molecular class. Horizontal: mean contrast orientation, negative towards POL. Vertical: mean lead-lag, negative when SYM precedes and positive when POL precedes. Marker size gives population size.

The constraint acts at synthesis, which allows every other property to be held fixed between the two circuits (Section *Barrel adaptation*). The Ornstein-Uhlenbeck background and the postsynaptic potential amplitudes were drawn independently in each circuit against the same targets (Methods, Section *Circuit comparison*). All comparisons in the next section are computed as POL vs SYM (POL-SYM), thus positive values indicate a preference for the POL circuit. The exception is the contrast orientation, which carries its own sign.

#### Structural differences

Removing the constraint changes the connectivity of the cells it applies to (Figure 8B). Layer 4 excitatory cells in C2 barrel receive more synapses from other C2 column cells in POL, 71 268 per cell for L4 SSC (*n* = 951) and 69 251 for L4 UPC (*n* = 139), and fewer from the septum and from the surrounding barrel groups (mean SD of the per-cell difference, POL SYM). The thalamocortical difference depends on where the postsynaptic cell sits. Layer 4 cells inside C2 barrel gain VPM synapses in POL, 39 132 per cell for L4 SSC and 19 114 for L4 UPC, cells of the same classes in the row, arc and diagonal barrels lose them, and septal cells gain a small number. Both effects concentrate on the constrained layer 4 classes and appear only weakly in the classes above and below. These differences follow from the dendritic constraint, and they are the structural starting point for the functional comparison. ‘

#### Difference in the touch response

We then looked more closely at the difference in the suprathreshold response of the two circuits. Cells share the same soma positions between the circuits, so the two responses can be compared directly in space and time. We subtracted the spike counts of SYM from POL within the same flattened three-dimensional voxels used in Figure 6C. The two circuits diverge from the first frames after touch (Figure 8C).

In the 10 ms frame, the difference is positive and confined to C2 barrel column, where it reaches a fifth of the POL response, with negative signal in neighbouring C-row barrels. Active touch evokes more spikes in POL inside the principal barrel and less immediately around it. At 15 ms we see positive values in infragranular layers of the C2 barrel column, whilst the barrel’s activity becomes negative, hence dominated by SYM signal. From 20 ms onwards the difference is weakly negative in C2 column, the septum and the arc, with a mean over 20-30 ms of −3.6%, −4.5% and −6.9% of the POL peak response in the respective area, and close to zero in the row (+0.4%).

The continuous time course resolves the difference (Figure 8F, left). In C2 barrel column, the difference follows the canonical laminar sweep of the response from layer 4 into layer 2/3 at 8 to 12 ms and then down through layer 5 and layer 6 by 20 ms. This positive band of POL activity is immediately followed by a negative band, indicating a time delay between the two circuits. The septum carries the same paired bands with a large positive infragranular difference at 15 to 22 ms. The row and arc neighbours reverse this ordering: their supragranular and granular response is larger in SYM from 10 ms, and their infragranular response is larger in POL from 15 ms. The row difference peaks later, at 18.5 ms, so the POL preference there arrives only after the principal column has been engaged. A positive band followed by a negative one at the same depth is the signature of a shift in timing rather than a change in magnitude, and the same pairing appears in every area.

The depth-by-time maps of the difference, repeated for each of the four main molecular classes (EXC, Pvalb, Sst, Vip), place the effect in the excitatory population (Supplementary Figure S11A-D). The excitatory difference reproduces the pooled pattern in all four areas. The Pvalb difference is small and positive from 10 to 20 ms without banding. The Sst difference is negative in layer 2/3 of C2 barrel column over the same window, so Sst runs against the other classes there. The Vip difference is sparse and weakly positive in the septum and the row neighbours after 12 ms. Membrane potential shows the paired bands most clearly, with a positive leading edge at every depth and a large positive infragranular difference at 15 to 25 ms in all four areas (Supplementary Figure S11E).

The two circuits produce a similar response. The time-resolved difference shows where and when the circuits diverge, but a subtraction alone cannot separate how much of that difference is a change in amplitude and how much a shift in timing. We therefore decomposed the response into components in which amplitude and timing are represented independently.

### Common and contrast decomposition

We applied the cyclicity decomposition to the touch response of both circuits. We observe two signals *γ*^POL^ and *γ*^SYM^ whose projections onto their cyclic components overlap substantially. For each *CyC*, we construct a plane where the overlap between the two projections is maximal, the common plane, and assign the remaining dimension to a contrast space. Each signal can then be expressed as a sum of an activity highly similar in both signals and activity unique to that signal (Figure 8D, top). We applied the decomposition to the first three components, showing CyC1 as the leading example and studying the sum of the three in the spatiotemporal maps below.

In the common plane of CyC1, which is shared by the two circuits by construction, the two trajectories nearly coincide, both peaking at 14 ms (Figure 8D, bottom left). Their spatiotemporal structure also matches: onset in C2 column at granular depth by 10 ms, spread across the barrel grid and the septum by 15 to 20 ms, and decay by 30 ms (Supplementary Figure S12). The common activity accounts for 95% of the signal power in POL and 90% in SYM, and the two trajectories correlate at *r* > 0.99. It carries the touch response described in Figure 6, and dendritic polarity leaves it unchanged.

The contrast trajectories separate the two circuits. POL peaks at 12 ms and SYM at 18 ms (Figure 8D, bottom right). Each contrast component can be read as the activity present in one circuit, and not in the other, so the two are described separately.

The activity unique to POL is early and confined to the principal barrel (Figure 8E, top). It appears in C2 barrel column at granular depth and peaks by 10 ms. By 15 ms it has reversed sign inside C2 column and moved to the edges of the grid, and it has decayed by 25 ms. By depth and time, it runs from layer 2/3 and layer 4 to the deeper layers between 7 and 15 ms in C2 column and in the septum, and this granular band is absent from the row and arc neighbours (Figure 8F, middle).

The activity unique to SYM is late and spread across the grid (Figure 8E, bottom). It is absent before 15 ms, appears first in C2 barrel column, and covers every barrel by 25 to 30 ms. By depth and time, it is mostly concentrated in layer 4 in C2 barrel, where it forms a single sustained band from 15 ms to the end of the window without descending through the layers (Figure 8F, right). The septum follows later than in POL and stays active longer, and in the row and arc neighbours the activity includes layer 4 rather than avoiding it. The same ordering holds in each molecular class (Supplementary Figure S11A-D, middle and right). Negative values in the contrast maps are part of the decomposed signal rather than a deficit, since each component approximates the response by a single harmonic, corrected by the higher components.

The two circuit-specific signals therefore differ in when they arrive and in where they sit. In POL, the signal is early and stays within C2 barrel column and the septum, and layer 4 of the neighbouring columns is excluded from it. Polarisation therefore sharpens the response in space and advances it in time. The single-cell counterpart of both follows from the circuit preference of individual cells.

#### Circuit preference of individual cells

We then asked whether individual cells contribute preferentially to the contrast activity of one circuit. The contrast plane of CyC1 separates them geometrically: the two responses trace distinct loops that both start at the origin and run in opposite directions, one belonging to POL and one to SYM. Each cell has a fixed position in the same plane, and cells positioned along one loop are the ones whose spikes build it. We quantified this per cell as an orientation, running from 1 for cells building the POL loop to 1 for cells building the SYM loop. Alongside, we computed the lead-lag of each cell, the difference in the time at which its contrast activity peaks in the two circuits, positive when the peak comes earlier in POL and negative when it comes earlier in SYM.

Orientation measures the contribution of each cell to the contrast activities of CyC1, from 1 for cells that contribute only to the contrast component of POL to 1 for cells that contribute only to the contrast component of SYM. It is defined per cell and can be mapped across the volume. Of 12,417 cells passing the participation floor, 476 are SYM-oriented and 313 POL-oriented; the rest lean to neither circuit. Mapped by depth against flatmap position, cells of both preferences occur throughout the grid, with a small number of dense groups standing out against that background (Figure 8G). The clearest is a SYM-oriented group of layer 4 excitatory cells occupying all the barrels neighbouring C2 column, densest along the sides they share with it, while C2 column itself is spared (Supplementary Figure S13C). Outside C2 barrel, 9.5% of layer 4 excitatory cells are SYM-oriented; inside it, 0.9%. A second SYM-oriented cluster sits in layer 6 of C2 column. POL-oriented cells are sparser and spread across the whole space, denser in layer 4 and in layer 2/3 around C2 column, where they are mostly Pvalb and Vip cells. The most extreme POL values of the panel are found in layer 2/3 inhibitory neurons: of the twenty most POL-oriented cells, eleven are layer 2/3 Vip or Pvalb, and none are excitatory.

Orientation and lead-lag separate the same populations (Figure 8H). Each population is placed by its mean orientation on the horizontal axis, negative towards POL, and by its mean lead-lag on the vertical, positive when its contrast activity comes earlier in POL. Three groups emerge. POL-oriented and POL-leading, in the upper left quadrant: Pvalb and Vip in layer 2/3, layer 4 and layer 5, with layer 5 Pvalb furthest out. POL-oriented and SYM-leading, in the lower left: layer 4 Vip, layer 4 Sst and layer 6 Vip. SYM-oriented and SYM-leading, in the lower right: layer 4 EXC, layer 6 EXC and layer 2/3 and layer 5 Sst. Layer 2/3 EXC, layer 5 EXC and layer 6 Pvalb sit on the zero-orientation line while leading in SYM, so they have no preference between the circuits and are timed by SYM alone. No population is SYM-oriented and POL-leading.

Circuit preference is also visible in which cells qualify as touch responders. Applying the criteria of Figure 7A to both circuits gives 1,298 responders in POL and 1,230 in SYM, 695 and 646 of them in C2 barrel column. Among layer 4 excitatory cells, those qualifying in one circuit alone divide by position (Supplementary Figure S13A). In C2 column, 230 cells are responders only in POL against 195 only in SYM, whereas in the surrounding barrels 135 are responders only in SYM against 98 only in POL.

## Discussion

### Model

We presented a morphologically detailed model of mouse barrel cortex (Figure 1B,C), built in atlas space with barrel-column geometry and thalamocortical projections. No previous model of the region reaches this level of detail. No dataset has been assembled specifically for bottom-up modelling of this kind, so the model’s components were sourced from existing data. The rat non-barrel somatosensory model drew on both rat and mouse sources, and the resulting recipes are available to build upon (Isbister et al., 2024; Reimann et al., 2024). We therefore based our model on the workflow for the rat somatosensory cortex, rather than reconstructing a circuit bottom-up from scratch. Densities, morphological synthesis, me-type composition, apposition-based connectivity and the thalamocortical fibre model were built with an established pipeline. New here are the mouse barrel cortex constraints and a comparison that isolates what dendritic polarity contributes to the touch response.

The rat datasets behind that pipeline were curated for the previous model and have no mouse equivalent: paired recordings of synapse kinetics and short-term dynamics, dendritic reconstructions, and the whole-cell recordings behind the e-models. To this we added mouse data where available: interneuron densities and the layer 4 excitatory composition, the barrel and septum density differences, postsynaptic potential amplitudes per pathway, thalamocortical depth profiles and absolute thalamocortical synapse density. Rat provides the form of the biophysics, and mouse its scale.

We annotated the 33 barrel columns in CCFv3 in previous work, together with a laminar coordinate system and a flatmap of the region (Bolaños-Puchet, Teska, et al., 2024). Every subsequent step depends on that annotation. Barrel and septum become distinct compartments, so layer 4 Vip and Sst densities can differ between them (Almási et al., 2019; Argunşah et al., 2026); VPM fibres can be confined to individual barrel outlines while POm is spread across the field; and boundary-guided synthesis becomes available, biasing dendritic growth against the wall of a cell’s own barrel. The flatmap gives the horizontal coordinates on which every spatial analysis in this study is built.

### Dendritic polarity

Dendritic polarity had not been measured quantitatively. We measured it across morphological classes, asking which classes carry it and whether it is confined to layer 4. Testing this required assembling a collection of morphologies. We drew reconstructions from public repositories and published datasets, curated them to a common standard and mapped them onto a single morphological-class scheme, giving 324 dendritic reconstructions from mouse barrel cortex. Polarity was measured from the basal dendritic segments of each cell as their length-weighted angular distribution in the flatmap plane (Figure 2A), reduced to a single score that rises with directional coherence and falls with symmetric bimodality. The same descriptors were applied to every cell, with no class-specific tuning, so classes can be ranked against one another.

Polarity is strongly class-dependent and concentrated in layer 4 (Figure 2C). We find that spiny stellate, non-Martinotti somatostatin and parvalbumin neurons are enriched in polarity above the population fraction; the supragranular and infragranular pyramidal types are not. These results match the literature findings in terms of morphological classes, and the property does not extend to the classes outside layer 4. The analysis was implemented to decide which classes to constrain. The per-class fractions are descriptive statistics of the assembled sample, and the first class-wide measure of polarity for mouse barrel cortex (Section *Limitations*). Polarity is a continuum (Figure 2B); the split into polar and low/moderate regimes was made for model building, which required a list of classes to constrain, and is not evidence of two populations.

We then used our polarity results to constrain the morphologies during synthesis. The three enriched classes were constrained, together with layer 4 untufted pyramidal cells, which showed no enrichment in our sample (*p* = 0.62, Figure 2C) but are reported as polarised (Staiger, 2004). Layer 4 spiny stellate cells were grown entirely from the curated mouse reconstructions, and the other constrained classes from a mixture of those and the adapted rat library. Most of the library remains rat, rescaled to mouse cortical thickness. The boundary-guided algorithm allowed us to control dendritic growth, such that each tree is largely confined to the column its soma occupies (Figure 2D,E, Supplementary Figure S2D). Synthesis gives every neuron a unique dendritic tree, whilst following the statistical properties of the reconstructions. Each morphology is paired with its own e-model. The boundary-guided algorithm is imposed during synthesis, so every other parameter of circuit building is preserved. Polarity therefore enters the model as a measured property of specific layer 4 classes rather than an assumption, and becomes a single anatomical property that can be removed with everything else held: the polarised and symmetric circuits.

### Connectivity

The model’s connectome is built from anatomy alone, with no rule imposing how likely one cell type is to connect to another. Model building imposes bouton density, laminar axon and apical dendrite targeting, and a revised statistic of synapses per connection. The rules are applied per morphological class and inherited from rat models (Markram et al., 2015; Reimann et al., 2024). The approach constrains the morphologies rather than imposing which cells connect. Several features of the connectome follow from that.

Connection probability between molecular classes (Figure 3F, G) is an output, not a target. No rule in the model acts on molecular class. Rules are defined on morphological and e-types, and the mapping to molecular identity is applied afterwards. Across the barrel grid, connectivity concentrates into the nine diagonal blocks (Figure 3G). No rule in the recipe references a barrel column, so the concentration follows from atlas geometry and axonal extent. With a small pool of exemplar morphologies, neurons of a class are copies of one another, and shared partners are inflated by that duplication rather than by circuit structure (Reimann et al., 2017). Synthesising a unique tree for every neuron removes that inflation: the common-neighbour bias we report in layer 4 (Supplementary Figure S3F) is a property of the connectome, not of the morphology pool. Sst receives the largest share of inhibitory drive in the C2 column (Supplementary Figure S3E), dominated by Pvalb rather than Vip. The recipe contains no inhibitory-to-inhibitory targeting rule, so the asymmetry follows from the morphological structure.

One feature of the connectome is imposed more directly. The translaminar motif carried by the pyramidal populations (Figure 3F) follows from placement rules that direct their axons to specific layers. We applied one deliberate change to the connectome algorithm. The linear apposition-to-synapse relation retains connections formed by one or two contacts, where the original square-root relation removed most of them. The resulting distribution sits closer to dense electron-microscopic reconstruction of mouse cortex (Schneider-Mizell et al., 2025). Connections in the model are therefore more often formed by a single synapse (Figure 3C,D).

The synaptic physiology is only partly mouse. Kinetics and short-term dynamics come from rat paired recordings, and only postsynaptic potential amplitudes were recalibrated, from sources reporting a small fraction of the pathways in the model. The amplitude recipe is largely extrapolated across layers and cell classes (Supplementary Figure S3C). The assignment of pathways to facilitating and depressing families (Figure 3A,B) originates from rat and is generalised to mouse without recalibration.

No cortical circuit motif was imposed in the model. Functional results appearing in the simulations are not the outcome of a dedicated wiring rule: the connectome carries only what follows from morphology, placement and pruning. Axonal targeting is the only constraint with a laminar target, guiding where axons go rather than which cells they contact.

### Thalamocortical projections

The model includes two thalamocortical nuclei built as digital projections: somatotopic VPM and POm. Both the anatomy of the fibres and their strength were constrained by mouse data. VPM fibres were placed within individual barrel outlines and POm across the barrel field, septum included (Figure 4 A, C). A digital fibre carries only a depth profile of synapse density, with no branching geometry. Several features of the projections were constrained from measurement. Depth profiles of synapse density (Figure 4A) follow the fluorescence measurements of El-Boustani et al. (2020), scaled to absolute density using the electron microscopy of Bopp et al. (2017). Lateral spread was set from the extent of thalamocortical axons in CCFv3, projected onto the flatmap. Postsynaptic potential amplitudes were fitted per pathway to Sermet et al. (2019), so Pvalb receiving larger potentials than excitatory cells is a target rather than an outcome (Figure 4E). Short-term dynamics were set from rat measurements, so the depressing-dominant family assignment (Figure 4B) is an input to the model rather than a finding.

No rule assigns thalamic synapses to particular cell types, so what each neuron receives depends on where its dendrites reach. The number of synapses per connected pair is highest onto layer 5 excitatory cells in both pathways, and for VPM also onto layer 4 and layer 6 excitatory cells (Figure 4B, left). VPM forms more synapses per connected pair than POm and carries the longer tail (Figure 4D). As in the cortical connectome, nothing acts at the level of molecular class. The amplitude recipe constrains the relative strength of thalamic drive across classes, while its absolute scale rests on a single unitary reference from rat (Bruno, 2006). Fibre counts carry their own assumptions: VPM extrapolated from rat on a conserved fibre-to-whisker ratio and POm scaled from the rat model. The projections are the one part of the model where strong structure was imposed. VPM fibres are confined to barrel outlines and POm spread across the field, so any result that turns on thalamic drive is constrained in a way the cortical connectome is not.

## Experiments

We simulated three experimental contexts (Figure 1D): the quiet and whisking brain states, and active touch of the C2 whisker. Simulations were run on a subvolume of approximately 130,000 neurons. Analyses use the 3 3 grid of barrel columns (rows B, C and D by arcs 1, 2 and 3, Figure 1B) and the associated septum, approximately 55,000 neurons, with the central C2 column as the principal column throughout and C1 barrel column as the reported neighbour. The grid allows the touch response to be followed beyond the principal column (Figure 6C). The touch input enters through C2 barrel column (Figure 6A), so the response in C1 column is driven principally by corticocortical activation. In C1 column, the excitatory population responds mainly below threshold with a limited suprathreshold response, while the inhibitory classes spike, Pvalb and Vip most strongly (Figure 6B, D, Supplementary Figure S7A,B).

Previous functional simulations of the region were restricted to one cell type or to simplified neurons: the transformation of synaptic input into spike output in layer 5 pyramidal tract neurons responding to passive whisker deflection in rat (Bast et al., 2024), and touch representation in the granular and supragranular layers (Huang et al., 2022). The present model simulates all layers and molecular classes of mouse barrel cortex together.

The model reproduces three patterns reported in mouse (Kiritani et al., 2023; Yu et al., 2019). The touch response in C2 column follows a laminar sequence, starting in layer 4 and propagating into layer 2/3 and then into the deeper layers 5 and 6 (Figure 6B, H). The sequence does not survive beyond the principal column and the septum (Figure 6C, H). The molecular classes follow the sequence: Pvalb then EXC then Sst, above and below threshold (Figure 6D, E). Firing and recruitment change with state (Figure 5E, F; Supplementary Figure S4C), and Sst hyperpolarises at whisking onset while the other classes depolarise (Figure 5G). None of these patterns was fitted.

Cell densities, morphologies, synaptic physiology and the thalamocortical projection were each constrained on their own data. Background conductance was calibrated to reported spontaneous rates, but no target was set for the laminar sequence, the class order or the class-specific response to the change of state.

### Sst hyperpolarisation without the canonical motif

One class responds to whisking onset differently from the rest. Pvalb, Vip and EXC depolarise throughout, while Sst depolarises briefly before turning to a weak hyperpolarisation (0.18 0.10 mV, mean SEM across 5 seeds; Figure 5G; Supplementary Figure S4E). Gentet et al. (2012) and Pala and Petersen (2018) report the same divergence independently, with layer 2/3 Pvalb depolarising and Sst hyperpolarising during active states. The effect is present in layers 2/3 to 5, extending deeper than reported by Kiritani et al. (2023), who observe the hyperpolarisation in superficial but not deep neurons.

At onset every cell receives the same relative step in background conductance, and the thalamic drive rises across the population, so a class-specific response has to come from how that drive is distributed by the circuit. Sst neurons are only weakly driven by the thalamus (Figure 4E), unlike EXC and Pvalb, whose depolarisation follows the thalamocortical input directly. In vivo, the hyperpolarisation is attributed to the excitation of Vip cells, which innervate Sst strongly (Kiritani et al., 2023). No inhibitory-to-inhibitory rule was imposed here, and Vip is not the dominant source of the inhibition Sst receives. Sst is the only major class whose inhibitory drive exceeds its excitatory drive, and most of that inhibition comes from Pvalb (Supplementary Figure S3E). The model therefore reproduces the reported hyperpolarisation through a different source of inhibition, one that follows from the connectivity as built rather than from an imposed Vip-to-Sst motif. This does not exclude the Vip route in vivo, but it shows the route is not required for the divergence.

The transient hyperpolarisation of excitatory cells that Kiritani et al. (2023) report at whisking onset, and leave unexplained, is not present in the model (Figure 5G). Behaviour reverses during the touch response. Sst is recruited across layers 2/3 to 5, with a delay (Figure 6D,E). The inhibitory dominance in its input is unchanged, but the excitatory cortical response to touch is far larger than to whisking (Figures 5F and 6E), and Sst in C2 barrel column depolarises with the rest of the circuit (Figure 6D).

### Whisking phase locking

During whisking, the membrane potential of layer 2/3 neurons tracks whisker position (Crochet & Petersen, 2006; Pala & Petersen, 2018). The whisking cycle is encoded in the VPM input (Figure 5A, Supplementary Figure S5, Urbain et al., 2015), and it is represented in the cortical membrane potential: close to half the cells in C2 column lock to the rhythm below threshold, with every major molecular class represented (Figure 5H).

VPM is the only rhythmic input in the model: the background conductance is held constant during whisking and POm carries no phase preference. Locking therefore requires VPM, and it survives disconnection of the cortex (Supplementary Figure S6E), so the connected circuit modulates its amplitude rather than creating it. Modulation amplitude is largest in Pvalb and Vip (Figure 5H), whereas Kiritani et al. (2023) report the opposite ranking, with the largest amplitudes in excitatory and Pvalb cells and the smallest in Vip. Whisking-related input from other cortical areas and from POm is absent from the model, and may account for the difference. The preferred phase is class-specific (Supplementary Figure S6G), with excitatory cells of layers 4 and 5 concentrating on a preferred phase while those of layers 2/3 and 6 spread more widely.

### Vip is driven through cortex

During whisking, Vip cells are recruited more broadly than any other class (Supplementary Figure S4C), as reported in vivo (Yu et al., 2019). At touch, the Vip response arrives after the excitatory and Pvalb responses (Figure 6E), as Kiritani et al. (2023) report. They attribute this to weak thalamic input to Vip, which agrees with the cortical route reported above. The delay is shorter in the model and the response is larger (Figure 6D,E), which may reflect the absence of long-range cortical drive, neuromodulation and other inputs the model does not contain.

### Columnar and horizontal components of the touch response

The cyclic decomposition (Figure 6F) returns two orthogonal modes with distinct spatial profiles: one focused on the principal column, the other crossing the septum and the neighbouring barrels in laminar bands concentrated in layers 4 and 6 (Figure 6G, H). The phase gradients differ in direction (Figure 6 I): CyC1 advances through the depth of the column, while CyC2 advances horizontally across the barrel field. CyC2 is the smallest of the two, roughly a third of CyC1 in amplitude and a quarter of its contribution to the reconstruction (Supplementary Figure S8D). It is also the slowest and the repeating one, peaking at 20 ms against 14 ms for CyC1 and passing through similar states twice within the response.

Orthogonality is a property of the planes, not of the cells. Cells participate in CyC1 alone, in CyC2 alone, or in both, and the strongest participants in the two overlap substantially (Supplementary Figure S8A, B). The two modes therefore separate the response dynamics, not the population.

### Anatomical predictors of the two modes

CyC1 participation is far better predicted than CyC2 (full-model *R*^2^ 0.53 0.14 against 0.22 0.10, mean SD across the eight cell classes; Figure 6J). Within CyC1, prediction weakens with depth in the excitatory classes, so the columnar mode is best explained where the thalamocortical input arrives in layers 4 and 2/3. CyC2 is poorly explained by connectivity, and what prediction there is rests mostly on cortico-cortical excitatory drive (Figure 6J). Participation in the horizontal mode is therefore not a fixed property of a cell but depends on the activity it sits in. A related distinction appears in Bast et al. (2024), where the trial-to-trial and cell-to-cell variability of layer 5 pyramidal tract responses is set by the synaptic input arriving from the network rather than by a cell’s morphology or biophysics.

### Properties of the selected populations

We selected three groups of cells on single-cell criteria: reliable touch reporting (Figure 7A), and coupling strength and delay during whisking (Bimbard et al., 2026). Touch responders receive slightly more VPM synapses than non-responders (276 against 240, Figure 7E), a small difference given that these cells were selected on their response to a thalamic input.

Choristers, selected from whisking spiking alone with no reference to touch or connectivity, stand well above the soloists (382 against 226 VPM synapses, Figure 7E). The two populations partly overlap, most of all in layer 4 excitatory cells, and both sit at high radius and early phase in the two cyclic components (Figure 7C). Many of the same cells therefore carry a high median thalamic input (Figure 7E), the tightest coupling to ongoing activity, and the earliest participation in the touch response (Figure 7C). Population coupling is an in vivo observable with no measured anatomical correlate. Spiking and afferent connectivity are known for the same cells here, so the model supplies a candidate one.

Leaders share none of these properties. Their VPM count sits only marginally above the soloists (254 against 226), their combined response magnitude falls below them, they share almost no cells with the responders, and they never co-occur with the choristers. Coupling delay is also the least reliable of the three groups: computed separately on two halves of the data, the per-cell values agree only weakly.

### Contributions of dendritic polarisation

The functional consequence of dendritic polarity in layer 4 has not been studied. We therefore built a second model without this feature to quantify it (Figure 8A).

Removing the constraint redistributes synapses onto layer 4 excitatory cells: in POL, layer 4 spiny stellate cells of C2 barrel receive more cortical input from C2 column and fewer from the septum and the surrounding barrels, and more synapses from C2 VPM fibres (Figure 8B). The suprathreshold activity of the circuits diverges from the onset (Figure 8C). Initially POL spikes remain within the principal column (10 ms). The subtracted signal then turns negative, indicating the neighbouring barrels are more active in SYM (15 ms onwards). In C2 column the activity forms a laminar response, with a positive band followed by a negative one at the same depth, indicating that SYM lags behind POL. From spiking activity alone, we see amplitude that moves rather than one that grows, but subtraction alone cannot separate amplitude from timing. Further investigation through decomposition divides the signal into common and contrast planes (Figure 8D). In the common plane the two activities nearly coincide, with the same spatiotemporal structure. This activity carries most of the response in both circuits and the dendritic constraint leaves it unchanged. The contrast plane highlights the individual contributions of the polarity. POL-unique activity is early and stays within C2 column and the septum, running from layer 2/3 and layer 4 downwards between 7 and 15 ms, with negative signal in the neighbouring barrels. SYM-unique activity is delayed, starts in layer 4 of C2, and covers the grid by 25 to 30 ms, including layer 4 of the neighbours rather than avoiding it. Contrast peaks at 12 ms in POL against 18 ms in SYM. Polarisation sharpens the response in space and advances it in time.

For both circuits the selective cells sit throughout the grid (Figure 8G). POL-preferring cells are largely Pvalb and Vip in layers 2/3 and 5; SYM-preferring cells are largely layer 4 excitatory cells outside the principal column. Polarisation advances layer 2/3 and layer 5 Pvalb and Vip ahead of the layer 4 excitation that would otherwise carry the response across the barrel field (Figure 8H). A measure built on single-cell reliability points the same way: among layer 4 excitatory cells, more qualify as touch responders only in POL than only in SYM inside C2 barrel column (230 against 195), and the reverse holds in the surrounding barrels (98 against 135).

### Limitations

The model inherits its biophysics from rat, while mouse data constrain its anatomy and the scale of its synaptic and thalamic drive. Synaptic kinetics and short-term dynamics come from rat paired recordings and are used unchanged in mouse. The e-models were fitted to rat whole-cell recordings and exhibit a strong hyperpolarisation after an action potential, which contributes to the membrane potential variance in both the quiet and whisking states. We therefore did not find the expected larger within-epoch fluctuations of membrane potential in the quiet state, where up and down states alternate (Kiritani et al., 2023): the standard deviation of the membrane potential, averaged across classes, is comparable in the two states, 2.9 mV in quiet against 3.5 mV during whisking (Supplementary Figure S4B). Mouse whole-cell and paired recordings would allow the e-models, synaptic kinetics and short-term dynamics to be refitted for mouse, improving the biophysical fidelity of the model.

The polarity measurements rest on reconstructions from different laboratories, protocols and animal ages, with small and uneven per-class counts. The sample covers a subset of the morphological types in the model. Only layer 4 spiny stellate cells were synthesised entirely from these reconstructions, the other constrained classes combine them with the adapted rat library, rescaled to mouse cortical thickness. Layer 4 untufted pyramidal cells were constrained on prior reports (Staiger, 2004) rather than on our sample, where they showed no enrichment (*p* = 0.62). A dedicated dendritic reconstruction dataset from mouse barrel cortex, acquired under a single protocol and covering all morphological types, would turn the per-class fractions into population statistics, and can be run through the same measurement unchanged.

The connectome is built from anatomy alone, with no rule imposing how likely one cell type is to connect to another. Cortex carries more specificity than apposition and pruning of proximal morphologies alone can reproduce (Staiger & Petersen, 2021). Motifs that depend on it are not reproduced: Vip, which innervates Sst strongly in vivo (Kiritani et al., 2023), is not the dominant source of the inhibition Sst receives in the model (Supplementary Figure S3E). The class-specific dynamics reported here therefore arise from anatomy alone, a baseline on which added specificity can be tested.

### Conclusions

We presented a mouse barrel cortex model, built with an established rat pipeline. We integrated mouse data where applicable and ran three in-silico experiments. Class-specific dynamics appear without class-specific wiring. The model has no imposed rules to build a canonical structure of molecular classes. Connectivity follows the morphological shape and placement rules instead. The Sst hyperpolarisation at whisking onset and the late Vip response to touch are present nonetheless. In the model the Sst hyperpolarisation is driven by inhibitory input, dominated by Pvalb.

We studied the neurons of the principal barrel column under two conditions: whisking and active touch. We selected the cells by their coupling to spontaneous activity and by the strength and reliability of their touch response. Choristers and touch responders identified under these conditions turned out to be largely the same layer 4 excitatory cells. These cells carry high VPM input, but not exclusively.

After cyclic decomposition, we found two main modes of activity: columnar (CyC1) and horizontal (CyC2). Participation in the columnar mode is better predicted from static connectivity than participation in the horizontal mode. This suggests that participation in the horizontal mode depends on the input a cell receives during the response.

Finally, we compared two circuits differing in one anatomical property. Polarised basal dendritic trees in layer 4 of the barrels have a direct effect on the touch response. Through decomposition we found the common activity shared between the two circuits and contrast activity, presenting unique contributions. This revealed that polarisation sharpens the touch response in space and advances it in time, acting as a spatial filter of the response.

## Methods

The Methods follow the order of the work. The model was built from atlas. It was simulated under a thalamic drive reproducing the quiet and whisking states, and the resulting activity analysed by the methods below.

### Atlas and geometry

The model is built in the Allen Mouse Brain Common Coordinate Framework at 25 *μ*m resolution (Wang et al., 2020), with the barrel field enhancement of Bolaños-Puchet, Teska, et al. (2024). The enhancement supplies the barrel and septum annotations, the laminar depth coordinate, the per-voxel orientation vectors, the flatmap and the barrel column meshes, together with the software used to build and query them. Cell density volumes are from Keller et al. (2018), which also supplies the thalamic densities combined with CCFv3 nucleus volumes in Section *Thalamic nucleus size*.

### Cell composition

Densities and me-type composition were taken from the rat somatosensory cortex model (Reimann et al., 2024), which draws total neuron and glia densities per region from Keller et al. (2018) and distributes the inhibitory populations within them according to the brain-wide maps of Kim et al. (2017). The resulting me-type and e-type composition is given in Supplementary Figure S1C, D, and the per-type counts of the C2 barrel column in Supplementary Figure S1E, F. Neurons are placed voxel by voxel from the density recipe, each soma inheriting the orientation vector of its voxel.

Four mouse-specific refinements were applied. Layer 4 spiny stellate cells were set to 76% of excitatory neurons and layer 4 Sst neurons to 96% non-Martinotti (Scala et al., 2019). Layer 4 Vip and Sst densities were set separately for the barrel and septum compartments following Almási et al. (2019): Sst 2,661 against 4,773 and Vip 1,575 against 3,205 cells per mm^3^. No other compartment distinction was applied. The layer 5 Sst depth profile was refined following Tremblay et al. (2016), whose layer 4 Pvalb peak the profiles do not reproduce.

### Molecular identity

The mapping of Roussel et al. (2023) was obtained from the authors as a distribution over the four markers per me-type (Supplementary Figure S1B). Every inhibitory neuron drew its marker independently from the distribution of its me-type.

### Dendritic polarity

#### Angular distributions

Basal dendritic segments were reduced to soma-centred midpoints; each contributed an angle *θ* = atan2 *y*, *x* weighted by segment length. Reconstructions with too few resolvable branches were excluded. Each distribution was rotated to a common mean direction before feature extraction.

#### Von Mises fitting

Each distribution was fitted with a von Mises, the circular analogue of a Gaussian, with mean direction *μ* and concentration *κ*:

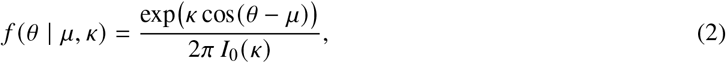

where *I*_0_ is the modified Bessel function of the first kind of order zero. The resultant length *R* is the magnitude of the length-weighted sum of unit vectors at each segment angle, divided by total dendritic length. The expected resultant length of a von Mises is *A κ* = *I*_1_ *κ I*_0_ *κ*, which increases monotonically, so *κ* was obtained by inverting *A* at the measured *R* (Fisher approximation, pycircstat); *μ* was the length-weighted circular mean.

#### Mixture model

Each distribution was additionally fit with a two-component von Mises mixture (*K* = 2 throughout),

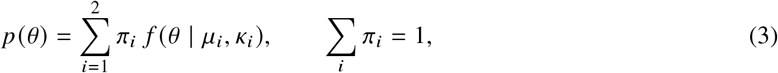

by expectation-maximisation from random initial weights, mean directions and concentrations. Component concentrations were updated by numerical inversion of *A* at the responsibility-weighted resultant length. Iteration stopped when the summed squared change in *π*, *μ*, *κ* fell below 5 10^−4^. A component that lost all mass triggered re-initialisation, with the tolerance relaxed if re-initialisation recurred. Fitting ran under a fixed random seed.

#### Polarity descriptors

Five descriptors were taken per reconstruction. The single-fit concentration *κ* comes from Eq. 2. From the mixture, the two components were ordered by dominant mass *πκ*, giving the primary concentration *κ*_primary_ = *κ*_1_, its weight *w*_1_ = *π*_1_, and the angular separation Δ*μ* = *μ*_1_ *μ*_2_ wrapped to 0, *π*, which is largest for two opposed lobes. Overall directionality was summarised by the effective resultant length,

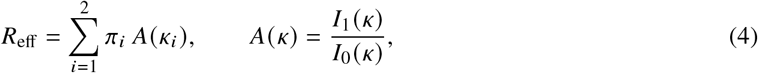

the resultant length each component would have averaged by weight. All five were standardised across reconstructions before combination, and the composite score was z-scored again for display.

#### Clustering

The standardised descriptors were reduced by PCA to two components. Cluster number was evaluated over *k* = 2-10 by silhouette, Calinski-Harabasz and Davies-Bouldin score; *k* = 2 was selected. Final labels came from agglomerative clustering (Ward linkage) on the PCA scores, with the cluster of higher median *κ*_primary_ designated polar. UMAP (n_neighbors = 30, min_dist = 0.6) on the same PCA scores was used for two-dimensional visualisation only. PCA, UMAP and clustering used a fixed random seed.

#### Enrichment

Enrichment of polar cells in each morphological class was tested against the pooled remaining classes by one-sided Fisher exact test on the 2 2 table of polar and non-polar counts; class labels come from the morphology database and not from the clustering. Per-class polar fractions are given with Wilson 95% confidence intervals and the number of reconstructions per class, alongside the pooled layer-4 versus non-layer-4 contrast. Reported p-values are uncorrected; under Benjamini-Hochberg correction across the eleven classes the three enriched layer-4 classes remain significant (*q* = 0.007, 0.012 and 0.038 for L4 SSC, L4 NMC and L4 Pvalb), and no other class reaches *q* < 0.05. As a control on the cluster boundary, the polar set was redefined as the top 30, 25 and 20% of the composite score and the test repeated (two-sided), recovering the same three classes at each threshold.

### Morphologies

#### Morphology sets

The dendritic polarity analysis (Section *Dendritic polarity across morphological classes*) drew on 324 mouse reconstructions; synthesis drew on the 100 curated mouse reconstructions, with per-m-type counts for both the dendrite and axon pools in Table S2. The two sets overlap in 92 reconstructions, listed by morphological type and source archive in Table S3. The remaining 8 were taken from the MouseLight dataset (Winnubst et al., 2019).

#### Topological synthesis

Dendrites were generated by topological synthesis (Kanari et al., 2019, 2022), oriented in the atlas by the local principal direction and cortical depth of the soma. Input statistics came from the curated release, repaired and corrected for tracing artefacts, pooled per m-type across 1,204 reconstructions spanning the 60 m-types (median 18 per m-type, range 1 to 82). Sample sizes are uneven, and 14 m-types contributed fewer than five reconstructions each, so synthesised trees for those types rest on correspondingly narrow input distributions; the four classes carrying the barrel constraint had 21 neurons (L4 SSC), 7 neurons (L4 NMC), 15 neurons (L4 Pvalb) and 36 neurons (L4 UPC). Every cell was grown from an independently jittered barcode, with rotational jitter of standard deviation 10^◦^ and scaling jitter of standard deviation 0.2, under a fixed random seed. Cells whose synthesis failed were dropped, with a tolerated drop ratio of 0.1.

#### Boundary repulsion

For polarised neurons each candidate growth step was accepted with probability

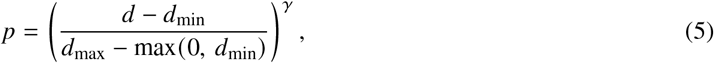

for *d* the distance of the step from the mesh wall, clipped to 0, 1 . The barrel-column meshes were the containing boundary, so each cell was constrained to the barrel its soma occupies. The four constrained classes map to synthesis m-types L4_SSC, L4_BTC (L4 NMC), L4_SBC (L4 Pvalb) and L4_UPC. L4 SSC and L4 NMC used *γ* = 1.3, L4 Pvalb and L4 UPC *γ* = 1.25; all four used *d*_min_ = 5 *μ*m and *d*_max_ = 40 *μ*m, so steps were rejected outright 5 *μ*m beyond the wall and accepted freely from 35 *μ*m inside it. Axons, apical dendrites and all other m-types were synthesised without repulsion.

#### Dendrite containment

Containment was measured per cell as the fraction of basal dendritic length lying inside the outline of the cell’s own barrel column. Each dendritic segment was projected onto the flatmap and tested against the barrel outline, then weighted by its length. The measure was applied to the four constrained classes in the C2 barrel column, in both the polarised and the symmetric circuit.

#### Axons

Each neuron received an axon grafted from a reconstruction of its own m-type, chosen by scoring candidates against placement rules and selecting the best-scoring candidate for that soma position (Reimann et al., 2024). Rules specify the layer interval an axon should target, together with hard limits keeping dendrites below the pia and axons within 40 *μ*m of it, and a bias term favouring candidates that need little rescaling.

#### Laminar placement rules

Axons and apical dendrites are placed by laminar targeting rules defined per morphological type, inherited unchanged from the rat model (Reimann et al., 2024). Each rule gives a target interval in cortical depth, bounded by a layer and a fractional position within it. Intervals are expressed in the atlas layer coordinate, so one rule applies at every position of the barrel field. Two global rules apply to all types: dendrites must remain below the pia, and axons may exceed it by at most 40 *μ*m. Candidates are further weighted by a bias favouring those that need little rescaling (*σ* = 3.5). Each axon was rotated by an angle drawn uniformly on [−*π*, *π*] about the vertical axis. The full rule set is given in Table 1.

#### Morphology dataset

The mouse morphologies used here are published with the model. The dataset comprises the 100 curated mouse barrel cortex reconstructions and the full set of synthesised dendritic trees, one per neuron. All 100 contributed dendritic barcodes to synthesis and 80 contributed axons to the grafting pool, which spans the full curated release rather than the mouse cells alone (Table S2). Each was assigned to an existing morphological type, and no new type was added to the scheme.

### Electrophysiological models

Each e-model was paired to a synthesised morphology by the workflow of Arnaudon et al. (2023), which rescales the soma and axon initial segment until two input resistance ratios match the target for its e-type: *ρ*, somatic to dendritic, and *ρ*_axon_, somatic to axon initial segment. Ratios were computed per e-model on its canonical morphology, *ρ* ranging 4.0 to 14.8 and *ρ*_axon_ ranging 33.5 to 237.1.

Evaluations ran at 34 ^◦^C from *v*_init_ = 80 mV, with holding current set to 85 mV and a step from 500 to 2000 ms. Threshold current was found by recursive search over 0 to 0.2 nA, depth 10, with spikes detected at the axon initial segment and at most 2 spikes at threshold. Stochastic channels were run deterministically with a conductance correction of 0.9. A pairing entered the circuit only if every electrophysiological feature fell within five standard deviations of the experimental mean.

### Connectome

#### Appositions and pruning

Appositions were detected with TouchDetector (Reimann et al., 2015) at a maximum touch distance of 2.5 *μ*m for excitatory and 0.5 *μ*m for inhibitory presynaptic axons, and kept only where the postsynaptic compartment can form that type of synapse. Pruning followed the three steps of Reimann et al. (2015).

The three pruning parameters follow from the mean and standard deviation of synapses per connection (*S_m_*, *S_sd_*) and the bouton density (Reimann et al., 2015). Targets came from barrel cortex measurements of synapses per connection and of bouton density, the latter converted assuming one synapse per bouton. Only the prediction of *S_m_* from the mean number of appositions per connection *n* was changed, to

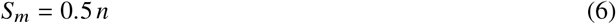

for every pathway, in place of 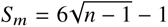 for most pathways and *S_m_* = 1.5 *n* for excitatory to excitatory. *S_m_* was capped at 25, and *S_sd_* was set from a coefficient of variation of 0.32 across all pathways. Chandelier cell pathways were exempted from pruning, since their axons target the axon initial segment and the apposition counts are not comparable.

#### Connectivity recipe

The connectivity recipe was taken unchanged from the rat somatosensory cortex model (Reimann et al., 2024), apart from the apposition-to-synapse relation above.

Inter-bouton interval was drawn between 5 and 7 *μ*m, with a 5 *μ*m gap at region boundaries. Spine length was 2.5 *μ*m for most morphological types and 0.5 *μ*m for the bipolar, double bouquet and neurogliaform types. Minimum distance from the postsynaptic soma was 25 *μ*m for excitatory and 5 *μ*m for inhibitory synapses, applied wherever no more specific rule applies.

Excitatory axons contact dendrites of every type and the somata of inhibitory types. Inhibitory axons contact dendrites of every type; basket and chandelier types additionally contact pyramidal somata, whereas bitufted, bipolar, double bouquet, Martinotti and neurogliaform types do not. Appositions not matched by a rule were discarded, and chandelier synapses were repositioned to the axon initial segment.

Synapse kinetics, short-term dynamics and peak conductances were assigned per pathway by morphological and electrical type, from 41 synapse classes (30 excitatory, 11 inhibitory).

#### PSP calibration

Peak conductances were calibrated per pathway against the amplitude recipe (Supplementary Figure S3A, B, Table S4), following the implementation of Reimann et al. (2024) with psp-validation. Pathways were defined by presynaptic and postsynaptic morphological class, giving 183 pathways, each with its own target amplitude and holding potential. For each pathway, 80 connected pairs were sampled from the C1 barrel column and each pair simulated 20 times: the postsynaptic cell was held at the holding potential of that pathway, sodium channels left intact, a presynaptic spike evoked at 800 ms, and the somatic potential recorded at 0.1 ms resolution for the following 100 ms. Simulations ran in CoreNEURON with a timestep of 0.025 ms under a fixed random seed, with GABA rise times randomised per synapse. The mean amplitude across pairs and repetitions was compared to the target and the peak conductance rescaled accordingly, scaled to reference over five iterations until simulated amplitudes matched the recipe (Supplementary Figure S3 C). Calibration was run separately for the two circuits, so the conductances differ between POL and SYM while both reproduce the same amplitude recipe.

### Thalamocortical projections

#### Thalamic nucleus size

Combining reported neuron densities (Keller et al., 2018) (VPM 83,100 mm^-3^, PO 41,477 mm^-3^) with the corresponding atlas volumes (VPM 0.82 mm^3^, PO 0.62 mm^3^) gives 68,142 VPM and 25,716 PO neurons. CCFv3 does not subdivide PO, so we assigned 40% of it to POm (10,286) in the absence of an annotated subvolume.

#### Fibre placement

Fibres were placed on the flatmap by Poisson disc sampling. The number of fibres per region was fixed in advance, so we set the spacing to fit it: a larger disc radius admits fewer points, and we searched by bisection for the largest radius that still fits the required number inside the convex hull of the region outline (tolerance 10^−5^ in flatmap units). VPM fibres were sampled within each barrel outline separately, POm fibres within the hull of the full SSp-bfd flatmap. Each sampled point was assigned the local principal direction at the depth of layer 4 and extended below layer 6, so that every fibre crosses the full cortical depth.

#### Synapse density

Fluorescence gives no absolute count, so we set the scale from the electron microscopy of Bopp et al. (2017) in mouse S1, who report 0.225 VGluT2-positive synapses per *μ*m^3^ in L4. Since a fibre is defined by its boutons rather than its synapses, we converted using their mean of 2.1 synapses per thalamocortical bouton, giving 1.07 10^8^ boutons per mm^3^ at the L4 peak. Scaling the remainder of each profile by the normalised fluorescence then fixes the bouton count along a fibre.

This density exceeds the 4.4 10^7^ boutons per mm^3^ counted in rat by Meyer et al. (2010), consistent with the uncorrected 11% linear shrinkage in the electron microscopy and with VGluT2 labelling the full thalamic input to L4. Starting from synapse density also avoids equating each bouton seen under the light microscope with a single synapse, which under-counts thalamocortical synapses in mouse S1 L4 roughly twofold (Bopp et al., 2017).

#### Lateral spread

Thalamocortical axon reconstructions were projected onto the flatmap and, for the points falling in layers 3 to 5, a covariance ellipse at two standard deviations was fitted per axon. Its radii were scaled by the nominal flatmap extent of 3570 *μ*m to convert from flatmap to metric units. VPM axons gave a mean radius of 78.7 *μ*m (82.0 27.7 *μ*m and 75.5 31.0 *μ*m along the two axes) and POm axons 108.5 *μ*m (114.2 58.3 *μ*m and 102.8 56.6 *μ*m). Rounded down to the nearest 5 *μ*m, these set the width of the lateral fall-off used in synapse assignment.

#### Synapse assignment

Synapses were drawn along each fibre from its depth profile with Projectionizer (v3.0.0.dev0). Each voxel considered its closest fibres, 50 for VPM and 100 for POm, and assigned each synapse to one of them with probability falling off as a Gaussian in lateral distance, with standard deviation 75 *μ*m for VPM and 105 *μ*m for POm.

#### Thalamocortical synaptic physiology

Thalamocortical fibres carry the cortical axonal conduction velocity of 300 *μ*m/ms, with a transmitter release delay of 0.1 ms.

Peak conductances, short-term dynamics and release probabilities were taken from the thalamocortical recipes of the rat somatosensory cortex model (Reimann et al., 2024), which draw on the paired recordings of Bruno (2006), assigned per postsynaptic morphological and electrical type, and served as the starting point for calibration. VPM conductances are specified per layer band: 1.1 nS onto layer 4 excitatory cells, 2.36 nS onto layer 5 and layer 6 excitatory cells, and 1.73 nS onto layer 2/3. POm carries a single excitatory value of 1.73 nS for all layers. Basket and chandelier types receive higher conductances than excitatory cells throughout, 1.35, 3.13 and 2.24 nS for VPM across the same three bands and 2.24 nS for POm, following the fast-spiking measurements of Beierlein et al. (2003) and Beierlein and Connors (2002).

The Martinotti, bipolar, double bouquet and bitufted types receive facilitating synapses in both pathways, with a peak conductance of 0.17 nS. All remaining inhibitory targets are depressing. Release probability for thalamocortical excitatory synapses is 0.75, higher than the intracortical value, following Gil et al. (1997).

#### Thalamocortical PSP calibration

Calibration followed the cortical procedure, run separately for the VPM and POm projections, with 80 pairs per pathway and 50 repetitions and each postsynaptic neuron used at most once. Pathways were defined by postsynaptic morphological class, with the presynaptic side left as the whole projection. The postsynaptic cell was held at the holding potential of that pathway, a single fibre spike evoked at 800 ms, and the somatic potential recorded to 900 ms at 0.1 ms resolution, with sodium channels intact and extracellular calcium at 2 mM. Targets were the normalised amplitudes of Sermet et al. (2019), reported per postsynaptic layer and molecular class. Their values are normalised to layer 4 excitatory cells for VPM and to layer 5A excitatory cells for POm, and we set that reference to 1 mV (Section *Thalamocortical synaptic physiology*). Calibration was run separately for POL and SYM, so thalamocortical conductances differ between the circuits while both reproduce the target amplitudes.

### Common neighbour bias

Common-neighbour bias was tested in layer 4 only, on the excitatory and Pvalb populations of the C2 barrel column. We took every pair of layer 4 neurons in C2 barrel column lying within 150 *μ*m of each other, and for each pair counted how many other layer 4 neurons of C2 connect to both members. Pairs were sorted by this count and split into eight groups of equal size, from those sharing the fewest partners to those sharing the most. Connection probability was computed within each group and normalised to the population mean.

Connection probability falls with distance, and neurons close together share more partners, so the rise could follow from distance alone. To control for this, we compared against a null that keeps the distance dependence of connectivity but removes any other structure, so that any rise in the null is due to distance alone. Pairs were grouped by the distance between their somata, and connections were shuffled at random within each distance group. This holds the total number of connections and their dependence on distance fixed, while breaking any relationship between being connected and sharing partners. The width of the distance groups was halved until a two-sample Kolmogorov-Smirnov test no longer distinguished the distribution of connected-pair distances in the null from the observed one.

### Structural synaptic drive

We weighted each synapse by its peak conductance and its utilisation of synaptic efficacy, the release probability at rest. Their product estimates the drive delivered by one presynaptic spike, before any activity is simulated.

For each postsynaptic cell in the C2 barrel column, we summed these weights over its inputs from within the column, grouped by the molecular class of the presynaptic cell, and divided by the cell’s total to give the fraction of drive received from each class. Classes with fewer than 15 cells were excluded. Excitatory and inhibitory weights therefore lie on different scales, and the fractions are comparable across postsynaptic classes rather than absolute.

### Simulation setup

#### Simulated volume

Simulations used the 3 3 grid together with the surrounding tissue, defined as the union of nine cylinders of radius 365 *μ*m about the axis of each barrel, intersected with the SSp-bfd annotation of the left hemisphere. The radius is the largest barrel radius across the field plus a padding of 250 *μ*m against subvolume edge effects (Isbister et al., 2024; Reimann et al., 2024), scaled from rat to mouse (Bolaños-Puchet, Teska, et al., 2024). The node set contains 130,310 neurons.

#### Analysis volume

Analyses of the population response used the nine barrel annotations, all layers, together with the septal neurons lying within 180 *μ*m of the flatmap centroid of C2, giving 56,426 neurons. The volume is defined in the flatmap, not as the simulated volume at a smaller radius. The C2 barrel column contains 5,207 neurons and C1 5,835. Analyses of the brain states and whisking onset used C2 alone.

#### Circuit comparison

Layer 4 dendrites of the constrained classes were synthesised under the boundary repulsion rule in the polarised circuit (POL) and without it in the symmetric circuit (SYM); every other neuron carries the same synthesised tree in both. The connectome was rebuilt for SYM under the same algorithm and parameters, so the connectivity differences follow from the layer 4 dendrites alone. The Ornstein-Uhlenbeck background conductance and the postsynaptic potential amplitudes were drawn independently in each circuit and calibrated by the same procedure against the same targets, the reported spontaneous firing rates and the amplitude recipe respectively. The calibrated values therefore differ between the circuits, while neither the background drive nor the synaptic strengths differ systematically between them. Simulations were matched by seed, so a POL and a SYM run of the same seed received the same thalamocortical input realisation. Comparisons take POL minus SYM; the contrast orientation of Section *Contrast orientation and lead-lag per cell* carries its own sign.

#### Simulator

Simulations were run with Neurodamus (January 2026 release) on NEURON 9.0awith libsonata 0.1.29, at a timestep of 0.025 ms. Extracellular calcium was 1.1 mM in every simulation. Synaptic conductances were calibrated at 2 mM (Section *PSP calibration*) and simulations run at the lower in vivo concentration, so release probabilities during simulation lie below their calibration values.

#### Recording

Spikes were recorded from every simulated neuron. Somatic membrane potential was recorded from every simulated neuron for the full duration of each run, sampled at 0.1 ms.

#### Simulation types

Four types of simulation were run (Table 2). The seed sets the random stream and the sampling of thalamocortical fibres, so the input realisation differs between seeds. The alternating and whisking-onset simulations run both states in 600,ms blocks, ten of each per run, beginning in the quiet state. In the whisking-onset simulations the phase of the background alternation at whisking onset varies between seeds. The whisking-only simulations hold the whisking state throughout. The touch simulations deliver fourteen touches to C2 during whisking, at intervals of approximately 500,ms, jittered by up to 2.9,ms, and were run in both circuits.

**Table 2.**
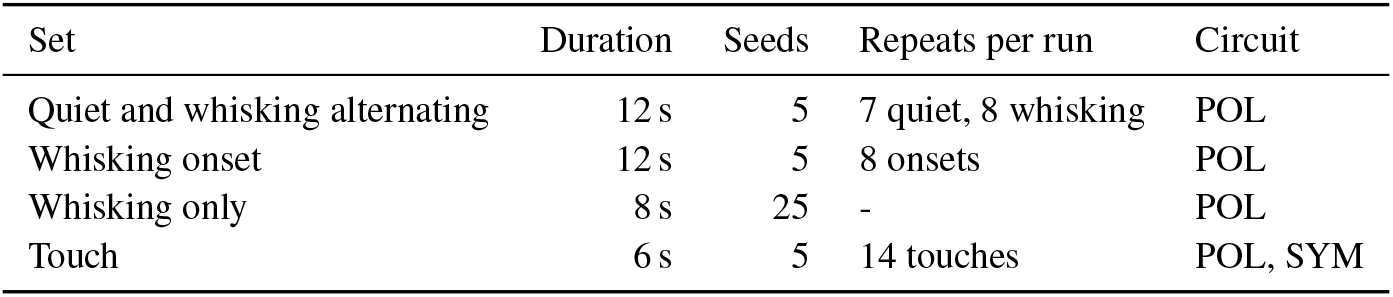
Simulation sets. Repeats are per run and are the counts entering analysis, after the initial transient is discarded.

| Set | Duration | Seeds | Repeats per run | Circuit |
|---|---|---|---|---|
| Quiet and whisking alternating | 12 s | 5 | 7 quiet, 8 whisking | POL |
| Whisking onset | 12 s | 5 | 8 onsets | POL |
| Whisking only | 8 s | 25 | - | POL |
| Touch | 6 s | 5 | 14 touches | POL, SYM |

#### Initial transient

The first 1 s of the alternating and onset runs was discarded, removing the first quiet block and the first whisking block and leaving nine epochs of each state per run. The whisking-only runs discard the same interval. In the touch runs the first 500 ms was discarded and all fourteen touches retained, the first falling at 501.6 ms.

### Background conductance

The simulated volume receives no input from the rest of cortex. That drive was replaced by somatic conductance injection of an Ornstein-Uhlenbeck process per neuron, representing aggregated background synaptic input (Destexhe et al., 2001). Its level was set in three steps, the first two giving a calibrated reference conductance and the third setting the brain state (Supplementary Figure S4).

#### Reference level

First, mean and standard deviation were calibrated by the algorithm of Isbister et al. (2024). Both are set as a percentage of each cell’s input conductance at rest, with their ratio fixed at 0.4 across all neurons. The calibration targets the in vivo reference rates of Table 3 in full rather than a fraction of them. It was run with no thalamocortical input, the projections carrying neither evoked nor spontaneous release, and gives reference values per layer and per excitatory or inhibitory class. Target rates follow Isbister et al. (2024) and were revised against the mouse barrel cortex recordings of Kiritani et al. (2023).

**Table 3.** Target spontaneous firing rates used for the background calibration.

| Layer | Excitatory (Hz) | Inhibitory (Hz) |
|---|---|---|
| L1 | - | 1.50 |
| L2/3 | 0.86 | 12.37 |
| L4 | 2.01 | 13.28 |
| L5 | 3.64 | 2.36 |
| L6 | 0.47 | 14.90 |

Second, within each layer the reference values were scaled per molecular class. The multipliers were selected by a grid search in which candidate values were simulated together with the thalamic input. A candidate was accepted if spontaneous rates fell within the reported range and network-wide bursting was absent. Achieved rates per molecular class are reported in Figure 5F.

#### State scale

Third, the brain state is set by a single scale on the calibrated conductance, with mean and standard deviation scaled together so that their ratio is unchanged across states. During whisking the scale is held at 0.45 throughout, producing a sustained depolarisation. During quiet it alternates at 5 Hz between 0.30, giving the down phase, and 0.30 0.20 = 0.50, giving the up phase, so the whisking level lies between the two. The alternation is in phase with the 5 Hz modulation of the thalamic input, so cortical and thalamic drive rise together into the up phase. This gives explicit control over the state of the circuit. It reproduces up and down alternation and the whisking depolarisation in a sub-volume that cannot generate them on its own.

Calibration was performed separately in each circuit by the same procedure and against the same targets.

### Thalamocortical input generation

VPM and POm input was generated independently per fibre as stochastic point processes, tuned to the firing rates reported for the quiet and whisking states (Urbain et al., 2015). Because the processes are stochastic, the input differs between seeds, and input statistics are reported as averages across fibres (Figure 5B). Half of the available fibres of each nucleus were used, drawn once per run, so the set of active fibres also differs between seeds. 118 VPM fibres project to C2, 59 of them active. 925 POm fibres fall within the simulated grid and its septum, of which 462 are active in a given run. Spike times were generated per fibre from an instantaneous rate *λ t* sampled at 1 ms, over the full run duration (Table 2).

Whisking was imposed at 10 Hz in fixed 600 ms periods alternating with the quiet state, the periods shaded in Figure 5. In a behaving animal, whisking occurs in bouts of variable onset and duration. Fixed periods were used to keep the two states directly comparable.

For the modulated conditions,

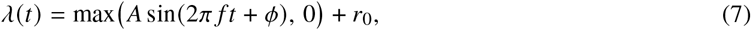

with *φ* the fibre’s preferred phase.

#### VPM, whisking

*f* = 10 Hz, *r*_0_ = 12 Hz, inhomogeneous Poisson process with a 2 ms refractory period. Fibres were split by modulation strength following Urbain et al. (2015): of every ten, two were strongly modulated (*A* = 26 Hz), four weakly modulated (*A* = 10 Hz) and four unmodulated (*λ t* = *r*_0_). Modulated fibres fire preferentially at their preferred phase within the 100 ms cycle (Supplementary Figure S5A, D).

#### POm, whisking

*λ t* = 11.77 Hz, inhomogeneous gamma process of shape factor 1.50, with no phase preference. POm is gamma in whisking and Poisson in quiet, so the fraction of spikes fired in bursts rises with the state.

#### Quiet, VPM and Pom

*f* = 5 Hz, *A* = 10 Hz, *r*_0_ = 3 Hz, *φ* = 0 for every fibre. Inhomogeneous Poisson process with a 2 ms refractory period.

#### Alternation

Quiet and whisking trains were generated for VPM and POm over the full duration and merged in alternating blocks, on the same boundaries in each.

#### Phase assignment

Preferred phases were set by index at generation, *φ* = 90^◦^ 30^◦^ *i* mod 12 for train index *i* within each modulated group, and the resulting trains then permuted onto fibres. Per train, spike times were folded onto the 100 ms cycle in 5 ms bins and fitted with *b a* sin 2*π t c* 100, giving modulation strength *a b* and preferred phase *c* (Figure 5B, right). Each fibre’s angle about the centroid of the flatmap positions of its barrel’s fibres was matched to the fitted phases by linear sum assignment, so that preferred phase rotates around the fibres (Supplementary Figure S5C).

#### Touch

Touch was delivered by VPM alone. The recruited fibres are a random half of the C2 whisking fibres, drawn once per run and used for every touch of that run: 29 of the 59 whisking fibres, a quarter of the projection. They carry the touch spikes in addition to their ongoing whisking input. Spike times were drawn from a lognormal distribution relative to touch onset, *μ* = 1.4963 and *σ* = 0.4733, fitted so that the sampled population response reproduces the VPM stimulus response reported by Yu et al. (2019). A total of 697 spikes was drawn per touch and assigned at random to the recruited fibres, so a fibre carries a variable number of spikes and the times are independent of fibre identity. Each of the fourteen touches of a run is an independent realisation of the spike times and of their assignment. Touches were spaced 500 ms apart, jittered per touch, and delivered in isolation. The interval is long relative to the 100 ms response window, so no touch falls within the response to previous stimuli.

### State epochs and per-state metrics

Both states are sampled within the same run, under the same connectome, input realisation and background calibration. Analysis used 5 seeds of the alternating simulation set and the C2 barrel column throughout. Blocks were assigned to a state by their position in the alternation and taken whole, the first quiet block of each run excluded. Seven quiet and eight whisking epochs enter the analysis, spanning 1200-9000 ms and 600-9600 ms respectively.

#### Epoch definition

We ran simulations in 600 ms blocks, referred to as epochs, which include whisking and quiet epochs.

#### Spike removal

Action potentials were removed from the membrane potential per cell before analysis. Onsets were detected where the rate of change of the trace exceeds 20 mV/ms, with a minimum inter-spike interval of 3 ms; the trace was then held constant from the point at which it rises 5 mV above the onset voltage until it returns below that level. The subthreshold depolarisation preceding each spike is retained.

#### Membrane potential

Per cell and per epoch, the membrane potential was averaged over the samples of that epoch; the per-epoch values were then averaged across epochs and seeds, giving one value per cell per state (Figure 5E). The cell is the unit of replication, so class summaries are taken across cells. The two states were balanced to seven epochs each for this analysis. The standard deviation within an epoch was computed on the same samples and averaged the same way (Supplementary Figure S4B), so for the quiet state it includes the up and down alternation.

#### Firing rate

Spikes were counted per cell over each epoch and divided by its duration, then averaged across epochs and seeds (Figure 5F). Cells that never fire enter as zero rather than being excluded, so class means are over the full class. All available epochs were used, eight for whisking and seven for quiet.

#### Recruitment

Recruitment is the fraction of cells in a class firing at least one spike in a state, computed per seed over all epochs of that state and averaged across seeds, and reported per layer and molecular class (Supplementary Figure S4C). The window is the whole state within a run rather than a single epoch, so a cell firing once in that run counts as recruited.

#### Whisking onset

The whisking onset set was used, nine onsets per run, every 1200 ms from 600 ms. Membrane potential was aligned to onset per cell and baseline-subtracted over the 50 ms preceding it, averaged across onsets within a cell, then across cells within a molecular class, and finally across seeds, with the standard error taken across seeds (Figure 5G, Supplementary Figure S4D). Firing rate was binned at 1 ms, smoothed with a Gaussian kernel of one bin, normalised by the number of cells in the class and baseline-subtracted over the 200 ms preceding onset (Supplementary Figure S4E). L1 Lamp5 was excluded from this analysis.

### Whisking phase locking

Analysis used 25 seeds of the whisking-only simulation set, 8 s each, discarding the first 1 s. Membrane potential was sampled at 10 kHz and action potentials were removed as in Section *State epochs and per-state metrics*. Cells of the C2 barrel column were analysed (*n* = 5,207).

#### Phase clock

Spike times of the VPM fibres innervating C2 were binned at the membrane potential sampling rate and smoothed with a 5 ms boxcar. The rate was mean-subtracted, bandpassed with a third-order Butterworth filter at 7-13 Hz applied forwards and backwards, and the Hilbert transform taken. Instantaneous phase and envelope were the angle and modulus of the analytic signal (instantaneous frequency 9.95 Hz, IQR 1.89 Hz, 67 to 74 cycles per seed); cycle boundaries were the 2*π* crossings of the unwrapped phase, which was then interpolated onto the membrane potential time base. The band was restricted to the fundamental because rectification of the sinusoidal thalamic drive into a firing rate generates harmonics: under the 5-25 Hz band applied to whisker angle by Kiritani et al. (2023), the analytic phase tracked the 20 Hz component in 44% of samples. The commanded 10 Hz sinusoid was not used as the reference, since its stationarity causes residual noise to accumulate in fixed phase bins.

#### Cycle inclusion

All cycles were retained. Kiritani et al. (2023) required whisker amplitude above 15^◦^, selecting the periods in which the animal whisked; the drive here is present throughout, so no cycle-level exclusion applies. Thresholding on envelope amplitude is not a neutral filter: a sweep over the 25th, 50th and 75th percentile of pooled cycle amplitudes raised median modulation amplitude from 0.36 mV to 0.73 mV and shifted preferred phase by a median of 22^◦^, while reducing the cycles available per seed from 70 to 19. Selecting the cycles of strongest drive therefore inflates the amplitude it measures. Across-seed coherence, on which the locking criterion rests, was stable over the same range (0.31-0.33), so the classification of cells as phase-locked does not depend on the choice.

#### Per-cell trajectory

Within each cycle the mean membrane potential was subtracted per cell, removing between-cycle drift. Samples were then pooled across cycles and averaged into 100 phase bins; because bins are defined in phase rather than in time, cycles of differing duration superimpose correctly. Cells with fewer than 80% of bins filled, or fewer than 10 cycles, were excluded. Modulation amplitude and preferred phase were taken from the first-harmonic resultant of the binned trajectory, *V* = *_k_ x_k_ e^iθk^*, equivalent to a least-squares sinusoidal fit, with amplitude reported as peak-to-trough (4 *V n*) and preferred phase as arg *V* . The peak-to-trough range of the raw binned trajectory is 2.4 times larger, since its expected value scales with membrane potential noise.

#### Null and locking criterion

A time-roll null was built per seed by circularly shifting the phase clock against the membrane potential by a random offset of between one tenth and nine tenths of the recording, constrained to be close to antiphase modulo the cycle period. Per-cell metrics were computed from the null as from the real data. Across-seed phase coherence was the resultant length of a cell’s per-seed preferred phases. A cell was called phase-locked if its coherence exceeded the 95th percentile of the null coherence distribution (0.347) and it contributed at least 20 of the 25 seeds; 46.4% of cells met both. Per-cell modulation amplitude is reported relative to that cell’s own null amplitude, since the null floor differs systematically between classes (0.24-0.62 mV). Amplitudes were compared against the matched null per class by Wilcoxon signed-rank test over all cells of that class, gated and ungated cells alike. Phases are referenced to the peak of the C2 VPM firing rate rather than to peak whisker protraction, so absolute values are not directly comparable to experimental ones.

#### Class summaries and error

Amplitude, preferred phase and within-seed phase alignment were computed per seed and averaged, with the standard error taken across the 25 seeds. The error on the fraction of phase-locked cells was estimated by split-half resampling: 50 random partitions of the 25 seeds into two halves of 12, with the null floor recomputed within each half. Two resultant lengths are reported and are not interchangeable: the within-seed value, which measures how aligned the cells of a class are within a single run, and the across-cell value computed on the seed-averaged preferred phases, which is larger because per-cell averaging removes seed-to-seed jitter.

#### Four-condition comparison

One seed was rerun in four configurations, with VPM input present or absent and cortical connectivity intact or removed, and the phase-binned trajectory computed as above against the clock of the VPM-present connected run. Population means are the mean across cells with the standard error across cells, and fall below per-cell amplitudes because preferred phases are scattered across cells.

### Spatiotemporal maps

Frame maps give the spatial distribution of spiking in successive time windows after touch, in two projections of the same volume. They are built from the spike times of every neuron of the simulated volume, aligned to touch onset, and pool 5 seeds of 14 touches each.

#### Per-cell counts

Spike times were expressed relative to the nearest preceding touch onset and counted in 5 ms windows centred on 5, 10, 15, 20, 25 and 30 ms after onset. Counts were divided by the number of touches and then averaged across seeds, giving each cell one trial-averaged spike count per window.

#### Binning

Soma positions were mapped to the flatmap of the enhanced atlas (Section *Atlas and geometry*), giving each cell two flatmap coordinates and a cortical depth. Flatmap coordinates are dimensionless and were converted to micrometres by the nominal scale of 3570 *μ*m. Both projections use a 60 60 grid spanning the extent of the analysed cells rather than of the whole flatmap, 756 *μ*m by 624 *μ*m laterally and 1321 *μ*m in depth, giving bins of 12.6 *μ*m and 10.4 *μ*m along the two flatmap axes and 22 *μ*m in depth. The value of a bin is the sum of the trial-averaged counts of the cells falling in it. The flatmap view bins the two flatmap coordinates and collapses depth, so a bin corresponds to a curved, layer-spanning column of tissue. The coronal view bins one flatmap axis against depth and collapses the other, so a bin corresponds to a slab spanning the rows of the grid. Intensity is therefore a spike count per flattened voxel, not a rate or a density, and the two views are not comparable to each other in absolute value. The coronal panels are displayed to 1200 *μ*m, so the deepest bins fall outside the plotted range.

#### Scale and overlays

Within a figure the two projections share one linear colour scale, from zero to the largest bin value across both. Barrel outlines on the flatmap views are the flatmap intersections of the atlas barrel annotation, drawn for the nine barrels of the grid and, in lighter weight, for the surrounding barrels; on the coronal views the layer boundaries are the median depth of each layer interface computed in 30 lateral bins, and the vertical lines mark the flatmap extent of C2. The flatmap axes are plotted with the two coordinates exchanged relative to their order in the circuit files, so that the barrel grid appears in the orientation of the atlas overlay used throughout.

### Touch response metrics

Both analyses of the touch response use the touch simulations, 70 trials per cell over 5 seeds, with spike times aligned to touch onset after correcting for the jitter applied to the nominal touch times. Lamp5 cells were excluded throughout, too sparsely active to characterise. Responses were characterised per cell in C2 barrel column, the septum adjoining the grid, the row neighbours C1 and C3, and the arc neighbours B2 and D2.

#### Spiking

Rates were counted per trial in a 0-100 ms response window and a 100-0 ms baseline window. A cell silent on a trial contributes a zero. Per cell, we averaged the rate change over all 70 trials, the first-spike latency over the trials on which it fired, and the reliability, the fraction of trials with a spike in the response window. Latency counts the first spike within 0-50 ms of onset. Only cells that fired contribute, so latencies are means over responding cells.

#### Membrane potential

Action potentials were removed using the recorded spike times. The trace is held at its value 0.5 ms before the peak until it returns below that value. Blocks of 20 to 60 ms around each touch were baseline-subtracted per cell and trial over the 20 ms before onset. We averaged across trials, then across the cells of a class within a seed, then across the 5 seeds. The standard error is taken across seeds. Pooling across layers weights each class by its cell count. Within 0-50 ms of onset we took the peak deviation, its latency and the mean deviation, per class and seed. The peak is the largest absolute value, so hyperpolarising responses keep their sign.

#### Population onset and offset

Population rates were counted per layer and barrel group in 1 ms bins and baseline-subtracted over the 50 ms before onset. Onset and offset are the first and last times the trace crosses a fifth of its own peak, so the two are comparable between groups of differing response amplitude.

### Responder selection

Responders were selected over the neurons of the analysis volume and then examined by barrel. A cell is a responder when it passes all conditions of Table 4.

**Table 4.**
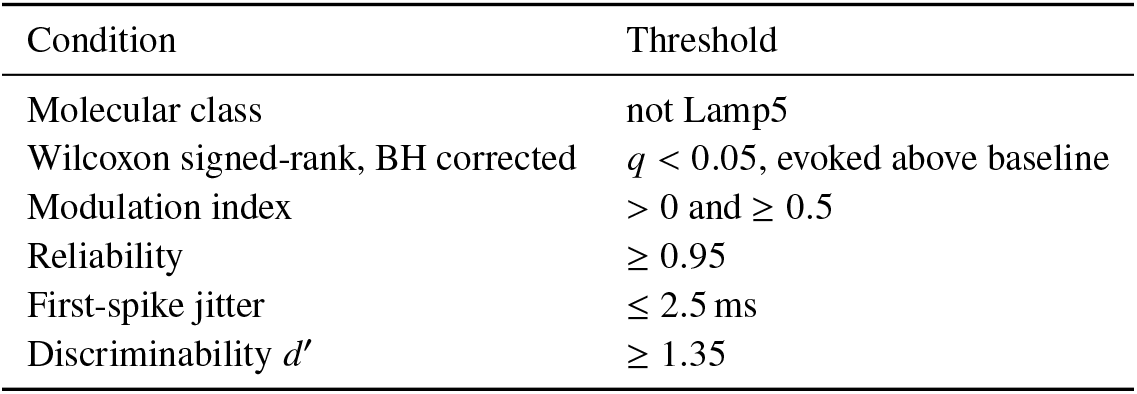
Conditions for selection as a touch responder. All six must hold.

#### Response window

The window is set per class, class being layer and molecular identity, since onset and duration differ between them. The population histogram of a class was pooled over the 5 seeds and computed from 100 to 100 ms in 2 ms bins, smoothed with a Gaussian of two bins. The threshold is the mean of the pre-touch bins plus two standard deviations. The window opens at the first bin after touch above the threshold. It closes at the first bin after the peak below it, capped at 100 ms. Classes whose histogram never crosses the threshold use a fixed window of 0-50 ms after touch onset. The baseline window ends at onset and matches the class’s response duration, capped at 100 ms. Spikes are counted in both windows, so the counts are comparable.

#### Criteria

Evoked and baseline counts were compared per cell across trials by one-sided Wilcoxon signed-rank test with Benjamini-Hochberg correction across cells. Cells silent in both windows, or with a constant difference across trials, were not tested. Reliability is the fraction of trials with at least one spike in the response window. The modulation index is *μ*_resp_ *μ*_base_ *μ*_resp_ *μ*_base_, running from 1 for a cell silenced by touch to 1 for one firing only after it. The threshold of 0.5 is a response three times the baseline. Jitter is the standard deviation across trials of the first-spike latency, over the trials on which the cell fired. Latency itself was not used, since the laminar spread of onset times is a property of the response rather than of response quality. Discriminability is

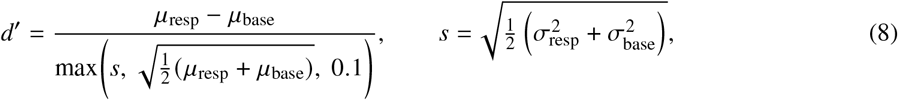

the pooled standard deviation floored at the Poisson expectation for the counts. Without a floor, a cell firing one spike on every trial and never at baseline has zero variance and unbounded *d*^′^.

#### Thresholds

Reliability was set below its ceiling of 1, which excludes a cell for failing a single trial in 70. Each threshold was then varied in turn with the others held fixed, reliability over 0.80, 0.95, *d*^′^ 1.30, 1.45, jitter 1.5, 4.0 ms and MI 0.30, 0.95 . The number of responders varies over these ranges. The composition by class and the fraction lying in C2 barrel column do not.

### Cyclicity analysis

The touch response was decomposed with cyclicity analysis (Nguyên et al., 2025), which recovers the recurrent temporal structure of a population response and assigns every cell a position within it. The method was applied here to the spiking activity of the simulated volume.

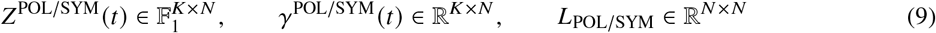

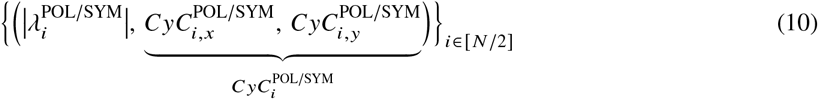

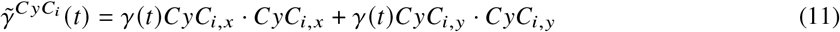

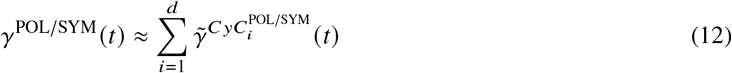

#### Filtering

Cyclicity analysis requires the signal to be differentiable to compare the rates of change between any two neurons. For that purpose, we filter the spike trains such that the spiking information is propagated forward in time and that the signal is continuous and differentiable almost everywhere.

The 2 ms time-binned spike raster *Z* (9), a binary signal of size *K* × *N* where *K* is the number of time bins and *N* the number of neurons, was reduced to a continuous signal per cell by convolving with a one-sided exponential kernel of *τ* = 5 ms, giving *γ_n_* (*t*) for cell *n* (9). The lead matrix *L*, indexed on neurons, was computed on the trials concatenated, written *γ*_concat,*n*_ (9). The projections and partial reconstructions that follow are taken on the trial-averaged signal *γ_i_* (*t*).

#### Lead matrix

Similarly to how PCA uses the eigen-decomposition of the covariance matrix to obtain an orthonormal basis, i.e. PCs, from which to decompose the cell space, Cyclicity Analysis relies on obtaining an orthonormal basis of planes from the decomposition of the lead matrix.

Pairwise temporal relations were collected in the lead matrix *L* (9), whose entry *L_i j_* is the lead of cell *i* over cell *j*, computed as the signed area enclosed by the joint trajectory of the two signals in the plane they span.

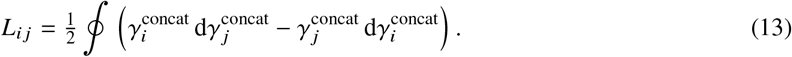

The entry is positive when cell *i* leads cell *j* and changes sign when the order reverses, so *L* is skew-symmetric by construction. A pair of cells whose activity rises and falls together encloses no area and contributes nothing, which is what distinguishes the measure from a covariance: it is sensitive to the order of activation and blind to shared amplitude.

#### Components

Because *L* is real and skew-symmetric, its eigenvalues *λ_i_* are purely imaginary and occur in conjugate pairs, each pair corresponding orthogonal eigenvectors span a two-dimensional invariant subspace of the cell space, which we call *CyC_i_* (and the underlying pair of eigenvectors is denoted as *CyC_i_*_,*x*_, *CyC_i_*_,*y*_respectively) (10). Much like for a symmetric matrix, such as the covariance matrix, all eigenvectors of a skew-symmetric matrix are orthogonal, resulting in a natural decomposition of the space by projecting onto the eigenvectors. Projecting *γ* onto such a plane gives a closed trajectory, and the area it encloses is given by the corresponding eigenvalue. The planes were therefore ordered by the magnitude of their eigenvalue, and the leading plane, the one maximising the enclosed area, is the first cyclic component CyC1.

#### Partial reconstruction

Projecting the activity back onto a plane and returning it to cell space gives *γ̃*^CyC^*^k^* (11), the part of the response carried by that component alone, which can be binned and mapped exactly as the spiking activity was. Each component approximates the response at a single frequency, so a partial reconstruction takes negative values over parts of the time course; these are corrected by the higher components and are part of the decomposed signal rather than a suppression of activity. Any reconstruction from a subset of components is therefore an approximation, and the sum over all components returns the filtered response (12).

#### Successive components

Another way to understand *CyC*_2_ is that once *CyC*_1_ has been identified as the subspace associated with the largest eigenvalues, then applying the same procedure recursively on *γ γ̃^C^*^*yC*1^ results in *CyC*_2_, so that each successive component captures the deviation from the planes already extracted.

#### Per-cell radius and phase

Each component is returned as a pair of eigenvectors with one entry per cell. The two entries place the cell at a point in the plane, whose modulus is the cell’s radius in that component and whose argument is its phase. The radius gives how strongly the cell contributes to the mode and the phase its timing within the cycle, and the two are reported together as a point in the complex plane. Cells were treated as participating in a component when their radius exceeded 0.003, a threshold applied consistently across all analyses of participation. It removes cells whose contribution is indistinguishable from zero and whose phase is correspondingly undefined. The value was set heuristically by thresholding the signal at a range of radii and asking how much of the enclosed area of *CyC*_1_ survived, and taking the largest threshold retaining at least 75% of it.

#### Component selection

Eigenvalues in cyclicity analysis are enclosed areas, so unlike principal components they do not measure explained variance. The decomposition was therefore assessed by reconstruction error. For each cell we took the ℓ_2_ error along time between the trial-averaged response and its reconstruction from the first *k* components, then averaged across cells. The error is expressed as a ratio to the across-time variance of the signal, which is the error made by a constant-rate approximation. The time-averaged rate was subtracted per cell beforehand, and the error computed from touch onset to the end of the trial (Supplementary Figure S8D). The ratio reaches 0.48 with one component, 0.60 with two and 0.65 with three, and rises by less than 0.01 per component beyond that. Three is therefore the upper bound worth retaining. The comparison between circuits uses the first three components, since the common and contrast decomposition is defined per component and the third contributes to the integrated maps.

#### Seeds

The decomposition was computed from a single simulation seed, whose fourteen touches enter the lead matrix concatenated. The eigenvectors define a fixed cell space that spatial averages are then read against. Spatial and temporal averages of participation pool 5 seeds of 14 touches, projected onto the eigenvectors of that one decomposition.

### Structural predictors of cyclic component participation

Participation in a cyclic component is a property of the response, while the connectivity of a cell is fixed before the simulation runs. We asked how much of the first is predicted by the second, fitting a model of participation from afferent connectivity alone, separately per cell class and separately for each component.

#### Response variable

The decomposition assigns each cell a radius and a phase within each component. Together they define a point in the complex plane, *z_i_* = *r_i_e*^i^*^φi^*. This complex value was used directly as the response, so amplitude and timing enter the fit jointly rather than as two separate regressions. It was standardised per class by subtracting its mean and dividing by its standard deviation. Cells were included when their radius exceeded 0.003 in the component being fitted, the same threshold used throughout the analysis. It removes cells whose participation is indistinguishable from zero and whose phase is correspondingly undefined.

#### Cell classes

Fits were run per class over the analysis volume. Excitatory cells were taken per layer and all inhibitory cells of a layer pooled, giving eight classes across layers 2/3 to 6. Classes with fewer than 30 cells passing the radius threshold were not fitted.

#### Predictors

Predictors were the structural synaptic drive received per cell, computed per synapse as in Section *Structural synaptic drive* and summed over the afferents of a given source. Three groups were used: thalamic drive from VPM, cortical excitatory drive and cortical inhibitory drive. Excitatory and inhibitory predictors therefore lie on different scales and are not comparable in magnitude to each other. Cells with no afferents of a given source take zero. All predictors were log 1 *x* transformed before fitting, since the relationship between drive and participation is closer to logarithmic than linear.

#### Fit

The model is a complex regression *z* = *Xw*, solved by least squares. The design matrix *X* is real and carries an intercept column, and the weights *w* C*^p^* are complex. Complex weights couple amplitude and phase through shared coefficients, which a joint fit to the real and imaginary parts as two independent responses does not. Goodness of fit was measured as

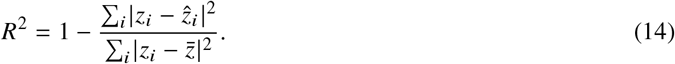

#### Variance partitioning

The unique *R*^2^ of a predictor group is the drop in *R*^2^ when that group is removed from the full model and the remaining groups refitted, floored at zero. Unique values are lower bounds on the contribution of each group, because the groups are collinear. A cell receiving more thalamic drive also tends to receive more cortical excitatory drive. The variance attributable to either alone is therefore smaller than the variance the two share. The unique values accordingly sum to less than the full-model *R*^2^. We report each group’s share of their sum alongside the absolute value. The comparison between components is less affected than the absolute values, since the same collinearity applies to both fits.

#### Cross-validation and significance

Predictive performance was estimated by 5-fold cross-validation with shuffling under a fixed seed. The model was refitted on each training split and *R*^2^ evaluated on the held-out fold. Cross-validated values fall below the full-model values throughout, and turn negative in the sparsest classes, where the number of cells passing the radius threshold is too small for the fit to generalise. Those rows are reported as descriptive only. Significance of the full model was assessed by permuting the response across cells 500 times and refitting. The p-value is the fraction of permutations reaching or exceeding the observed *R*^2^.

### Population coupling

Population coupling was computed following Bimbard et al. (2026), on the C2 barrel column during whisking. Analysis pooled 25 seeds of the whisking-only set, discarding the first 1 s of each.

#### Spike-triggered population rate

Spike trains were binned at 1 ms. For each cell, the profile is the population deviation averaged around that cell’s spikes,

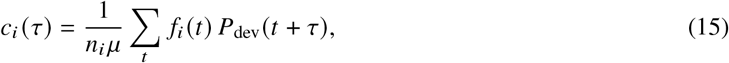

with *f_i_* the spike count of cell *i*, *n_i_* its total spikes, *μ* the mean population rate and *P*_dev_ the population deviation defined below. The cell’s own contribution is subtracted at every lag, so no cell is coupled to itself. Positive *τ* places the population after the cell’s spike. The population is the C2 barrel column alone and not the whole simulated volume. Profiles were computed over 80 ms and low-pass filtered at 20 Hz with a fourth-order Butterworth filter applied forwards and backwards.

#### Population deviation

The imposed whisking rhythm drives the whole population at 10 Hz. Subtracting only the mean rate would leave that deterministic modulation in every profile. The population rate was therefore binned by phase within the 100 ms cycle and the per-phase mean subtracted, leaving the fluctuation about the cycle-averaged rate.

#### Metrics

Coupling strength is the profile at zero lag. Coupling delay is the negated centre of mass of the profile over 35 ms. Negative values are clipped to zero beforehand so the weighting is non-negative, and the sign is flipped so that leaders are negative and followers positive. Coupling amplitude is the profile evaluated at the cell’s own delay, interpolated between neighbouring bins. It exceeds the strength whenever the profile peaks off zero. The two together separate how strongly a cell couples from how synchronously.

#### Seed combination and inclusion

Profiles were computed per seed over that seed’s own duration. They were combined as a mean weighted by the spikes each seed contributed for that cell. Cells were included when their delay was finite and they contributed at least 20 spikes to each of two disjoint halves of the seed pool, leaving 4,106 cells. Choristers are the top 15% by strength (616 cells) and leaders the earliest 10% by delay (411). All remaining included cells are soloists.

#### Reliability

The seed pool was split into two disjoint groups, alternate seeds to each. Both metrics were recomputed independently within each group and compared by Spearman correlation across cells (Supplementary Figure S9B). Delay reliability was assessed on the cells passing the 20-spike inclusion rule, the same filter used throughout. The delay-based classification is used at the population level rather than to assign individual cells.

#### Departure from the original

The centre of mass was taken over 35 ms rather than 80 ms, since the wider window left the estimate too noisy at the spike counts available. The narrower window coincides with the 35 ms sequence width reported across brain regions by Bimbard et al. (2026).

### Common and contrast decomposition

To separate the part of the touch response shared by the two circuits from the part unique to each, the cyclicity decomposition was applied to both and their components compared. The two projections overlap substantially. That is the observation the decomposition starts from: most of the response is common to the circuits, and only a part of it distinguishes them.

#### Setup

The decomposition was computed independently for each circuit. *γ*^POL^ and *γ*^SYM^ are the signals recorded in the two configurations. Each decomposition gives, per cyclic component *k*, a pair of eigenvectors, and the signal of that circuit is projected onto its own plane. The common and contrast planes are computed from the obtained cyclicity decomposition.

#### Common and contrast planes

Within each component, a common plane was defined as the plane maximising the overlap between the two projections (17). The dimensions remaining after the common plane is extracted were assigned to a contrast space. In all cases we inspected, this resulted in a common plane and a two-dimensional contrast space, accounting for the four dimensions spanned by two highly overlapping but linearly independent planes (18). Each signal then splits into a component lying in the common plane, shared with the other circuit, and a component in the contrast space, unique to it,

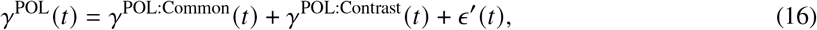

and the same for *γ*^SYM^, with *ε*^′^ the residual not captured by the component. The common plane is shared between the circuits by construction, so the two common trajectories are directly comparable within it. The contrast trajectories are not projections onto a shared plane. They are read as the activity present in one circuit and absent from the other.

#### Components

The decomposition was applied to the first three cyclic components. *CyC*_1_ is shown as the leading example (Figure 8D). The spatiotemporal maps of contrast and common activity integrate over all three (Figure 8E, Supplementary Figure S12). Reconstructions from the common and contrast parts (21), (22) follow the partial reconstruction of Section *Cyclicity analysis* and take negative values for the same reason.

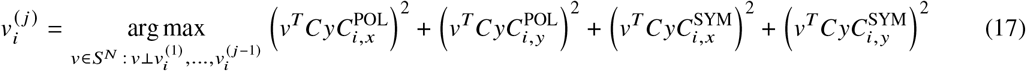

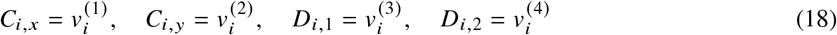

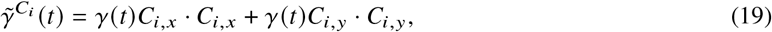

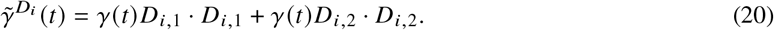

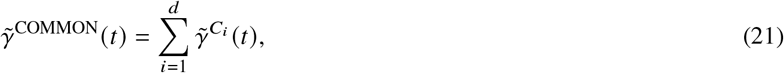

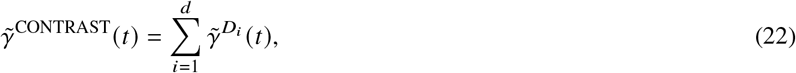

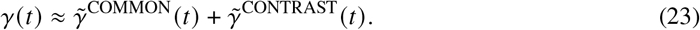

### Contrast orientation and lead-lag per cell

#### Cell position in a component

The cyclicity decomposition returns each cyclic component *CyC_k_* as a pair of eigenvectors of the lead matrix, *CyC_k_*_,*x*_ and *CyC_k_*_,*y*_, each with one entry per cell. For component *k* and cell *i*, the pair of entries *CyC_k_*_,*x*,*i*_, *CyC_k_*_,*y*,*i*_places that cell at a point in the plane spanned by the two eigenvectors. We call this point the cell’s loading, ℓ*_i_* R^2^. The population trajectory in the plane is the projection of the filtered activity onto the two eigenvectors,

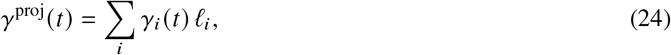

so activity in cell *i* displaces the trajectory along ℓ*_i_*, and the direction of ℓ*_i_* gives the direction in the associated plane.

#### Orientation

Orientation asks which cells drive the early separation of the two loops. It was computed in the contrast plane of the first component, where the projections of *γ*^POL^ and *γ*^SYM^ both start at the origin and run in opposite directions. Both trajectories were truncated at 20 ms post-touch onset and the endpoints *p*^POL^ and *p*^SYM^ of the truncated loops were taken. The truncation falls just past the contrast peaks of both circuits, 12 ms in POL and 18 ms in SYM, so the loops have separated but have not returned towards the origin. The line through the two endpoints divides the plane into two half planes, one containing the POL loop and one the SYM loop. Writing *d* = *p*^SYM^ - *p*^POL^ for the segment between them and *d*^⊥^ for the unit vector orthogonal to it (in the contrast plane), oriented towards the SYM half plane, the orientation of cell *i* is

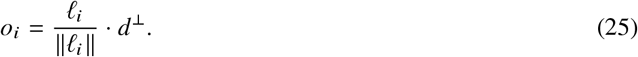

It takes values in 1, 1, negative for cells driving the POL loop and positive for cells driving the SYM loop. Its sign is the half plane the cell’s loading falls in, so the continuous value grades an assignment that is geometrically binary. Normalising removes the magnitude of a cell’s contribution and retains only its direction, so orientation reports which loop a cell builds and not how strongly it participates. The magnitude is reported separately as the radius of the component. All values reported here come from the first component.

### Lead-lag

Lead-lag is the lead of a cell’s POL response over its SYM response. For cell *i*, the two filtered responses are plotted against each other and the signed area enclosed by the resulting trajectory is taken. This is the functional of Equation 13, applied to one cell across the two circuits rather than to two cells within one. Values are averaged across trials. Lead-lag is positive when POL precedes and negative when SYM precedes.

### Inclusion

Cells contributing little to the component take small orientation values by construction, so they would count as not oriented rather than as absent. Cells were included when their radius in the first component reached the 75th percentile. This is a floor rather than a threshold: the radius distribution is smooth and right-skewed, with no separation between participating and non-participating cells. Lamp5 cells were excluded. Repeating at the 25th, 50th and 90th percentile leaves the ratio of POL- to SYM-oriented cells unchanged.

### Population summaries

The per-cell values of Figure 8G are the orientations of the included cells. The per-population values of Figure 8H are means in orientation and in lead-lag, taken across the included cells of a class with at least 20 such cells. The mean is calculated over cells.

## Data and Code Availability

Model produced is available in a public available at AWS through Open Brain Institute. Our code implementing the analysis and simulations is available in a public repository under the Apache-2.0 license.

## Author Contributions

**Aleksandra Teska:** Conceptualisation, Methodology, Software, Validation, Formal analysis, Investigation, Data Curation, Writing (original draft, review and editing), and Visualisation. **Tâm J Nguyên:** Signal decomposition, cyclic analysis. **Alexis Arnaudon:** Morphology synthesis, boundary algorithm, me-types setup. **Lida Kanari:** Morphology synthesis and dendritic analysis supervision. **Elvis Boci:** Graphical design of schematic figures. **James Isbister:** Circuit calibration. **Hugo Dictus:** Synaptic recipe setup. **Sylvain Crochet:** Supervision. **Felix Schürmann:** Conceptualization, Supervision. **Carl Petersen:** Conceptualisation, Supervision.

## Acknowledgements

This study was supported by funding to the Blue Brain Project, a research centre of the École polytechnique fédérale de Lausanne (EPFL), from the Swiss government’s ETH Board of the Swiss Federal Institutes of Technology.

We thank Dr Darshan Mandge for providing the e-models and their characterisation. We are also grateful to Dr Robin Dard, Lana Smith, Myriam Hamon and Morgane Storey for their feedback on the analysis and visualisations.

## Supplementary

**Supplementary Figure S1.**
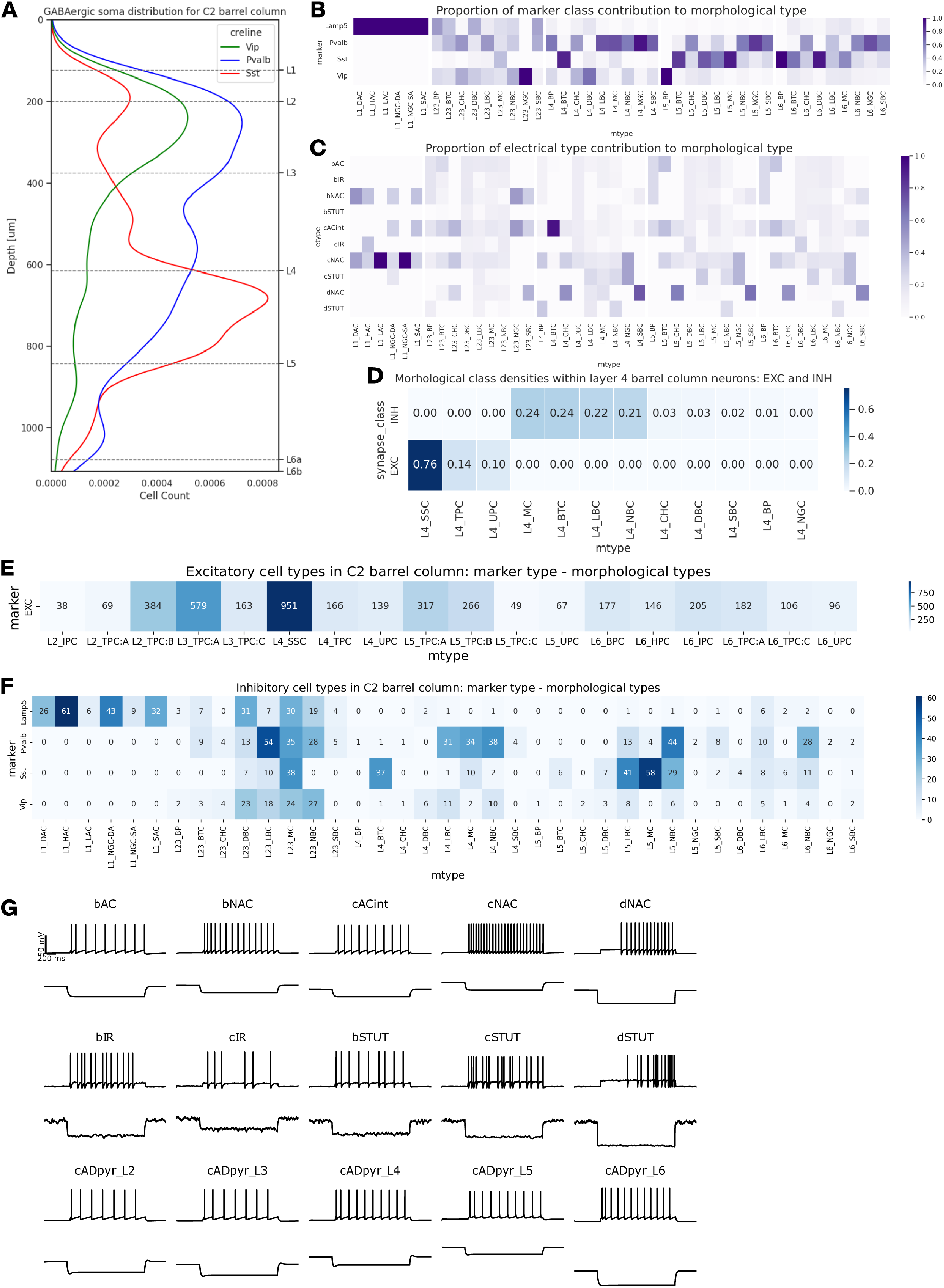
Circuit characterisation. **A.** Depth profile of inhibitory soma density across the C2 barrel column after model preparation, for Pvalb, Sst and Vip (Keller et al., 2018; Kim et al., 2017; Tremblay et al., 2016). **B.** Probabilistic mapping from morphological type to molecular marker (Roussel et al., 2023), used to assign markers in the model. Each column gives the distribution over markers for one m-type and sums to 1; each inhibitory neuron drew its marker stochastically from the column matching its type. Panels E and F show the resulting composition. **C.** Proportion of each electrical type per morphological type. Columns sum to 1. **D.** Morphological class composition of layer 4 neurons in the barrel column, excitatory and inhibitory separately. Excitatory composition adapted from Scala et al. (2019): spiny stellate cells make up 76% of layer 4 excitatory neurons. **E.** Excitatory neuron counts per morphological type in the C2 barrel column. **F.** Inhibitory neuron counts per morphological type in the C2 barrel column, split by assigned molecular marker. **G.** Firing behaviour of every electrical type in the model. *Top*: membrane potential in response to the somatic current injection shown below. *Bottom*: injected current, following the step protocol used to fit that e-type.

**Supplementary Figure S2.**
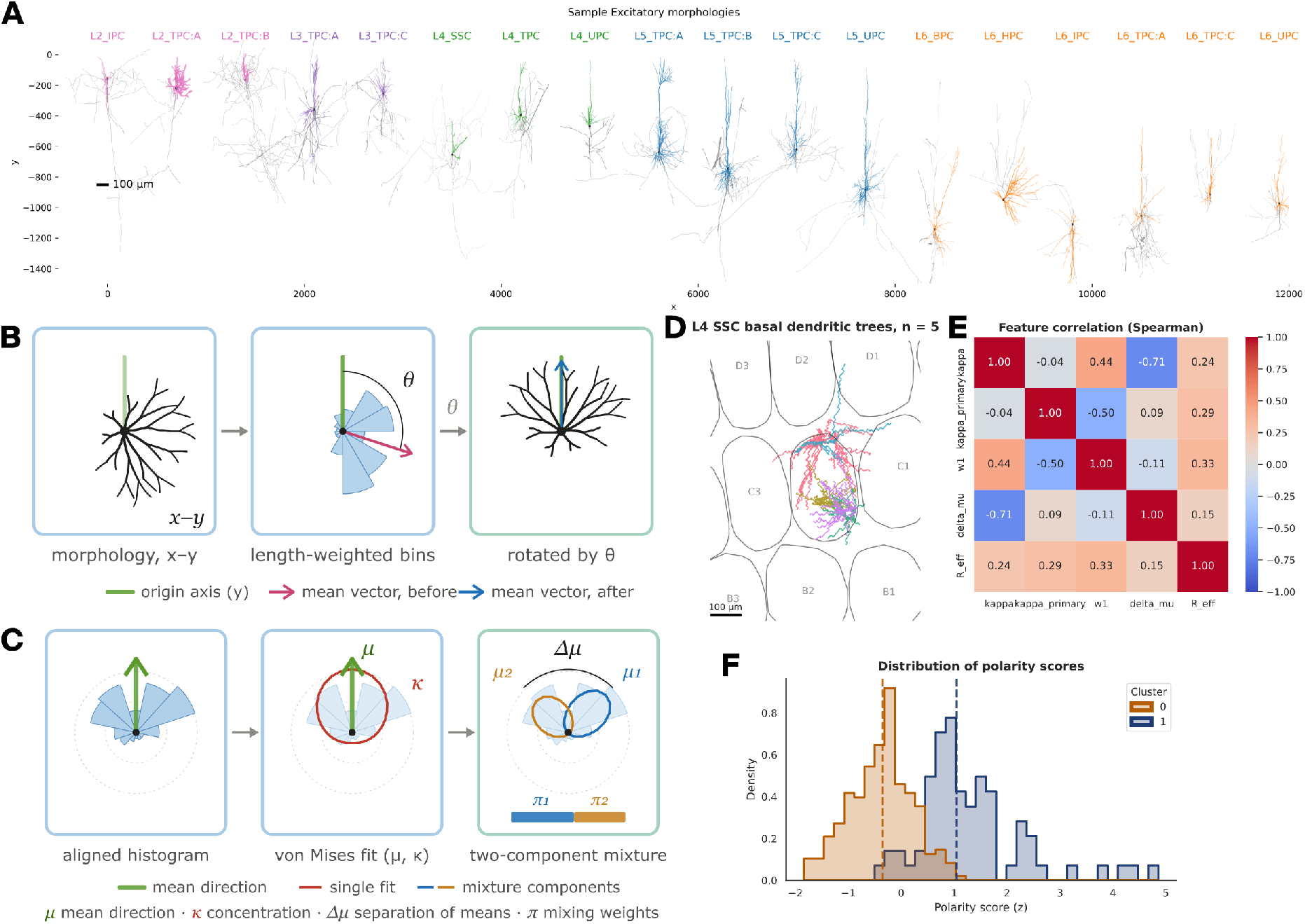
**Morphology sample and dendritic polarity quantification**. Related to Figure 2. **A.** Sample mousified excitatory morphologies, one per morphological type, positioned at their cortical depth. Dendrites coloured by layer, axons in grey. **B.** Alignment of dendritic orientation. Left: basal dendritic tree viewed in the flatmap plane. Middle: length-weighted angular distribution of dendritic mass around the soma, with the resulting mean vector (magenta) at angle *θ* from the origin axis (green). Right: the tree rotated by *θ*, bringing its mean direction onto a common axis so that distributions can be compared across cells. **C.** Descriptors extracted from each aligned histogram. Left: aligned distribution with its mean direction *μ*. Middle: single von Mises fit, giving the overall concentration *κ*. Right: two-component mixture, giving the concentration and weight of the dominant component and the angular separation Δ*μ* between component means. **D.** Basal dendritic trees of five L4 SSC reconstructions drawn at random, shown in the barrel grid. **E.** Spearman correlation between the five polarity descriptors across all reconstructions. **F.** Distribution of the composite polarity score for each cluster, with dashed lines marking cluster means.

**Supplementary Figure S3.**
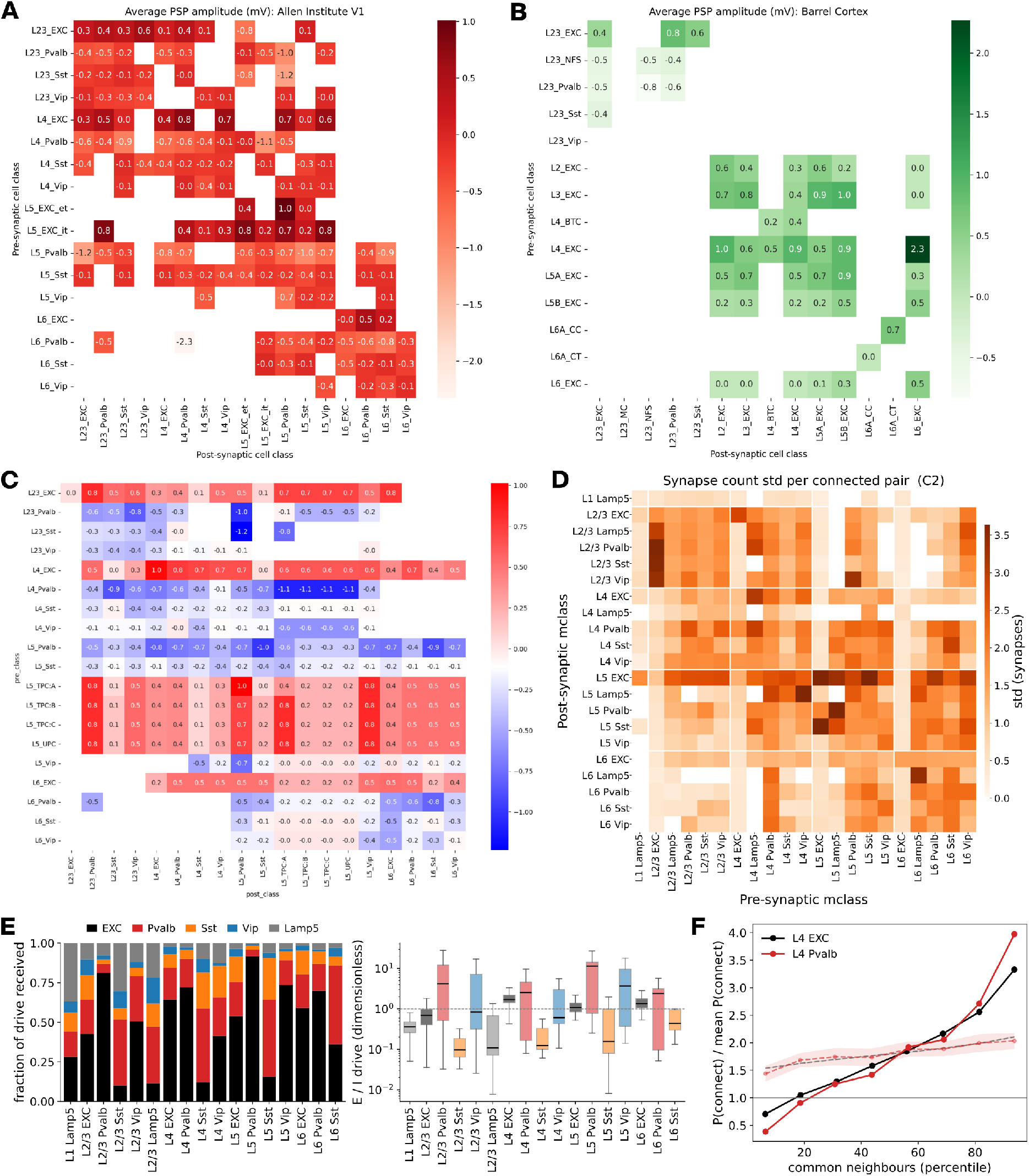
**Synaptic physiology, connectome characterisation, and synaptic input composition**. Related to Figure 3. **A.** Mean postsynaptic potential amplitude per molecular pathway from the Allen Institute V1 dataset, used as reference data for the model. **B.** Mean postsynaptic potential amplitude per pathway collected from the barrel cortex literature. Source studies characterised cells at different levels of specificity - molecular in some, morphological in others - so pathway labels differ between A and B and are reproduced here as reported. **C.** Postsynaptic potential amplitude recipe used in the model, obtained by extrapolating the reference data in A and B across layers and cell classes to cover all pathways. **D.** Standard deviation of synapse count per connected pair, per molecular pathway, within the C2 barrel column. Corresponding means in Figure 3D. **E.** Structural synaptic drive received by each molecular class in the C2 barrel column, from C2-internal inputs. Left: fraction of total drive received, split by presynaptic class. Right: ratio of excitatory to inhibitory drive per cell; boxes give median and interquartile range, whiskers 1.5 IQR, dashed line unity. Ratios are comparable across classes but not absolute. Classes with fewer than 15 cells omitted. **F.** Common-neighbour bias in layer 4 of the C2 barrel column, for L4 EXC (black) and L4 Pvalb (red). Points give the probability that two neurons within 150 *μ*m are connected, normalised to the population mean, against the percentile of their number of common neighbours, L4 neurons connected to both members of the pair. Dashed lines and shading give mean and SD across 50 distance-preserving shuffles.

**Supplementary Figure S4.**
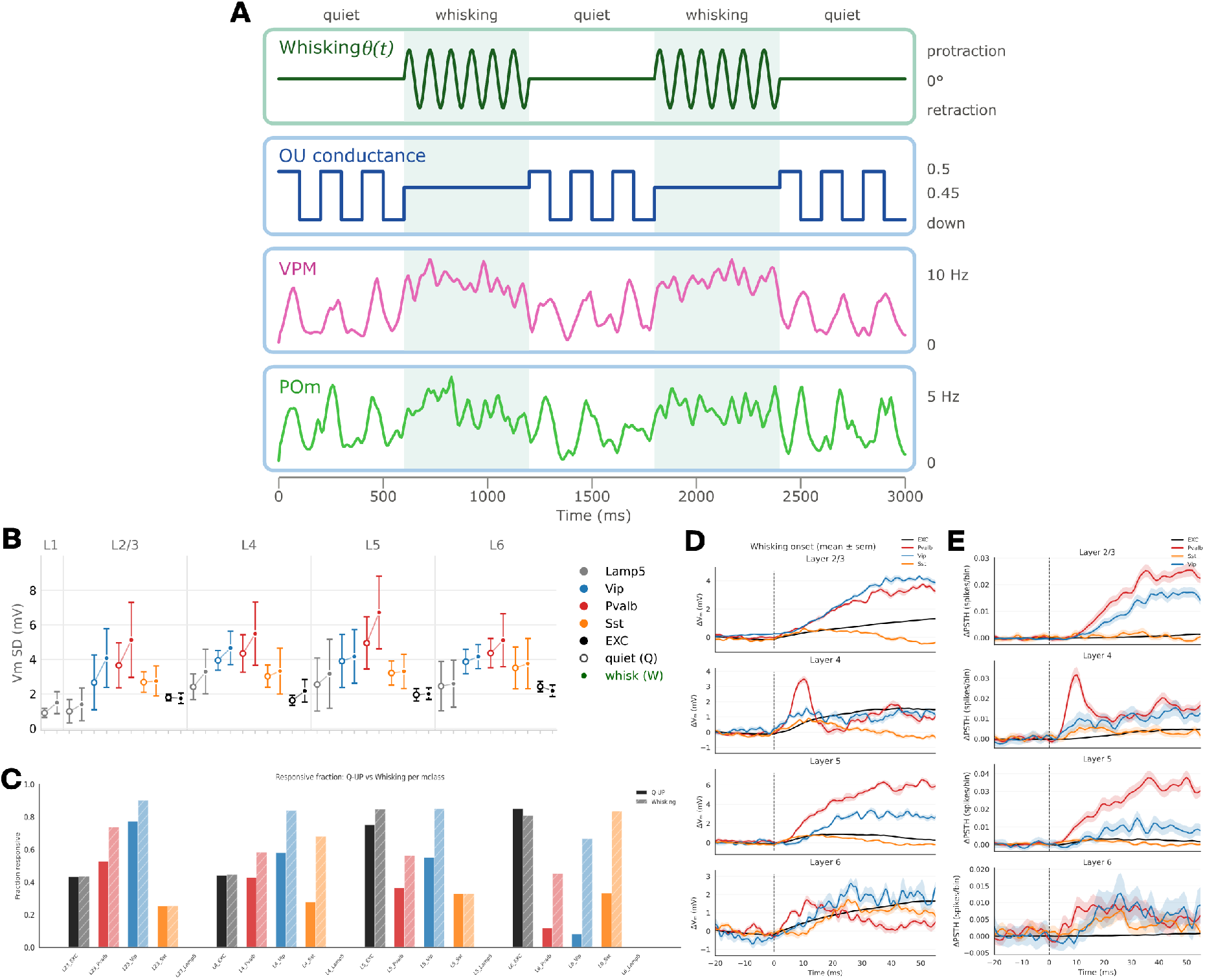
**Simulation inputs and state-dependent activity**. Related to Figure 5. **A.** Inputs driving the simulations. Whisker angle *θ t* sets the whisking periods, shaded throughout. Ornstein-Uhlenbeck noise supplies the cortico-cortical background, given as the relative ratio needed to reach the target spontaneous firing rate without VPM input, alternating between up and down levels during quiet and held at an intermediate level during whisking. VPM and POm rows give the measured population rates from the simulation. **B.** Standard deviation of the membrane potential in the C2 barrel column, per layer and molecular class, during quiet (Q, open) and whisking (W, filled). Mean SD across cells. **C.** Fraction of cells spiking at least once during the quiet up state and during whisking, per layer and molecular class. **D.** Membrane potential change at whisking onset, per molecular class and layer. From the simulation set of Figure 5G. Mean SEM across cells. **E.** Change in firing rate at whisking onset, same cells and alignment as D.

**Supplementary Figure S5.**
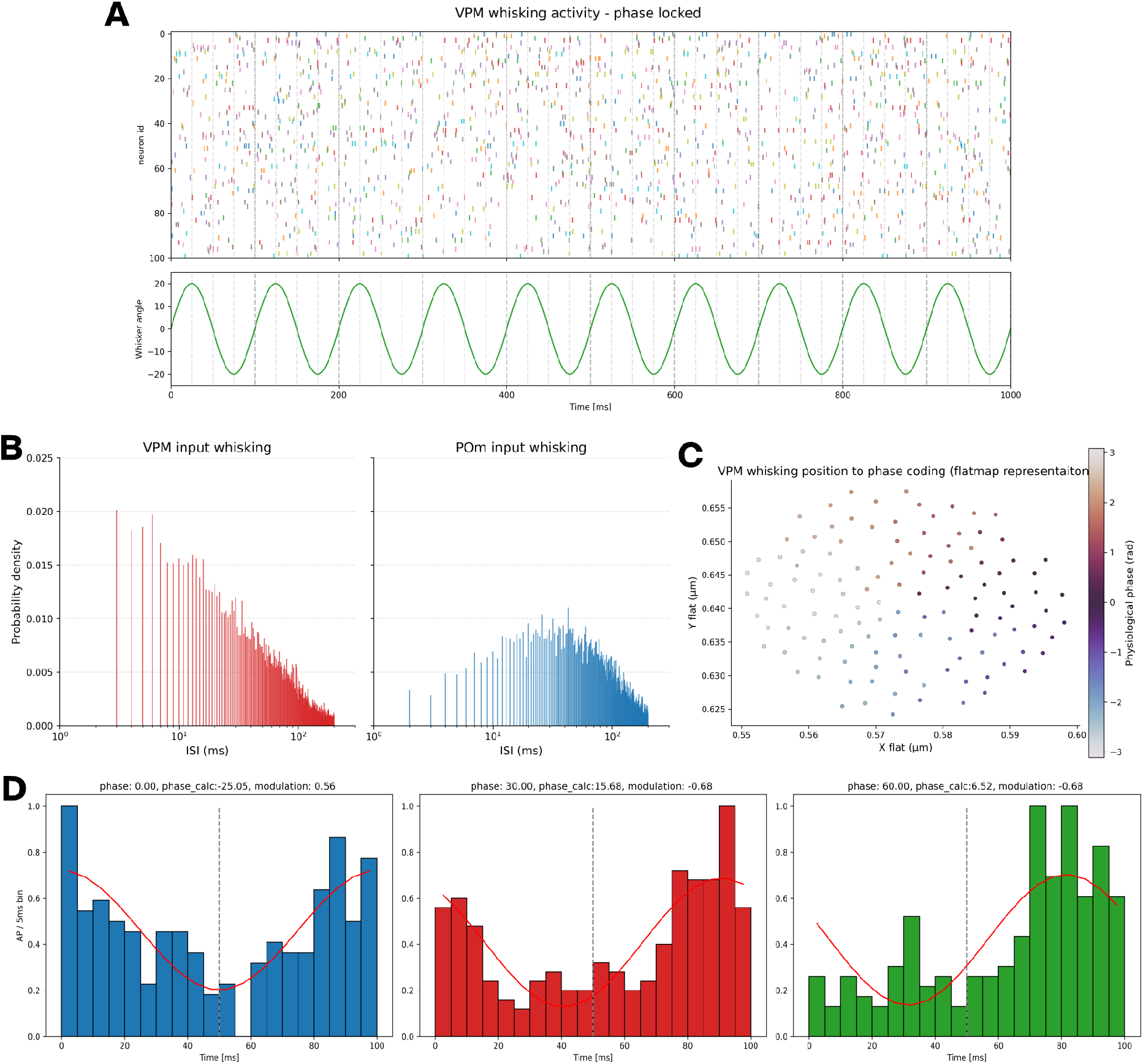
Whisking encoding in the thalamocortical input. Related to Figure 5. **A.** VPM spike raster sample during whisking, one colour per fibre, with the whisker angle below. Dashed lines mark 100 ms windows, within which the phase preference of successive fibres traces a diagonal. **B.** Interspike interval distributions of the whisking input. Left: VPM. Right: POm. **C.** Flatmap positions of the VPM fibres innervating C2, coloured by assigned whisking phase. **D.** Cycle-triggered histograms for three VPM fibres of differing phase preference, with the fitted sinusoid in red. Dashed line marks the midpoint of the cycle. Three cells sample.

**Supplementary Figure S6.**
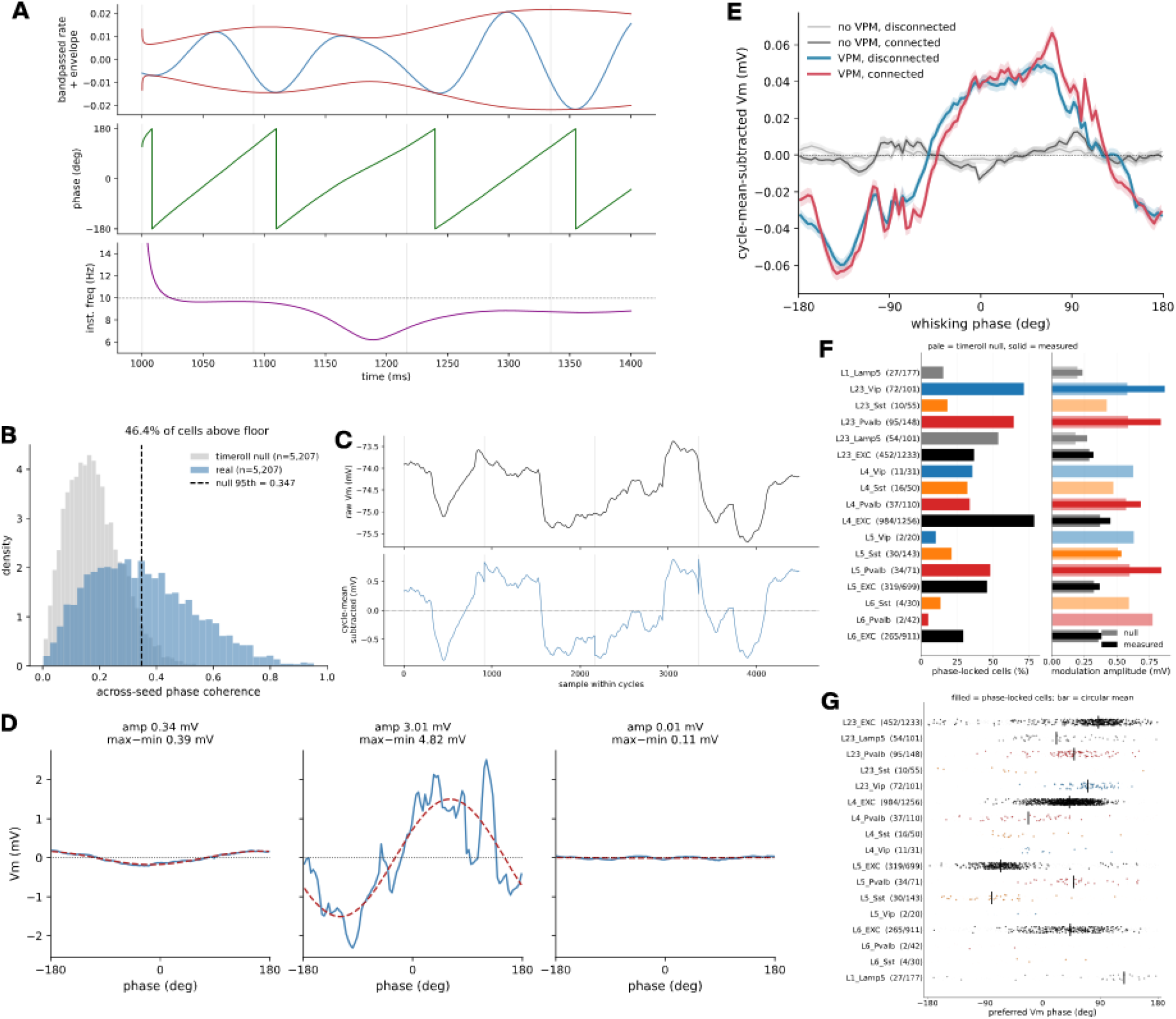
Membrane potential phase locking to the whisking rhythm. Related to Figure 5H. **A.** Phase clock derived from the C2 VPM population rate, for 500 ms window of one simulation. Top: band-passed rate with the envelope of its analytic signal. Middle: instantaneous phase. Bottom: instantaneous frequency. Grey lines mark cycle boundaries. **B.** Across-seed phase coherence for the real data and for a time-rolled null of the same cells. Dashed line gives the 95th percentile of the null, the phase-locking floor used throughout. **C.** Per-cycle mean subtraction, for one cell. Top: raw membrane potential across successive cycles. Bottom: the same trace after subtracting the mean of each cycle, removing drift between cycles while preserving within-cycle structure. Grey lines mark cycle boundaries. **D.** Three cells spanning the range of modulation, each showing the membrane potential averaged into 100 phase bins with its least-squares sinusoidal fit. Right: a cell below the phase-locking floor. The raw binned trace swings about 2.4 times wider than the fitted sinusoid, because it carries bin-to-bin noise that the fit does not follow. Amplitude is taken from the fit throughout. **E.** Population mean phase-binned membrane potential under four conditions, with VPM input present or absent and cortical connectivity intact or removed. Values are means across cells and fall below per-cell amplitudes because preferred phases are scattered. **F.** Phase locking per molecular class. Left: percentage of cells above the phase-locking floor. Right: modulation amplitude, pale bars the time-rolled null, solid bars the measured value across locked cells. Counts give locked cells over class size. Classes with fewer than 20 cells omitted. **G.** Preferred phase per molecular class, one point per cell. Cells above the phase-locking floor are drawn solid, the remainder faint. Bar gives the circular mean of the cells above the floor. L4 and L5 EXC concentrate on a preferred phase, while L2/3 and L6 EXC spread more widely. Inhibitory neurons are less numerous but still show a phase preference.

**Supplementary Figure S7.**
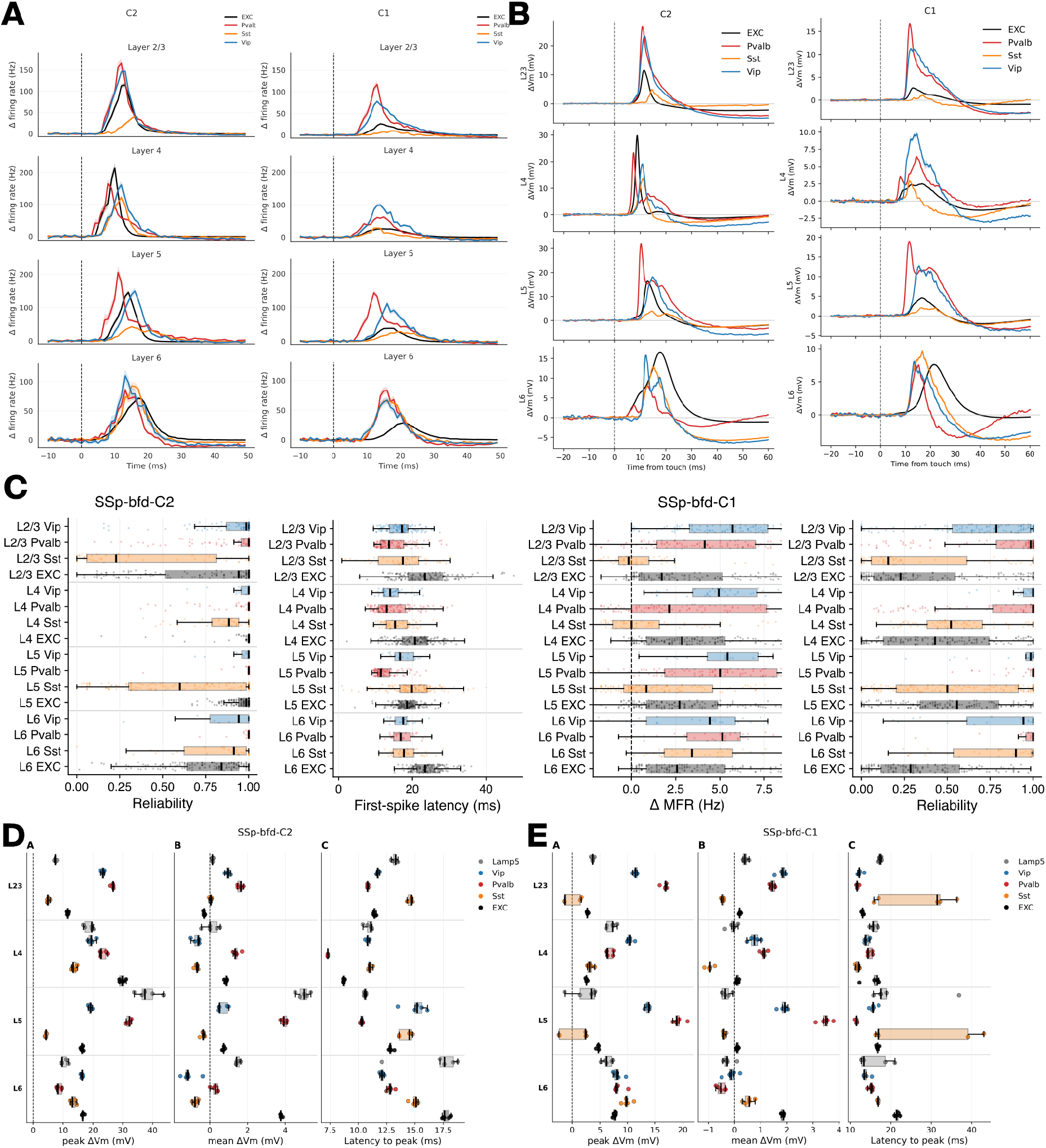
Extended characterisation of the cortical response to active touch. Related to Figure 6. **A.** Change in mean firing rate from touch onset, per molecular class and layer. Left: C2 barrel column. Right: C1. **B.** Change in membrane potential from touch onset, per molecular class and layer, same convention as A. **C.** Touch response characterisation per molecular class, extending Figure 6E. *Far left*: reliability in C2 barrel column, the other measures for C2 barrel column appearing in Figure 6E. *Remaining three*: first-spike latency, change in mean firing rate and reliability in C1 barrel column. Box plots with individual cells overlaid. **D.** Membrane potential response in C2 barrel column, per molecular class and layer. Left to right: peak change, mean change and latency to peak. **E.** Same as D, for C1 barrel column.

**Supplementary Figure S8.**
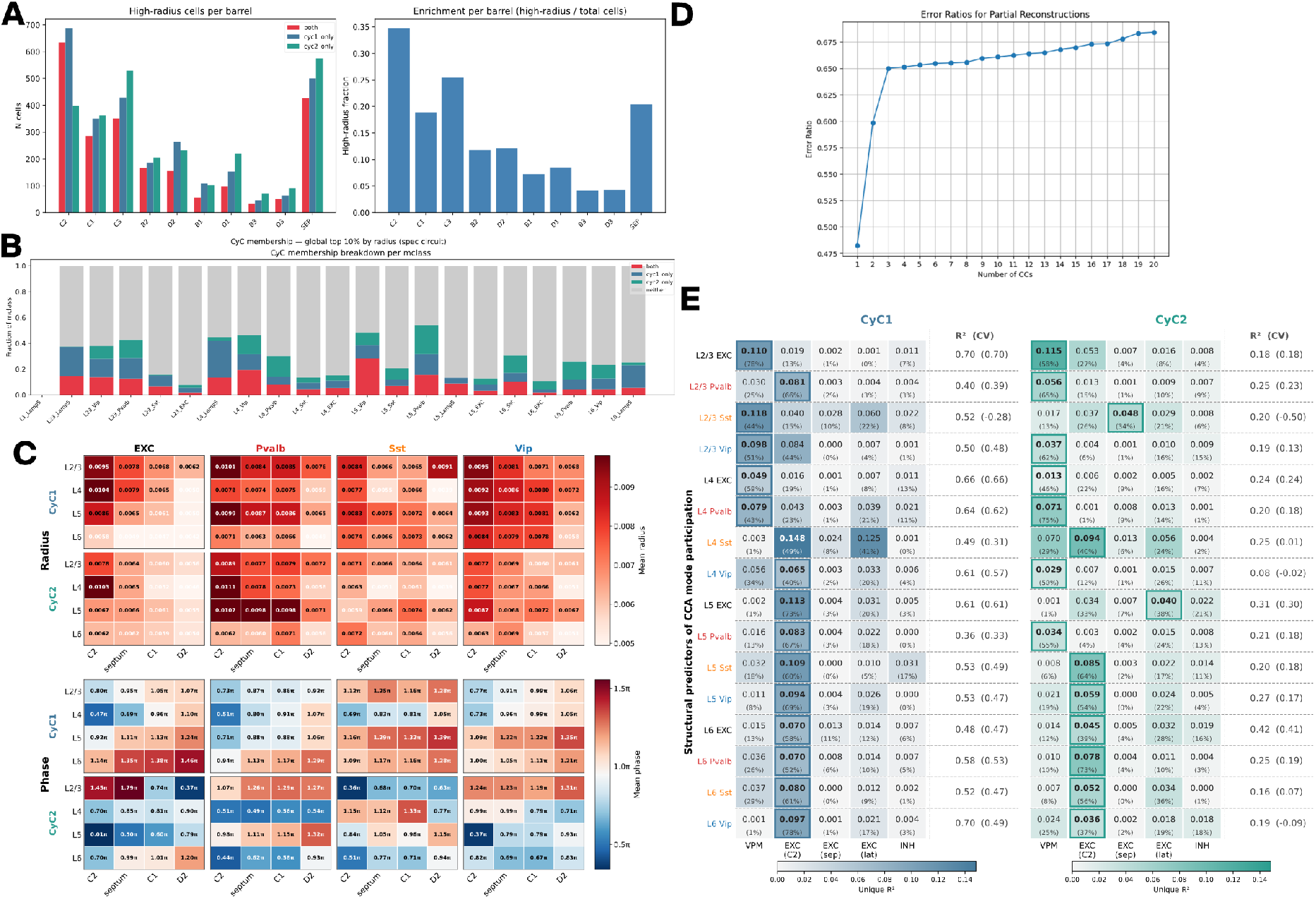
**Extended characterisation of CyC mode participation**. Related to Figure 6F-J. **A.** Cells in the top 10% by radius, per barrel and septum. Left: counts, split by whether a cell reaches the threshold for CyC1 only, CyC2 only, or both. Right: fraction of all cells in each barrel that reach it. **B.** The same membership, per molecular class. **C.** Mean radius (top) and mean phase (bottom) for CyC1 and CyC2, per layer, molecular class and barrel group. **D.** Reconstruction quality against number of cyclic components, POL circuit. Fraction of across-time variance of the trial-averaged response recovered by a partial reconstruction from the first *k* CyCs, averaged across neurons. Time-averaged rate subtracted per neuron before evaluation; error computed from touch onset to trial end. **E.** Structural predictors of CyC mode participation, extending Figure 6J to all layers and molecular classes and splitting the cortical excitatory predictor into within-C2, septal and lateral sources. Unique *R*^2^ per predictor group for CyC1 (blue) and CyC2 (green), with each group’s share of the summed unique *R*^2^ in parentheses. Right of each block: full-model *R*^2^, cross-validated *R*^2^ in parentheses. Boxed cells mark the dominant predictor. Cross-validated *R*^2^ falls well below the full-model value in the sparsest classes and turns negative in L2/3 Sst, L4 Vip and L6 Vip, where cell counts are too low for the fit to generalise; those rows should be read as descriptive only.

**Supplementary Figure S9.**
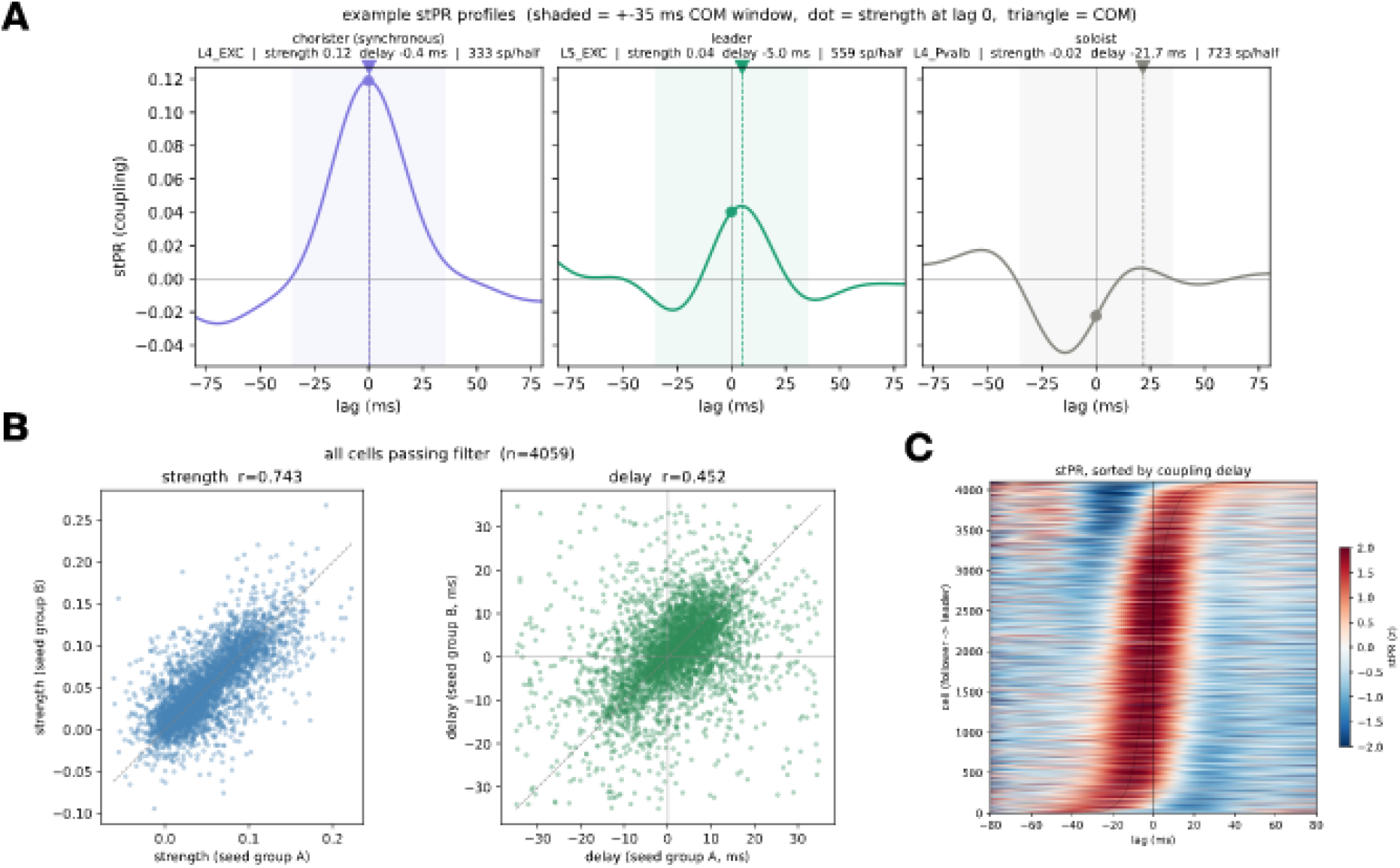
Population coupling classification. Related to Figure 7B. **A.** Spike-triggered population rate for three example cells, one per coupling class. Shading marks the 35 ms window used to compute the centre of mass; dot gives the value at zero lag, triangle the centre of mass. Class, coupling strength, delay and spike count inset. **B.** Split-half reliability of the coupling measures, for all cells passing the spike-count filter. Cells were divided into two seed groups and each measure computed separately. Left: coupling strength. Right: coupling delay. Dashed line is unity. **C.** Spike-triggered population rate for all cells, sorted by coupling delay.

**Supplementary Figure S10.**
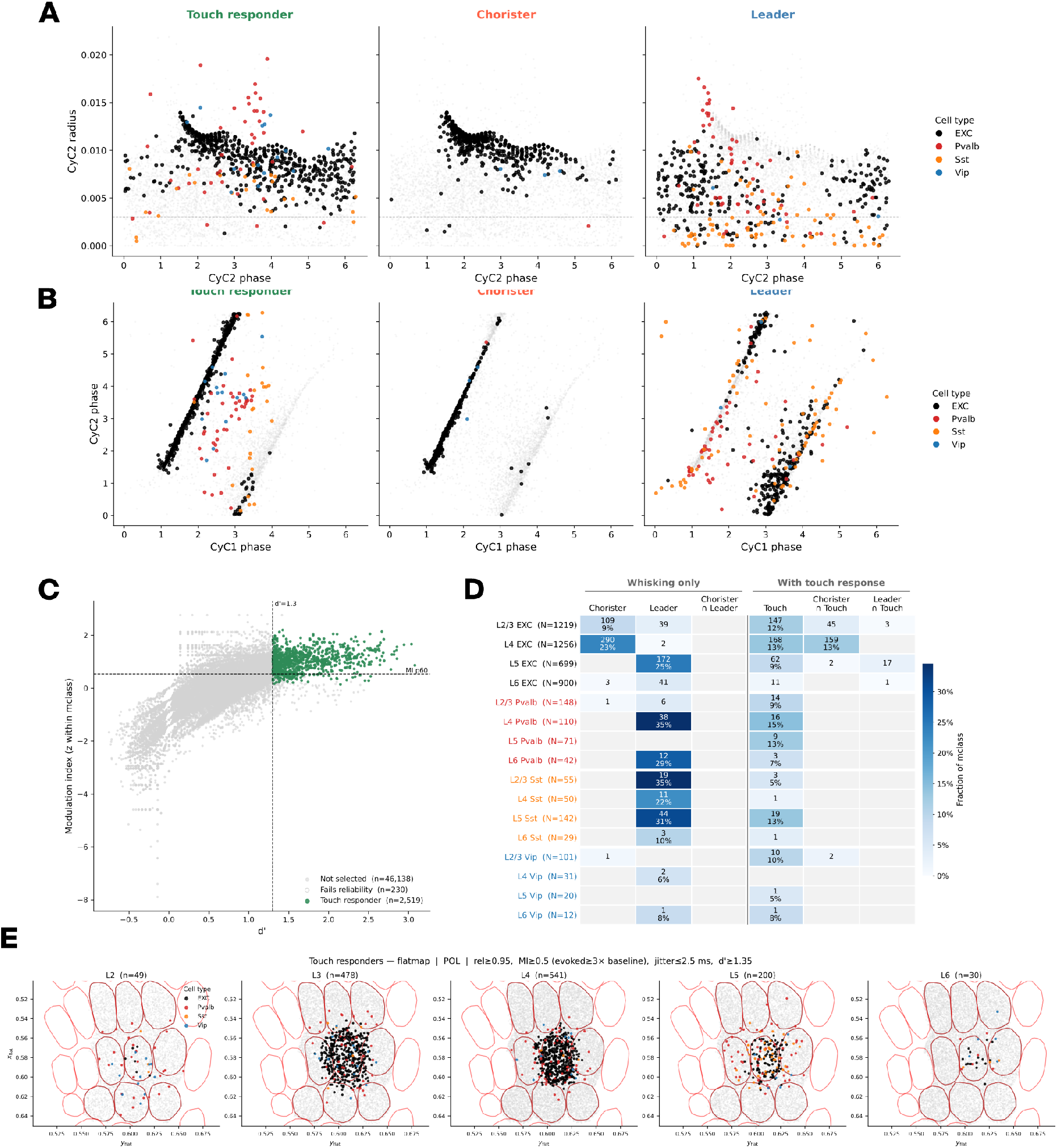
Extended characterisation of the functional cell populations. Related to Figure 7 **A.** CyC2 participation of each population, extending Figure 7C. Radius against phase for touch responders (left), choristers (middle) and leaders (right), coloured by molecular class; grey points are all cells. Dashed line marks the radius threshold of 0.003. Choristers occupy a narrow phase band, while leaders spread across the full cycle and include a Pvalb group at high radius and early phase. **B.** CyC1 against CyC2 phase for the same three populations. Cells fall along two bands: excitatory cells on a tight diagonal, where the two modes advance together, and inhibitory cells scattered off it. Choristers occupy the diagonal alone; leaders populate both. **C.** Touch responder selection, extending Figure 7A. Modulation index against *d*^′^ for all cells, with two gates drawn. Open points fail the reliability gate despite passing both. Counts cover all barrels, against C2 alone in Figure 7A. **D.** Overlap between the three populations, per molecular class. Left columns: cells belonging to one population only. Right columns: cells shared between each pair. Colour gives the fraction of the class, counts inset. L4 EXC is dominated by choristers and L4 Pvalb, L2/3 Sst and L5 Sst by leaders. Choristers and leaders never co-occur. **E.** Flatmap distribution of touch responders by layer, coloured by molecular class. Grey points are all cells; barrel outlines drawn. Responders concentrate in C2 in L3 to L5 and scatter across the field in L2 and L6. Selection thresholds inset.

**Supplementary Figure S11.**
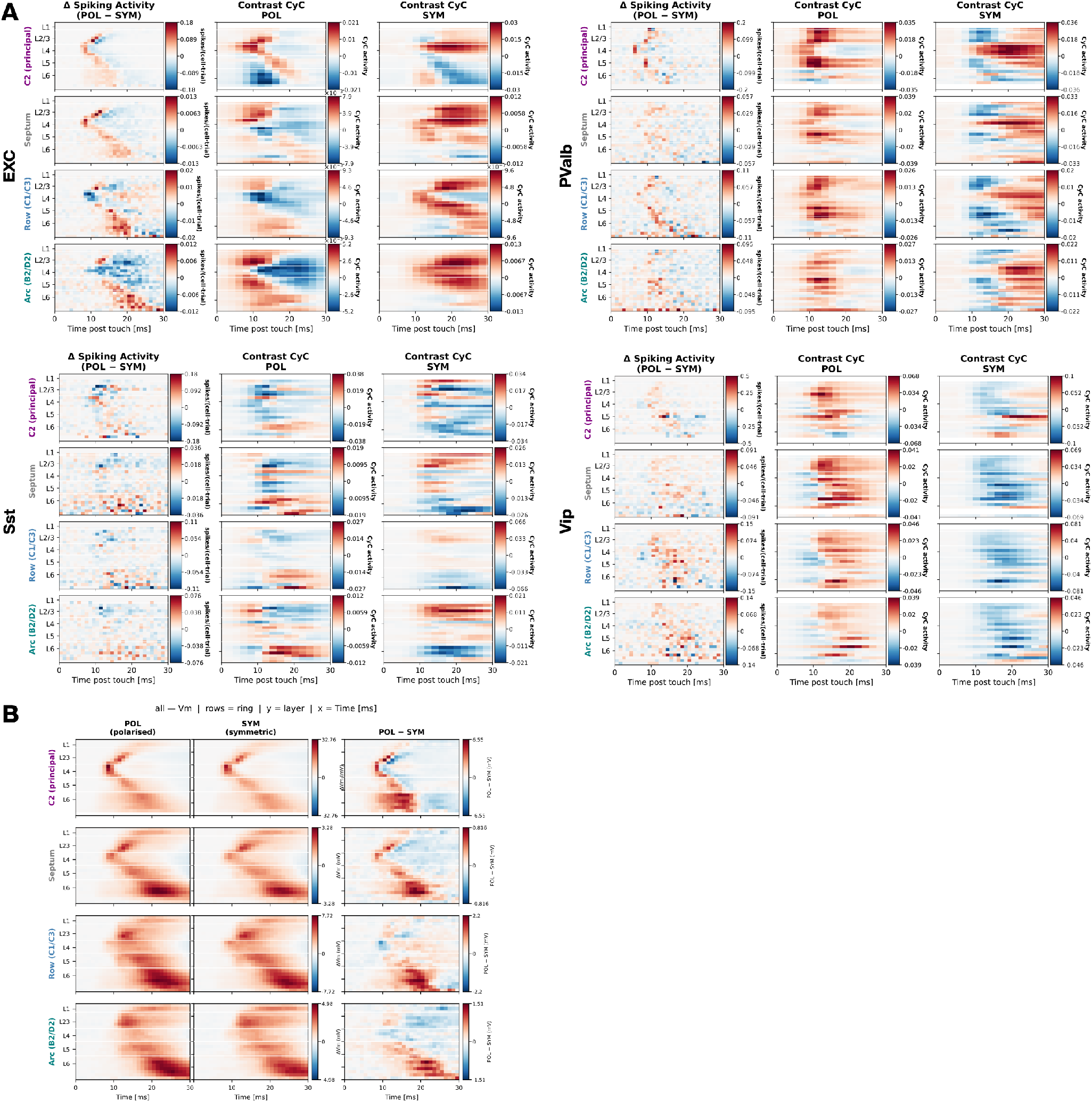
Circuit comparison per molecular class. Related to Figure 8F. Cortical depth on the vertical axis, time from touch on the horizontal, pooled over the area named at left - C2, septum, row neighbours (C1/C3) and arc neighbours (B2/D2). Within each block, left: spiking activity in POL minus SYM; middle and right: the component of activity unique to each circuit, in POL and in SYM. **A.** EXC. **B.** Pvalb. **C.** Sst. **D.** Vip. **E.** Membrane potential across all classes, same axes and areas. Left: POL. Middle: SYM. Right: POL minus SYM.

**Supplementary Figure S12.**
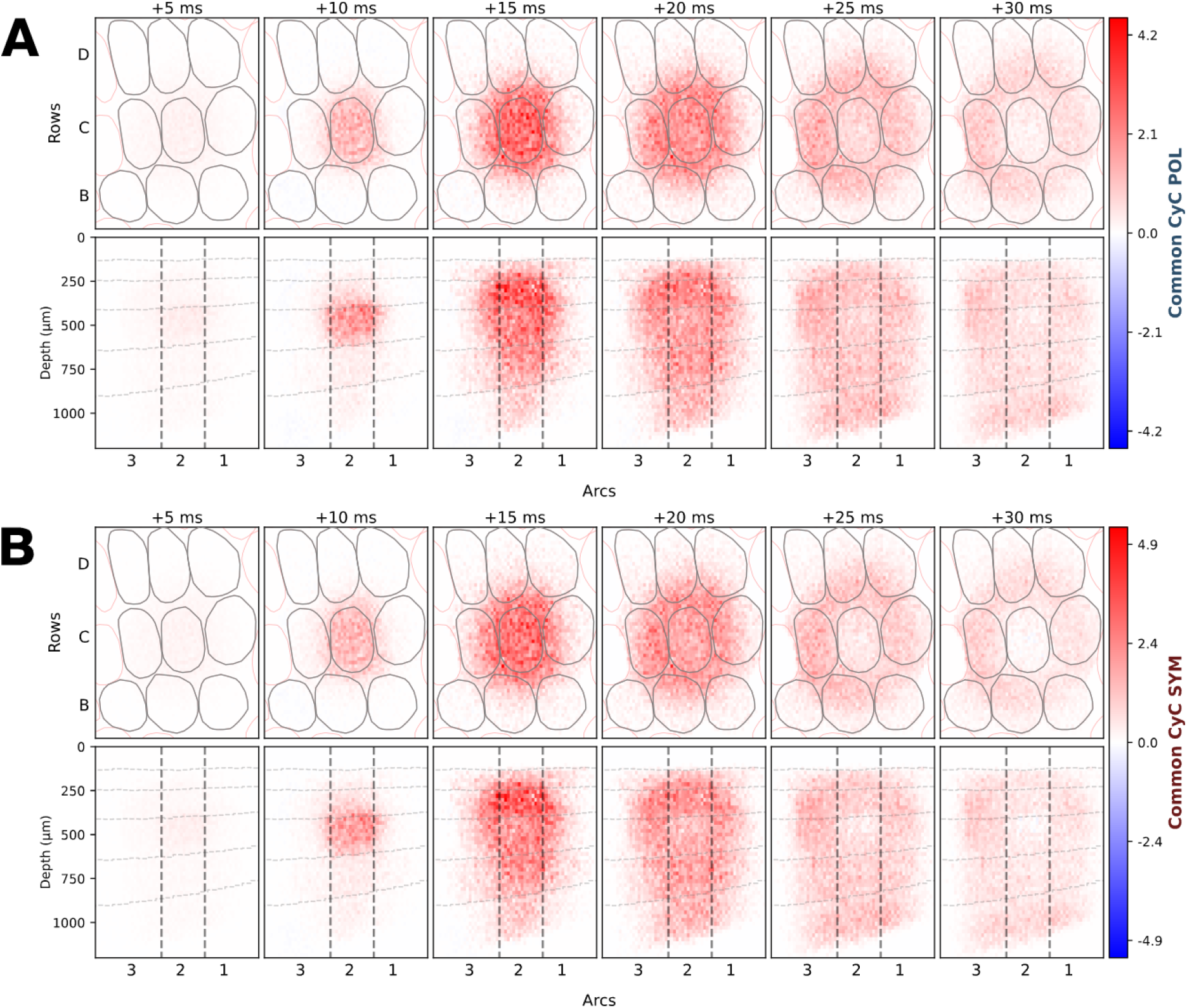
Common activity in the two circuits. Related to Figure 8D. Spatiotemporal structure of the common activity integrated over CyC1 to CyC3, plotted as in Figure 6C. **A.** POL. **B.** SYM. The two colour scales differ (4.2 against 4.9).

**Supplementary Figure S13.**
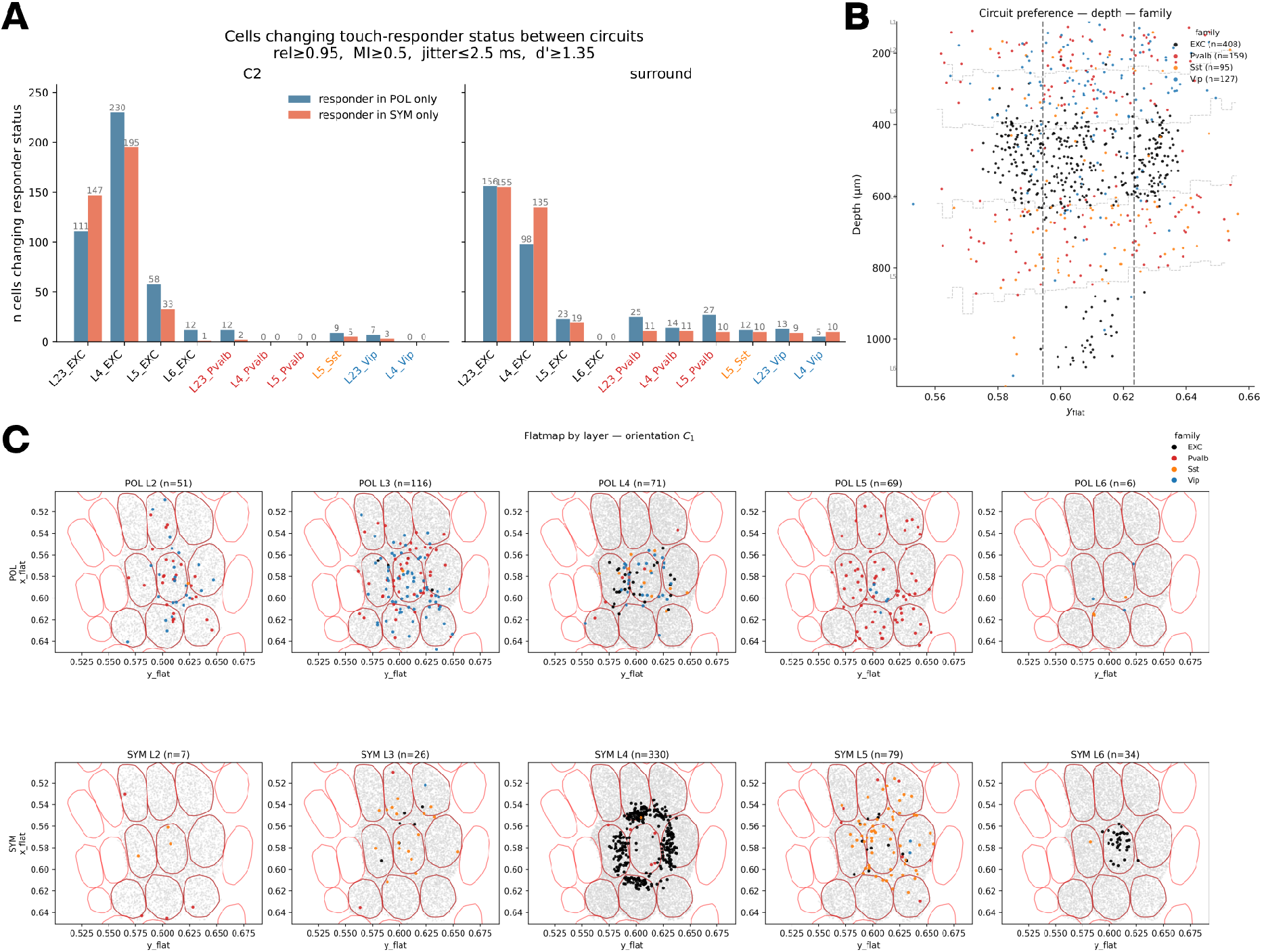
Circuit-exclusive responders and circuit preference. Related to Figure 8G and H. **A.** Cells changing touch-responder status between circuits, per molecular class, in C2 (left) and the surrounding barrels (right). Selection thresholds inset. **B.** Cells at the extremes of the circuit preference distribution, plotted by depth against flatmap position and coloured by molecular class. Dashed lines mark the C2 boundary. **C.** Flatmap distribution by layer of the cells preferring each circuit, coloured by molecular class. Top: POL. Bottom: SYM. Grey points are all cells; barrel outlines drawn. POL-preferring cells are largely Pvalb and Vip across L2/3 and L5, SYM-preferring cells largely L4 EXC.

**Table S1.**
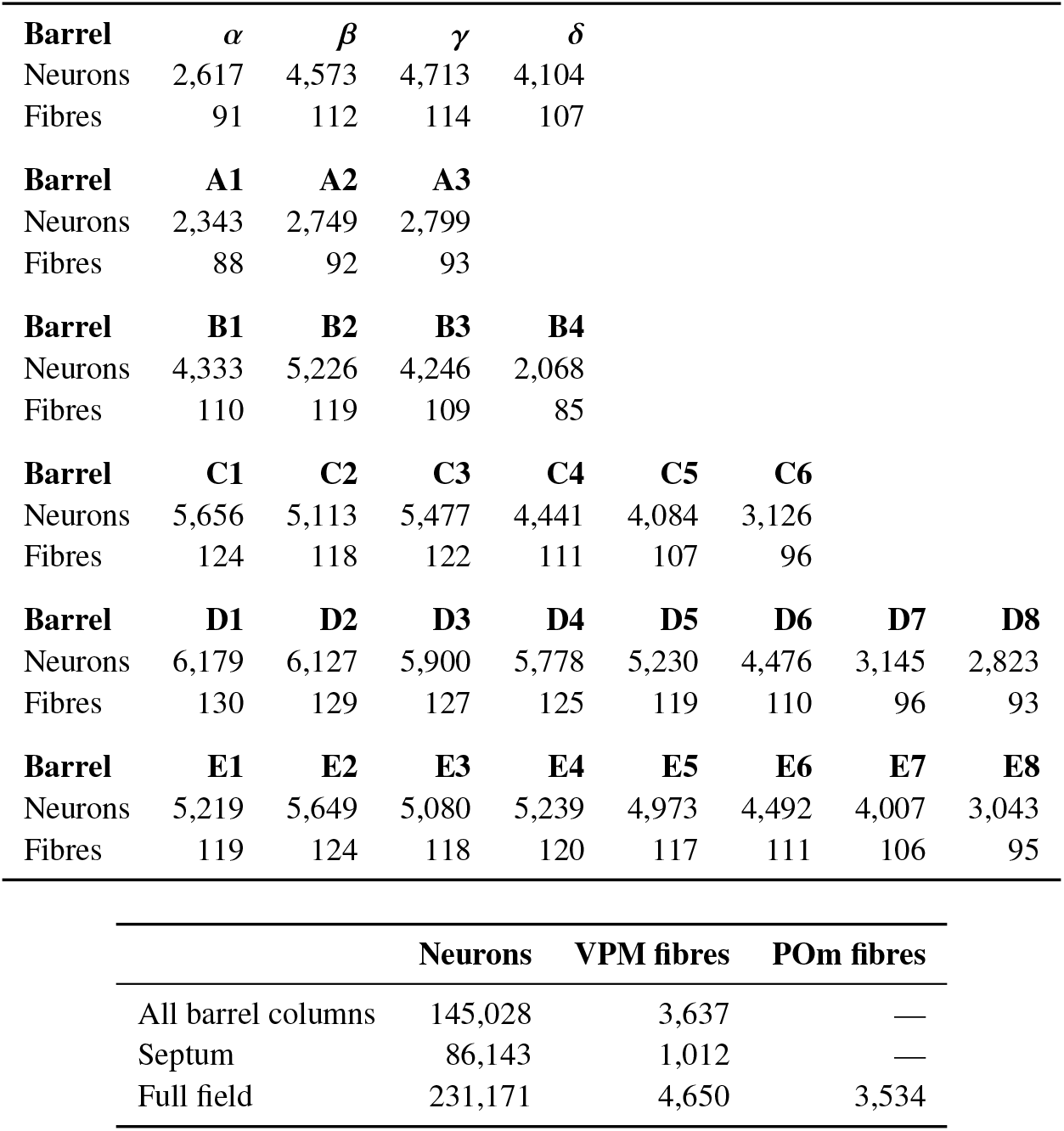
Neuron counts and VPM fibre counts per barrel column. Fibre counts per barreloid were obtained by applying the cortex-to-thalamus regression fitted to Meyer et al. (2013) (slope 0.011, intercept 61.2, *r*^2^
= 0.68) to the neuron count of each barrel column in the model. POm fibres carry no barrel-specific count and were scaled from the VPM total by the inter-nuclear ratio of 0.76.

**Table S2.** Reconstructions per morphological type in the curated release. Dendrite is the number contributing dendritic barcodes and parameter distributions to synthesis; axon is the number of axon entries available for grafting (some reconstructions appear more than once). Asterisks mark the four classes carrying the barrel constraint (Section *Boundary-guided dendritic growth*). Totals: 1,204 dendrite and 982 axon entries across 60 m-types.

| **Layer 1** | **L1_DAC** | **L1_HAC** | **L1_LAC** |  |  |
|---|---|---|---|---|---|
| Dendrite | 16 | 27 | 10 |  |  |
| Axon | 16 | 27 | 10 |  |  |
| **Layer 1** | **L1_NGC-DA** | **L1_NGC-SA** | **L1_SAC** |  |  |
| Dendrite | 19 | 19 | 14 |  |  |
| Axon | 19 | 19 | 14 |  |  |
| **Layer 2** | **L23_BP** | **L23_BTC** | **L23_CHC** | **L23_DBC** |  |
| Dendrite | 2 | 28 | 20 | 40 |  |
| Axon | 2 | 28 | 20 | 40 |  |
| **Layer 2** | **L23_LBC** | **L23_NBC** | **L23_NGC** | **L23_SBC** |  |
| Dendrite | 82 | 40 | 8 | 44 |  |
| Axon | 82 | 40 | 8 | 44 |  |
| **Layer 2** | **L2_IPC** | **L2_TPC:A** | **L2_TPC:B** |  |  |
| Dendrite | 4 | 6 | 38 |  |  |
| Axon | 2 | 4 | 27 |  |  |
| **Layer 3** | **L23_MC** | **L3_TPC:A** | **L3_TPC:C** |  |  |
| Dendrite | 42 | 49 | 11 |  |  |
| Axon | 40 | 19 | 5 |  |  |
| **Layer 4** | **L4_BP** | **L4_BTC<sup>\*</sup>** | **L4_CHC** | **L4_DBC** |  |
| Dendrite | 2 | 7 | 1 | 7 |  |
| Axon | 2 | 7 | 1 | 7 |  |
| **Layer 4** | **L4_LBC** | **L4_MC** | **L4_NBC** | **L4_NGC** |  |
| Dendrite | 31 | 17 | 25 | 1 |  |
| Axon | 31 | 17 | 25 | 1 |  |
| **Layer 4** | **L4_SBC<sup>\*</sup>** | **L4_SSC<sup>\*</sup>** | **L4_TPC** | **L4_UPC<sup>\*</sup>** |  |
| Dendrite | 15 | 21 | 31 | 36 |  |
| Axon | 15 | 8 | 21 | 21 |  |
| **Layer 5** | **L5_BP** | **L5_BTC** | **L5_CHC** | **L5_DBC** | **L5_LBC** |
| Dendrite | 3 | 11 | 3 | 9 | 21 |
| Axon | 3 | 11 | 3 | 9 | 21 |
| **Layer 5** | **L5_MC** | **L5_NBC** | **L5_NGC** | **L5_SBC** |  |
| Dendrite | 38 | 22 | 1 | 4 |  |
| Axon | 38 | 22 | 1 | 4 |  |
| **Layer 5** | **L5_TPC:A** | **L5_TPC:B** | **L5_TPC:C** | **L5_UPC** |  |
| Dendrite | 76 | 43 | 34 | 28 |  |
| Axon | 44 | 27 | 17 | 24 |  |
| **Layer 6** | **L6_BP** | **L6_BPC** | **L6_BTC** | **L6_CHC** | **L6_DBC** |
| Dendrite | 3 | 28 | 1 | 1 | 2 |
| Axon | 3 | 17 | 1 | 1 | 2 |
| **Layer 6** | **L6_HPC** | **L6_IPC** | **L6_LBC** | **L6_MC** | **L6_NBC** |
| Dendrite | 21 | 26 | 24 | 20 | 11 |
| Axon | 17 | 18 | 24 | 20 | 10 |
| **Layer 6** | **L6_NGC** | **L6_SBC** | **L6_TPC:A** | **L6_TPC:C** | **L6_UPC** |
| Dendrite | 1 | 5 | 20 | 18 | 17 |
| Axon | 1 | 5 | 6 | 6 | 5 |

**Table S3.** Mouse reconstructions common to the dendritic polarity analysis and the synthesis input set, by morphological type and source archive. Hundred of the 324 analysed mouse reconstructions also contributed to synthesis; m-types follow the synthesis nomenclature. Cells below origin from *NeuroMorpho* repositories.

| M-type | Archive | Cells |
|---|---|---|
| L23_DBC | Rudy | 1 |
| L23_DBC | Staiger | 8 |
| L23_MC | Staiger | 10 |
| L23_NBC | Rudy | 1 |
| L2_TPC:A | Petersen | 3 |
| L2_TPC:B | Peng | 4 |
| L2_TPC:B | Petersen | 6 |
| L3_TPC:A | Petersen | 1 |
| L3_TPC:C | Peng | 1 |
| L4_SBC | Peng | 3 |
| L4_SSC | Luebke | 15 |
| L4_SSC | Peng | 4 |
| L4_SSC | Staiger | 1 |
| L4_SSC | Tarusawa | 2 |
| L4_TPC | Peng | 2 |
| L4_UPC | Peng | 2 |
| L5_TPC:A | Peng | 16 |
| L5_TPC:B | Peng | 7 |
| L5_TPC:C | Peng | 4 |
| L5_UPC | Peng | 1 |
| Total |  | 92 |

**Table S4.** Postsynaptic potential amplitudes per pathway curated from the mouse barrel cortex literature. Amplitudes in mV, negative for inhibitory pathways; standard deviations are omitted where the source reports a single measurement. Entries with two references pool the studies listed.

| Reference | Pathway | Amplitude | SD |
|---|---|---|---|
| Lefort 2009, Jouhanneau 2015 | L2 EXC → L2 EXC | 0.535 | 0.065 |
| Lefort 2009 | L2 EXC → L3 EXC | 0.44 | 0.09 |
| Lefort 2009 | L2 EXC → L4 EXC | 0.31 |  |
| Lefort 2009 | L2 EXC → L5 EXC | 0.448 | 0.061 |
| Lefort 2009 | L3 EXC → L2 EXC | 0.71 | 0.15 |
| Lefort 2009 | L3 EXC → L3 EXC | 0.78 | 0.07 |
| Lefort 2009 | L3 EXC → L4 EXC | 0.36 | 0.09 |
| Lefort 2009 | L3 EXC → L5 EXC | 0.994 | 0.173 |
| Lefort 2009 | L4 EXC → L2 EXC | 0.98 | 0.24 |
| Lefort 2009 | L4 EXC → L3 EXC | 0.58 | 0.13 |
| Lefort 2009, Scala 2015 | L4 EXC → L4 EXC | 0.97 | 0.247 |
| Lefort 2009 | L4 EXC → L5 EXC | 0.627 | 0.105 |
| Lefort 2009 | L4 EXC → L6 EXC | 2.27 | 1.72 |
| Lefort 2009 | L5 EXC → L2 EXC | 0.489 | 0.087 |
| Lefort 2009 | L5 EXC → L3 EXC | 0.424 | 0.044 |
| Lefort 2009 | L5 EXC → L4 EXC | 0.377 | 0.000 |
| Lefort 2009 | L5 EXC → L6 EXC | 0.403 | 0.096 |
| Lefort 2009 | L6 EXC → L5 EXC | 0.227 | 0.000 |
| Lefort 2009 | L6 EXC → L6 EXC | 0.53 | 0.19 |
| Avermann 2012, Jouhanneau 2018 | L2/3 EXC → L2/3 EXC | 0.34 | 0.086 |
| Jouhanneau 2018, Pala 2015 | L2/3 EXC → L2/3 Pvalb | 0.82 | 0.623 |
| Jouhanneau 2018, Pala 2015 | L2/3 EXC → L2/3 Sst | 0.54 | 0.357 |
| Jouhanneau 2018 | L2/3 Pvalb → L2/3 EXC | −0.33 | 0.042 |
| Avermann 2012 | L2/3 Pvalb → L2/3 Pvalb | −0.56 | 0.14 |
| Avermann 2012 | L2/3 Pvalb → L2/3 Vip | −0.83 | 0.25 |
| Scala 2015 | L4 EXC → L4 BTC | 0.53 | 0.16 |
| Scala 2015 | L4 BTC → L4 EXC | 0.38 | 0.12 |
| Scala 2015 | L4 BTC → L4 BTC | 0.19 | 0.05 |

